# Diversified global vegetable oil production can mitigate climate change and achieve socio-environmental co-benefits

**DOI:** 10.1101/2024.07.16.603827

**Authors:** Siyan Zeng, Fu Chen, Chengcheng Zhang, Shenggen Fan, Baojing Gu, Erik Meijaard, Kyle Frankel Davis, Thomas Cherico Wanger

## Abstract

Global vegetable oil production is a crucial ingredient for food and household products but also a major source of greenhouse gas (GHG) emissions and other environmental externalities. A diversified supply may improve the sustainability and resilience of profitable vegetable oil production globally. Here, we provide the first spatial assessment of the environmental and economic effects of the globally dominant vegetable oil crops soybean, oil palm, rapeseed, sunflower, groundnut, and olive globally, based on new data from a meta-analysis, life cycle assessment, and process-based crop growth modelling. We show that the total annual GHG emissions, water use, land use, pesticide use, and farmer income of the six vegetable oil crops are 1.2 Gt CO_2_eq/year, 876.0 km^3^/year, 168.3 Mha/year, 953.6 kt/year, and 66.5 billion USD/year, respectively. We also found that diversified vegetable oil production, for instance, through increased fractions of a highly productive oil tree crop *Camellia oleifera* can reduce GHG emissions, water use, land use, and pesticide use by 14.0%, 4.9%, 7.3%, 9.3%, respectively and increase farmer income by 82.6%. When considering enabling conditions for technology and policy support in a technology-policy-enabling scenario, these benefits increase 1.3, 2.1, 1.8, 1.7, and 3.2 folds, respectively. Our results provide practical guidance on where and how diversified oil crop production can advance a sustainable food system transformation and help to mitigate climate change.

---

The global food system faces the dual challenge of meeting the demands for food, fiber, and energy from a growing population while addressing exacerbating environmental issues (1). More than 30% of the total agricultural land is currently used for vegetable oil crop production, 80% of which is occupied by the six major oil crops soybean (*Glycine max*), oil palm (*Elaeis guinensis)*, rapeseed (*Brassica napus*), groundnut (*Arachis hypogaea*), sunflower (*Helianthus annuus*), and olive (*Olea europaea*) (2). These oils are used in a variety of products including cooking oil, cosmetics, and biofuel production - with steadily increasing global demand (3). However, production supplies are strongly impacted by geopolitical tensions and climate change, both of which affect global trade and prices (4). Moreover, vegetable oil production is a major driver of cropland expansion, associated with up to 1.9 GtCO_2_ - eq (23%) of global food carbon emissions (5) and heavily relying on pesticides and fertilizers, which exert pressure on soil and water resources (6,7). Palm oil and soybean oil, two of the most widely-used oil crops for daily commodities, have been linked with biodiversity decline, deforestation, and regionally variable effects on human livelihoods (6,8,9). Geographically, these impacts are most prevalent in the Southeast Asian tropics for palm oil and South America for soybean oil (10,11,12). Previous studies have quantified aspects of these effects across spatial scales, showing that there is a large potential to diversify global vegetable oil production (13) and optimize crop distribution to better meet future demand (14). Despite decades of research efforts, a spatially explicit synthesis of environmental and economic effects of the major vegetable oil crops globally is currently missing (15,16). Furthermore, an analysis of where and how diversified oil crop production could mitigate climate change impacts and achieve socio-environmental co-benefits is lacking.

Efforts to mitigate environmental impacts while maintaining or even increasing vegetable oil yields have focused on improving supply chain transparency through effective policy (e.g. the EU deforestation free supply chain law including oil palm and soybean; 17,18), enhancing farmer income through sustainability standards (19), and various agricultural diversification practices (e.g., crop rotations or agroforestry; 20). Diversified supply of vegetable oil ideally contributes to a triple win-win-win solution whereby the outcomes are healthier oils with a lower environmental footprint and higher revenue (21). Identifying such new opportunities usually requires breeding or gene editing existing crops for preferable traits (22), improved management for instance through agroforestry (23,24), exploring alternative exotic oils such as microbial single-cell oils (25,26) or bioprospecting new species (27). Woody oil crops are potential candidates that may be able to combine high yields with high nutritional value and high environmental benefits. *Camellia oleifera* (hereafter Camellia) is currently cultivated almost exclusively in Asia, including China, Japan, and Vietnam (28) but it may provide such a triple win solution (23,28). Camellia yield potential of 2.8 t/ha is the second highest just after oil palm (29,30), the oil is of high nutritional value on monounsaturated fatty acid, squalene, and vitamin E (see Table S1) (28,31,32), and the tree crop has been shown to protect farmer livelihoods, prevent soil erosion, and increase biodiversity through agroforestry or intercropping (27,33,34,35). As a generalist species, Camellia also has low habitat requirements (28,33) and, hence, allows for cultivation on degraded land and in regions, where arable land is scarce. However, limited accessible data on Camellia’s environmental and social parameters (36) has prevented an integrated assessment of the potential environmental and social effects of its inclusion in diversified vegetable oil production mixes (6,15).

Here, we first provide a quantitative global and spatially explicit assessment of GHG emissions, water use, land use, fertilizer use, and farmer income for six major vegetable oil crops. We do this based on a meta-analysis, life cycle assessments, and process-based crop growth modelling with spatially interpolated yields for six major oil crops soybean, oil palm, rapeseed, sunflower, groundnut, and olive. We then identify socio-environmental co-benefits between current oil crop distributions and a potential replacement oil crop, for instance Camellia that can be used without yield loss but with increased farmer income (see Fig. S1). Our results enable decision makers to optimize global vegetable oil crop distribution for the least environmental impact but the greatest benefits without land expansion and oil yield loss based on new spatially explicit data. We provide a novel approach to evaluate opportunities for partial crop substitution achieving socio-environmental co-benefits.

## Results

### Environmental impact of vegetable oil crops

We conducted a global meta-analysis (16) based on 837 unique records in 285 studies to parameterize a life cycle assessment and then mapped the environmental impacts of groundnut, soybean, oil palm, sunflower, rapeseed, and olive (for details see Methods Section and Extended Section 1.2.3). In 2015, all vegetable oil crops in our study accounted for 298.3 Mha of harvested areas, equivalent to 21.4% of global total harvested areas, of which 40.5% was occupied by soybean, 11.6% by rapeseed, 8.9% by groundnut, 8.2% by sunflower, 7.6% by oil palm, and 3.4% by olive (2).

We estimate that the total annual GHG emissions of the six vegetable oil crops are 1.2 Gt CO_2_eq/year, with 44.5%, 34.9%, 8.9%, 6.1%, 4.3%, and 1.3% coming from soybean, oil palm, groundnut, rapeseed, sunflower, and olive oil, respectively (Fig. 1a and Fig. S14). The total global emissions from soybean are 547.1 Mt CO_2_eq/year, mainly originating in South America (466.4 Mt CO_2_eq/year) and North America (45.5 Mt CO_2_eq/year). At a country level, soybean production in Brazil generated the highest GHG emissions of 59.2%, followed by Argentina, the Republic of Paraguay, and the United States. For oil palm, the total amount is 429.2 Mt CO_2_eq/year and mainly emitted in Asia and Oceania (360.2 Mt CO_2_eq/year) and Africa (48.5 Mt CO_2_eq/year). The highest total annual GHG emissions of palm oil at country level come from Indonesia (64.4%), the world’s largest palm oil producer, followed by Malaysia (17.9%), and Nigeria (9.2%). These emissions likely come from N_2_O, and CO_2_ through fertilizer application and land use change from forest biomass or burning organic soil (see Methods). The GHG emission intensity (i.e., amount of GHG emitted per 1 liter oil) is highest in palm oil (9.7 kg CO_2_ eq/L) due mainly to deforestation, followed by olive oil, groundnut oil, soybean oil, rapeseed oil, and sunflower oil (Fig. S16). For more details on all oil crops see Fig. S15 and Fig. S33.

**Fig. 1:**
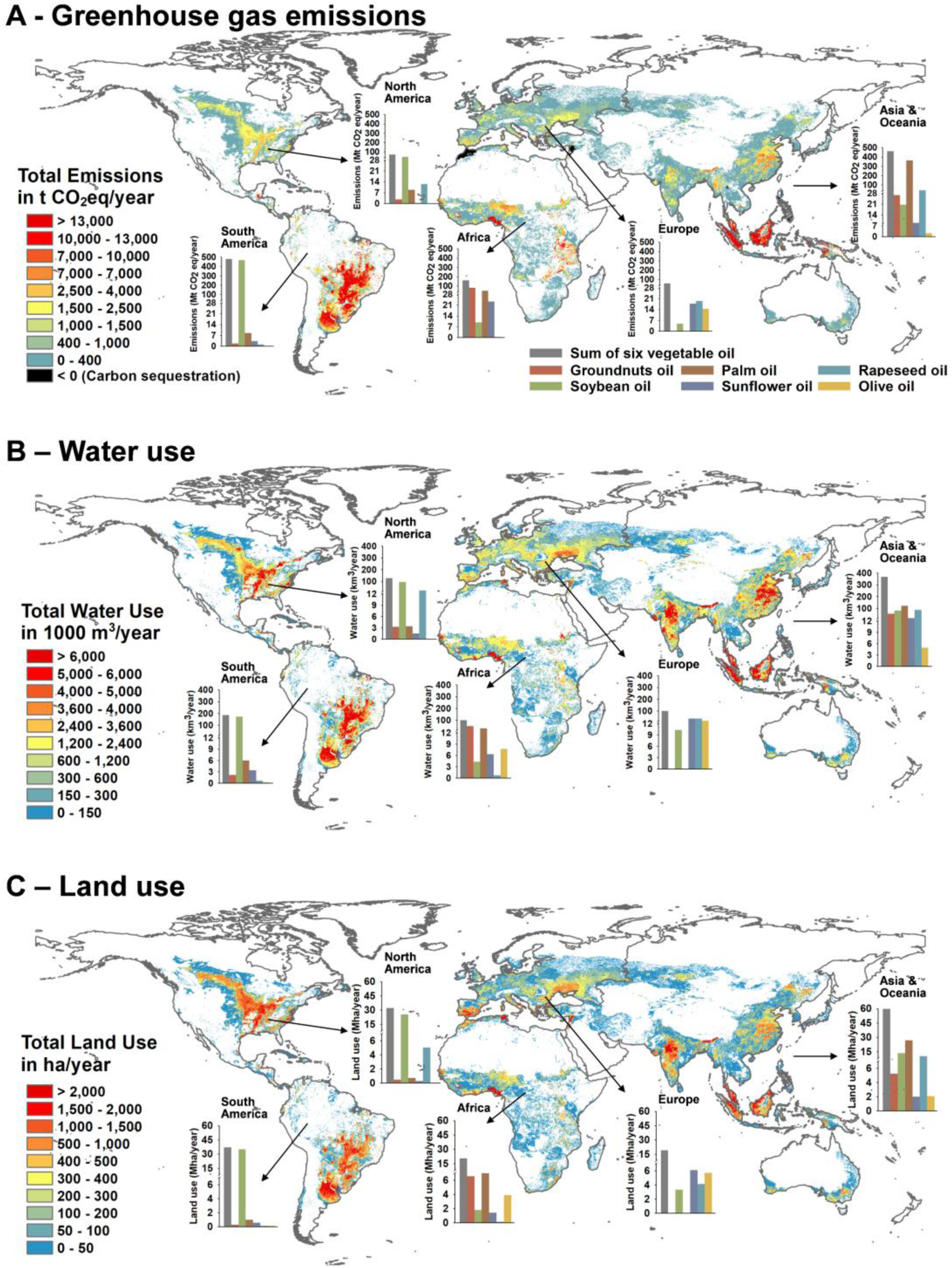
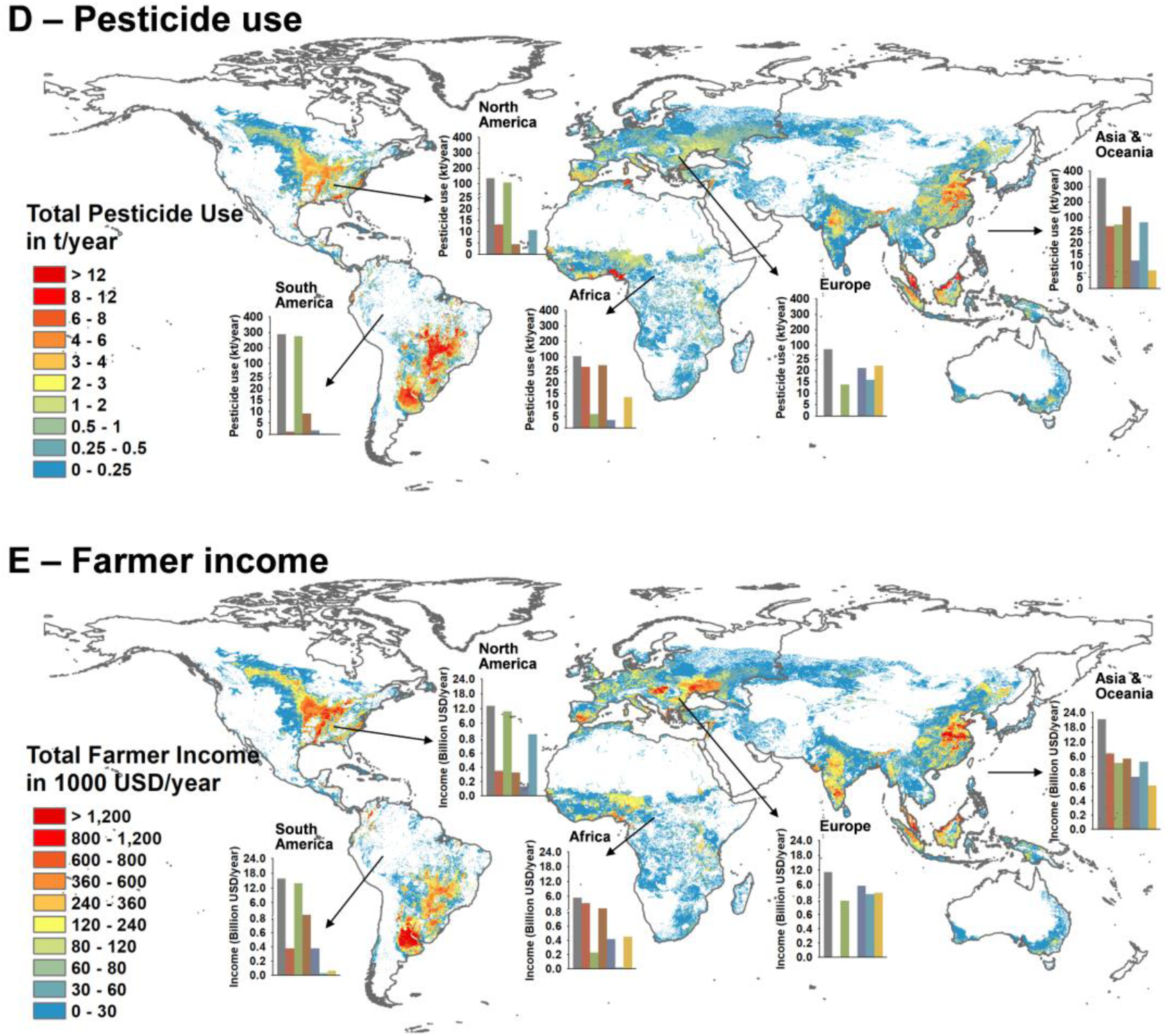
Global environmental and socioeconomic effect of the six main vegetable oil crops. It shows the effects of six vegetable oil crops - groundnut, soybean, oil palm, rapeseed, sunflower, and olive - on: GHG emissions (A), water use (B), land use (C), pesticide use (D), and farmer income (E).

The total annual water use (i.e., the aggregation of green, blue, and grey water use; see Methods for details) is 876.0 km^3^/year, with the descending order of soybean (364.2 km^3^/year), followed by oil palm (160.8 km^3^/year), rapeseed (145.7 km^3^/year), groundnut (109.9 km^3^/year), sunflower (65.2 km^3^/year), and olive (30.1 km^3^/year) (Fig. S15b and Fig. S14). Water use in Asia and Oceania is highest at 360.0 km^3^/year, followed by South America, North America, Africa, and Europe. The water-use intensity for producing 1-liter oil also varied across regions (Fig. S19), which is affected by the length of the crop growing period (especially for tree crops like oil palm and olive), evapotranspiration, crop-specific factors, and climate variables. The mean water-use intensity was highest for olive oil (14.5 m^3^/L) and soybean oil (13.9 m^3^/L) followed by groundnut oil, rapeseed oil, sunflower oil, and palm oil (Fig. S19).

The total annual land use of all six vegetable oil crops is 168.3 Mha/year when taking multi-cropping, inter-cropping, or fallow into account (see Methods). Total land use is highest for soybean with 79.1 Mha/year) and lowest for sunflower with 10.2 Mha/year (Fig. S14). Regionally, land use is highest in Asia and Oceania (59.7 Mha/year) and South America (36.9 Mha/year) followed by North America, Africa, and Europe (Fig. 1c). The regional variation in land-use intensity per liter oil was low in rapeseed and sunflower but relatively high for palm, groundnut, soybean and olive (Fig. S20). The mean value of land-use intensity differs for producing 1-liter vegetable oil among crops, ranging from 8.4 m^2^ for sunflower to 51.6 m^2^ for olive, respectively.

The total annual pesticide use of the six vegetable oil crops is 953.6 kt/year, with the majority being used in Asia and Oceania (37.1%) and South America (30.2%) (Fig. 1d). Pesticide use in South America from soybean production is 274.8 kt/year, which accounts for 60.9% of total pesticide use for soybean production globally and is higher than the total amount used for all other vegetable oil crops (Fig. S21).

We found the total farmer income from the six vegetable oil crops to be 66.5 billion USD/year, with the majority being earned in Asia and Oceania (31.9%) and South America (23.7%) (Fig. 1e). The farmer income differs between regions and is highest in Asia and Oceania for groundnut (7.3 billion USD/year), palm (5.2 billion USD/year), and rapeseed (3.8 billion USD/year), followed by South America for soybean (13.9 billion USD/year) and then Europe for both sunflower (5.5 billion USD/year) and olive (2.6 billion USD/year). For all details see Fig. S22.

### Crop replacement to mediate socioenvironmental effects of oil production

We use *Camellia oleifera* as a potential replacement oil crop case study that can reduce environmental impacts and enhance socioeconomic benefits of global vegetable oil production. Through land suitability mapping (Methods), we find that 52.2 Mha within the current vegetable oil production areas are suitable for Camellia production without land expansion for vegetable oil crop cultivation (Fig. S11b and Table S16). Brazil is the major soybean oil producer and could replace almost one quarter (24.4%) of its vegetable oil crop planting areas with Camellia. With a cumulative potential of over 50%, Argentina, the United States, Indonesia, and India could replace 19.8%, 11.1%, 8.4%, and 5.4% within the current vegetable oil production areas, respectively. China is the biggest producer and consumer of Camellia oil globally with the potential of 4.9% of Camellia planting in the current vegetable oil crop planting areas (Fig. S12).

Using yield modelling for Camellia cultivation, we predict a mean potential yield of 14.4 t/ha and the highest yield to be 32.0 t/ha for fresh fruit (Fig. S13d), which can be used to produce 2.1 t/ha and 4.6 t/ha Camellia oil (Fig. S5). Advanced biotechnology and management techniques such as genome editing for seed oil domestication and sprout anvil grafting technology could increase Camellia oil yields up to 22.5 t/ha with a further 7-10 times increase in the future (27,28,29). We therefore use three spatially explicit yield scenarios to simulate the environmental impacts of Camellia oil in a life cycle assessment: a conservative scenario (or 50% of potential yield, S1), strategic improvement scenario (or 75% of potential yield, S2), and a technology-policy-enabling scenario (or 100% of potential yield, S3) (Methods).

Camellia oil produces substantially lower GHG emission than most current oil crops, with 4.4 kgCO_2_ eq/L, 3.0 kgCO_2_ eq/L, and 2.3 kgCO_2_ eq/L in S1, S2 and S3, respectively. compared to the six main vegetable oil crops (3.5 kgCO_2_ eq/L for sunflower oil; 4.0 kgCO_2_ eq/L for rapeseed oil; 8.4 kgCO_2_ eq/L for groundnut oil and soybean oil, 8.5 kgCO_2_ eq/L for olive oil; 9.7 kgCO_2_ eq/L for palm oil; Fig. S16). Specially, Camellia oil is still advantageous in the yield scenario S2 and S3 above, if high GHG emission from land use change (the above and below-ground C stock change, forest burning, and organic soil burning) for soybean oil and palm oil are excluded (4.4 kgCO_2_ eq/L; 2.7 kgCO_2_ eq/L) (Fig. S17 and Fig. S18). Camellia oil also requires much less water (9.7 m^3^/L) starting from Scenario 2 than soybean oil (13.9 m^3^/L) and olive oil (14.5 m^3^/L) and compared to all vegetable oil crops in Scenario 3 (Fig. S19 and Fig. S24b). Land use of Camellia oil for all three scenarios (6.3, 4.2, and 3.1 m^2^/L for S1, S2, and S3, respectively; Fig. S25) is much lower than all other vegetable oils (7.9 m^2^/L, 8.4 m^2^/L, 9.2 m^2^/L, 10.4 m^2^/L, and 18.6 m^2^/L for groundnut, sunflower, soybean, rapeseed, and olive oil, respectively; Fig. S20) except for palm oil (2.9 m^2^/L).

### Camellia crop replacement without land expansion and yield loss

For tangible solutions, we consider only solutions without land expansion and where overall vegetable oil yields (i.e., the total yield produced by any combination of vegetable oil planted) per area are maintained. We firstly focused on replacing global vegetable oil crop production area with Camellia by simultaneously optimizing all five objectives across all six vegetable oil crops (i.e., reducing GHG emissions, water use, land use, and pesticide use per liter oil, and increasing farmer income per hectare land). Optimization for all five objectives leads to the lowest area replacement with 26.3 Mha (11.2%), 37.1 Mha (15.8%), and 42.0 Mha (17.9%), corresponding to Camellia yield scenarios 1 to 3, respectively (Fig. 3). Regarding spatial patterns of replacement, the main benefits for Camellia replacement in Africa are for groundnut, oil palm, and sunflower, in South America for soybean, in Asia and Oceania for rapeseed, and in Europe for olive (Fig. 2). Southern Brazil, Argentina, Paraguay, southern Nigeria, and Tanzania, central India, and southern China were among the top-ranked 10% areas for reducing global GHG emission (Fig. 3D; Fig. S30a, S31a, and S32a), and for optimizing all five objectives together (Fig. S30e, S31e, and S32e). The eastern United States, southern China, Brazil, central India, and Guinea had the highest potential to reduce water use (Fig. S30b, S31b, and S32b), land use (Fig. S30c, S31c, and S32c), pesticide use (Fig. S30d, S31d, and S32d), and to increase farmer income (Fig. S30e, S31e, and S32e). For more detailed results on replacing global vegetable oil crop production area with Camellia by individually optimizing specific objective across all six vegetable oil crops see Table S22 & Fig. S26.

**Fig. 2:**
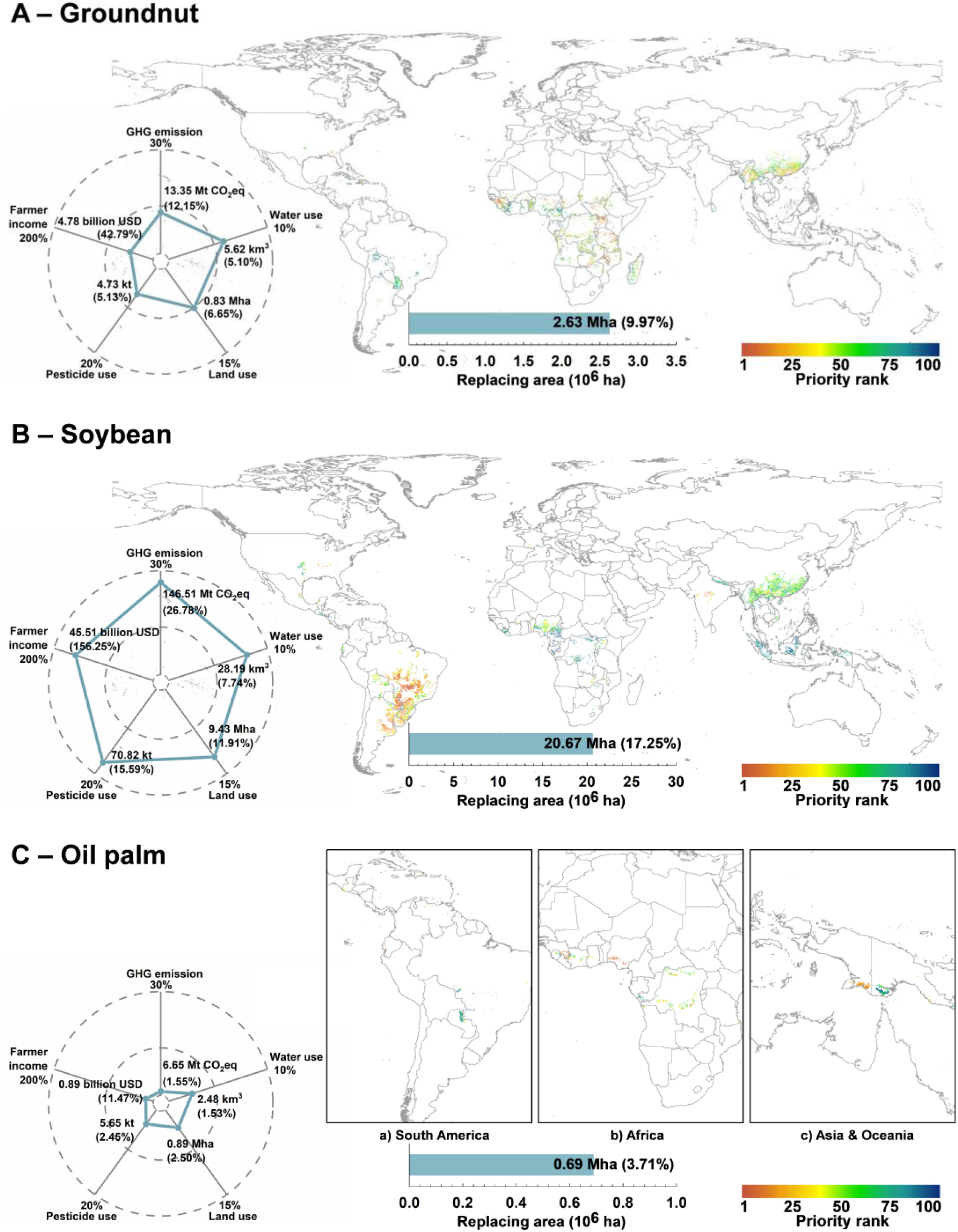
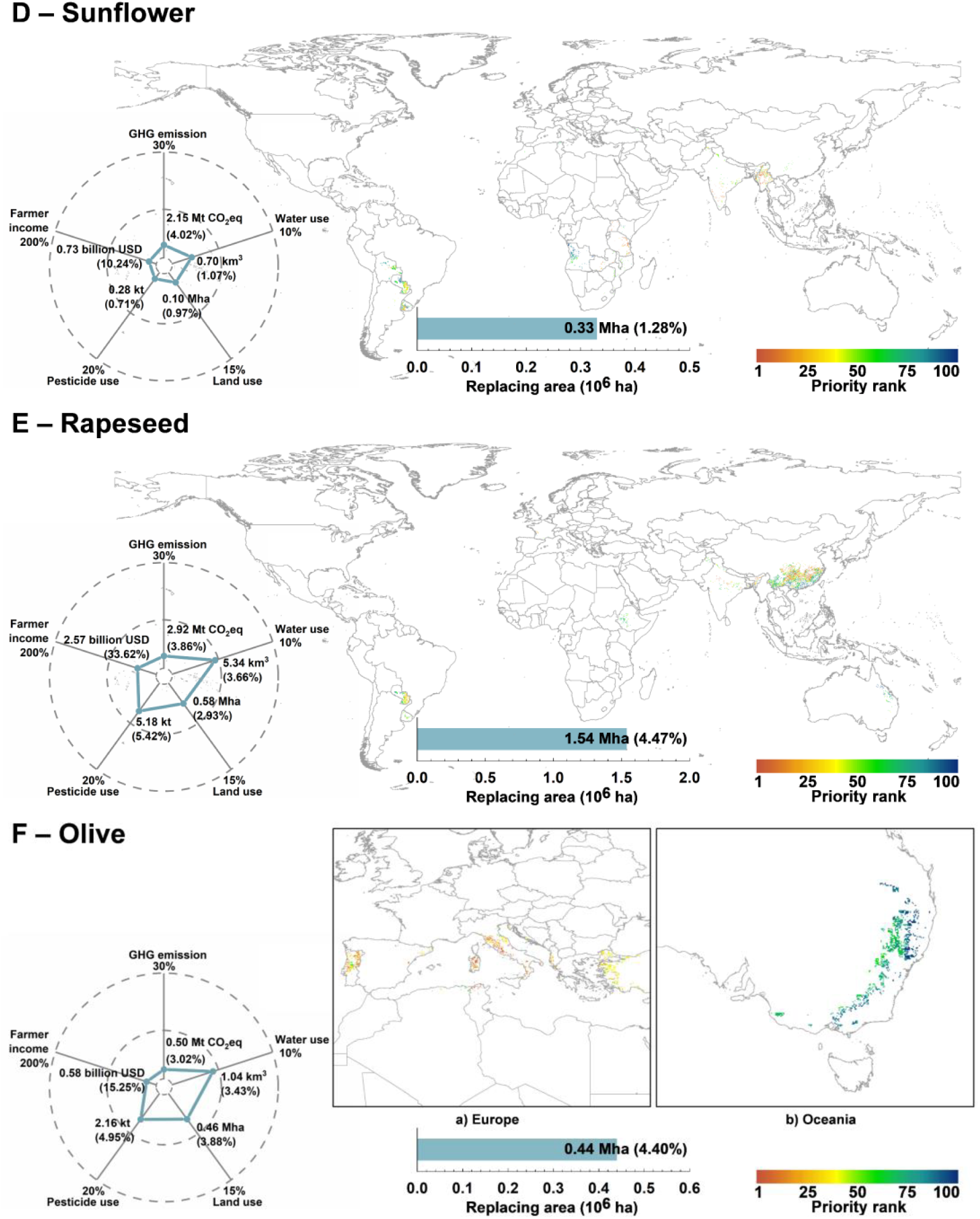
Priority areas for crop replacement with Camellia cultivation. Panels show the replacement areas for six vegetable oil crops under a conservative Camellia yield scenario (S1) and optimizing reduction of GHG emissions, water use, land use, pesticide use per liter oil, and increase of farmer income per hectare land simultaneously: groundnut (A), soybean (B), oil palm (C), sunflower (D), rapeseed (E), and olive (F). Note that for palm (C) and olive (F), we zoom in to continent level as replacing areas would be difficult to see otherwise. Priorities refer to the highest (red: 1–10) to lowest (blue: 90–100) valuable areas for Camellia replacement to achieve simultaneous objective optimization.

**Fig. 3:**
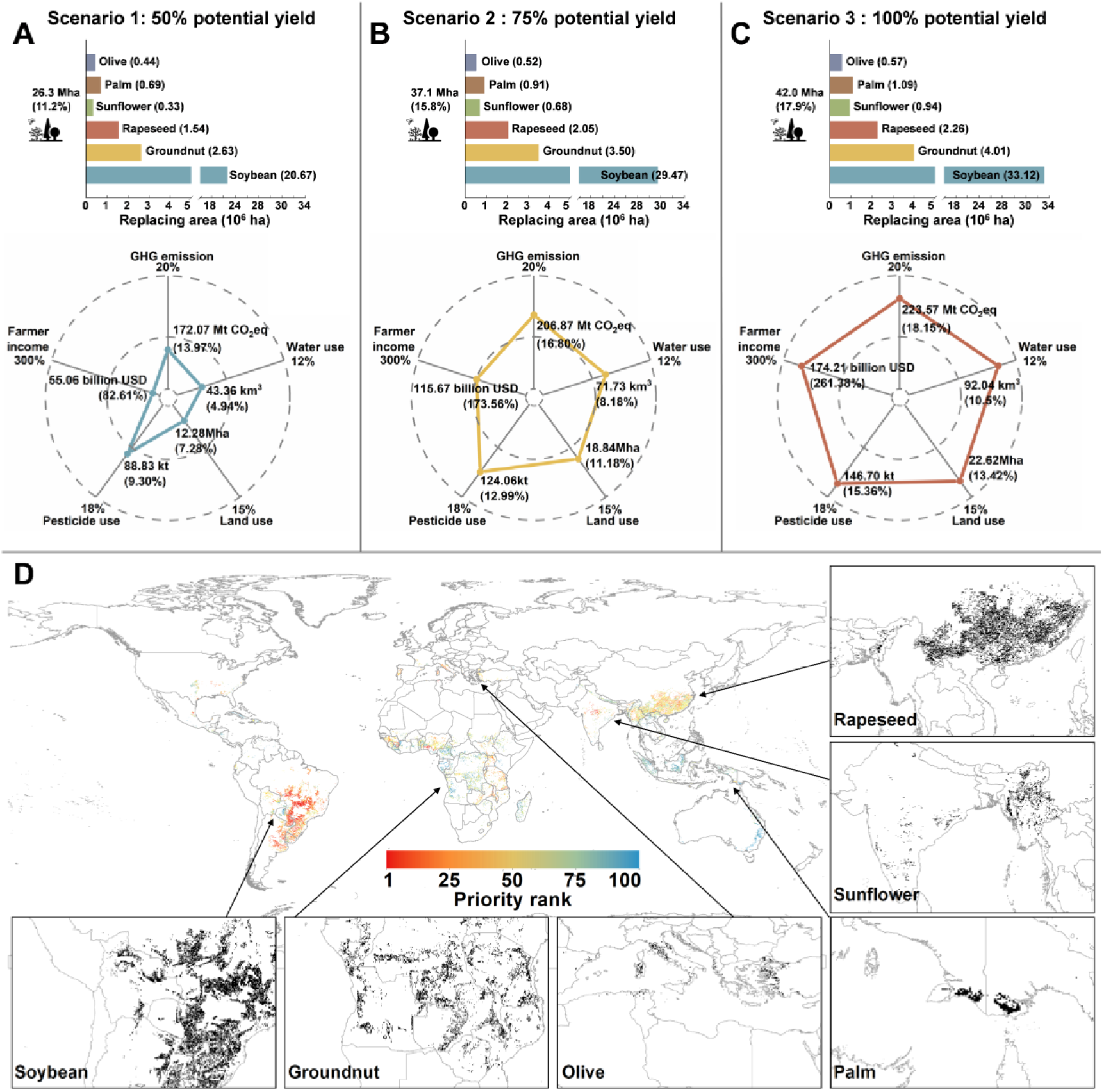
Global spatial priority areas for Camellia replacement to optimize all five socioenvironmental co-benefit objectives simultaneously. For a conservative (A), strategic (B), and technology-policy-enabling Camellia yield improvement scenario (C) when all five objectives are optimized simultaneously with reducing GHG emission, water use, land use, and pesticide use per liter oil, and increasing farmer income per hectare land. The histograms and radar plots show the total replacing areas for each vegetable oil crop and quantified benefits for each objective, respectively. Spatial benefits of vegetable oil crop replacement differ between continents and crops as shown for the conservative Camellia yield scenario (S1) (D). The black color shows the Camellia replacing pattern for each vegetable oil crop. For details see main text.

### Socio-environmental co-benefits of global diversified vegetable oil production

After showing how prioritization modelling allows spatially explicit crop replacement with Camellia as a case study, we quantify the environmental and socioeconomic effects of such an exemplary crop replacement for diversified global vegetable oil production. For simultaneously optimizing all five objectives across all six vegetable oil crops in the conservative Camellia yield scenario 1, we found that replacing 26.3 Mha (11.2%) of global vegetable oil production with Camellia resulted in reduction of GHG emissions, water, land, and pesticide use by 14.0% (172.1 Mt CO_2_ eq/year), 4.9% (43.4 km^3^/year), 7.3% (12.3 Mha/year), 9.3% (88.8 kt/year), respectively, and an increase of farmer income by 82.6% (55.1 billion USD/year) (Fig. 3A). Compared to the conservative yield scenario 1, Camellia yield improvement scenario 2 and the ambitious technology and policy enabling yield scenario 3 reduced GHG emissions 1.2 and 1.3 folds, water use 1.7 and 2.1 folds, land use 1.5 and 1.8 folds, pesticide use 1.4 and 1.7 folds, and increased farmer income 2.1 and 3.2 folds, respectively (for details see Fig. S26, Fig. 3A, Fig. 3B, and Fig. 3C). For single-objective results under the three Camellia yield scenarios, refer to Fig. S27, S28, and S29.

For simultaneously optimizing all five objectives and individual vegetable oil crops in the conservative Camellia yield scenario 1, the replacing area for these six oils production areas are as follows: soybean at 20.7 Mha (17.3%), groundnut at 2.6 Mha (10.0%), rapeseed at 1.5 Mha (4.5%), oil palm at 0.7 Mha (3.7%), olive at 0.4 Mha (4.4%), and sunflower at 0.3 Mha (1.3%) (Fig. 2), which are accounting for 8.8%, 1.1%, 0.7%, 0.3%, 0.2%, and 0.1% of all vegetable oil crops harvested areas. The soybean area replacement then results in the highest benefits of all five objectives, with 26.8% (146.5 Mt CO_2_eq), 7.7% (28.2 km^3^), 11.9% (9.4 Mha), 15.6% (70.8 kt), and 156.3% (45.5 billion USD) of decreasing GHG emission, water use, land use, pesticide use, and increasing farmer income, respectively. Replacing olive production areas have the lowest effect on decreasing GHG emissions (0.5 Mt CO_2_eq) and increasing farmer income (0.6 billion USD), while replacing sunflower production areas has the lowest effect on water use (0.7 km^3^), land use (0.1 Mha), and pesticide use (0.3 kt).

## Discussion

Our spatially explicit results on the environmental and economic impacts of global vegetable oil production allow for targeted spatial planning and close critically important knowledge gaps (15). The total GHG emissions of six main vegetable oil crops, for soybean and for oil palm are 1.2 Gt CO_2_eq/year, 0.5 Gt CO_2_eq/year, and 0.4 Gt CO_2_eq/year, respectively, which corresponds well with non-spatial values of 1.9 Gt CO_2_eq/year, 0.6 Gt CO_2_eq/year, and 0.9 Gt CO_2_eq/year, respectively (5). There is some variation in the total water uses for each oil crop, because other studies only considered the green and blue water use (37). However, irrigation and fertilization in traditional vegetable oil crops production results in a significant grey water footprint (38), and, hence, must be considered in the evaluations.

Our result also highlights the importance of using spatially explicit analysis to identify differences in regional patterns. The total land use of six vegetable oil crops is 168.3 Mha and, hence, lower compared to the harvested area of 222.1 Mha, because we consider spatially explicit land occupation by multiple cropping, intercropping, or fallow (see Methods) (2). We found regional variation in land-use intensity to be low in rapeseed and sunflower but high for the other four vegetable oil crops (Fig. S20). For instance, producing 1 liter oil requires up to 233.3 m^2^/year of land in Portugal for olive oil and 135.1 m^2^/year in South Africa for soybean oil (39,40). Our systematic literature review and meta-analysis covered 10,500 pieces of regional or farm-level input information in 43 countries (Fig. S3) and considered 108 scenarios of replacement prioritization under potential yield developments when assessing the impact of Camellia cultivation within all suitable areas. Therefore, our work provides a robust systematic mapping of vegetable oil crops’ environmental impacts and exploration of global priority areas for long-term potential crop substitution to date. More specially, with the increasing application of life cycle assessment on the spatial scale (41,42), our research emphasizes the importance of understanding the detailed spatial pattern of crop production and the carbon–land–water nexus to identify potential cultivation improvement strategies, land use planning, or crop site-specific redistribution/substitution. Due to the differences in local climatic, topographic, soil, and farming practices, the agricultural inputs, such as fertilizer and pesticides vary (43,44), leading to the corresponding crop yields, farmer income, and environmental output varying greatly in different regions. These variations would be obscured if a regional or national level life cycle inventory were used, resulting in missed opportunities for strategic crop substitution in priority areas.

Substitution of vegetable oil crop production with Camellia may not only supply cooking oil, feed, biofuel, and cosmetics production, but also bring additional opportunities for farmer livelihoods, nutrition security and biodiversity conservation. First, Camellia production will likely contribute to livelihood security, because Camellia replacement leads to vegetable oil production increase in all our scenarios. In China, (www.ce.cn) (2024) the annual income generated from the Camellia oil supply chain exceeded 8.3 billion USD in 2023 and promoted rural development by increasing employment by 24.0% (4 million people) in Jiangxi Province (45). Farmer income is diversified through Camellia cultivation and, hence, may be less likely to be affected by external shocks from climate change depending on specific crop combinations. Second, Camellia oil is also of high nutritional value and may contribute to nutrition security if supplies increase. For instance, Camellia oil has the highest monounsaturated fatty acid composition compared to all other vegetable oils (Fig. 4a) and is rich in antioxidants. The oil has also alleged medicinal benefits such as regulating blood lipids, alleviating liver damage, and preventing Alzheimer disease (46) (Fig. 4b). Lastly, several studies have shown biodiversity benefits related to Camellia cultivation in agricultural landscapes through promoting soil aggregate formation and improving soil fertility (27,34,35). Our results suggest shifting from annual soybean, groundnut, or rapeseed systems to a perennial production system. Such a shift has been shown to reduce soil erosion, nutrient depletion and high pesticide input requirements (15,16). Moreover, Camellia production is suitable for agroforestry production, which by itself is known to enhance agrobiodiversity providing essential ecosystem services (47,48) and has been reported with soybean, groundnut, and rapeseed (49). Camellia agroforestry with soybean, corn and Purple Sweet Potato in China have been shown to regulate plantation air and soil temperature, and soil relative humidity, which in turn can enhance shoot growth of Camellia and understory plant diversity (50,51). In China, changes in the national policy to promote Camellia cultivation has facilitated restoration of degraded lands and promoting the ‘ecological civilization’ concept (52,53,54). Future research will likely expand quantification of the above aspects – and inevitably uncover local tradeoffs – that we can only discuss here based on singular case studies.

**Fig. 4:**
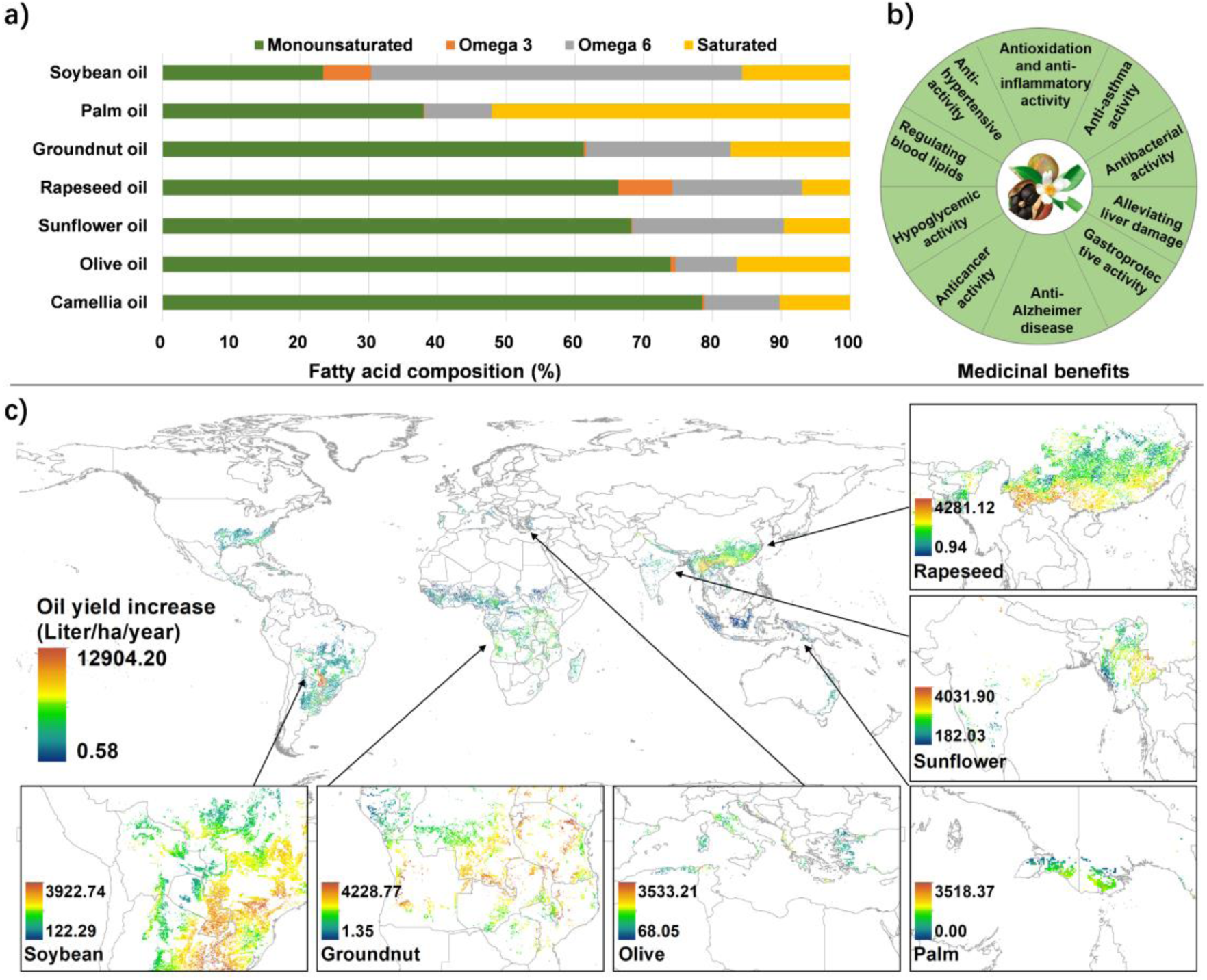
Spatial pattern of potential maximum yield increase from Camellia diversified global vegetable oil production. The nutritional value (a) and medicinal benefits (b) of Camellia oil. The oil yield improvement opportunity under a technology-policy-enabling scenario (S3; 100% potential yield) to replace local vegetable oil crops according to the comprehensive consideration of all objectives (c).

Benefits of a diversified global vegetable oil production are promising, but implementation requires careful local evaluation and facilitation of an enabling environment. First, infrastructure and financial subsidies are required to support Camellia cultivation with high labor input and profitability during the first 5 years of planting (55). For instance, large soybean estates only need 0.005 worker/ha in Brazil (15) but 0.247 worker/ha in Kenya (56), when Camellia production needs 0.385 worker/ha (Table S5; Fig. S34). At present, however, the price for Camellia oil is 8.9 ∼ 17.7 times higher than soybean oil (Fig. S22), which can somewhat compensate for higher labor costs in different regions of the world. In the future, planting and harvesting technology will advance and labor requirements for Camellia production will decrease, as it has for instance been the case in cotton (57,58). Suitable subsidy pathways may follow government schemes as seen for instance for cocoa agroforestry production in Ghana, West Africa or impact investments into sustainable agricultural landscapes (59,60). Second, lock-ins from cultural values, culinary preferences, palatability, perceived health benefits, and social norms (61,62) are of critical consideration. For instance, olive oil production in Italy is a cultural heritage and likely much more difficult to substitute compared to soy production in Brazil, which is driven by large corporations rather than culture (12,63). Lastly, technology innovations for instance in breeding and farming practices will close Camellia yield gaps and approaching yields similar to palm oil, making crop substitutions more attractive to all stakeholders (Fig. 4c; 29,30) with sizable socioenvironmental benefits (64).

Overall, diversified vegetable oil production provides an important perspective on how multi stakeholder collaboration can advance the Global Food systems transformation. At the consumer level, overconsumption of vegetable oil is an important issue around the world, with significant health concerns. In China, the per capita consumption of vegetable oil is double of the WHO recommended levels (65). Changing consumer behavior towards a decreased total consumption globally will therefore magnify the benefits discussed here. On the other hand, an increase in consumption of oils and fats among people with malnutrition provides an affordable way to meet recommendations for healthier diets (66). From a producer level, integrating Camellia with other oil crops on existing low-efficient production areas as a locally adapted crop diversification strategy enables income diversification and farm resilience, for instance to climate impacts or soil infertility. This can be done through agroforestry strategies or Camellia’s ability to be cultivated on mountainous wasteland without extensive land cover change, which is in stark contrast for instance to oil crops like oil palm and soybean with major emissions driven by forest biomass or organic soil burning. At the intergovernmental policy arena, spatial optimization on a global scale is challenging due to the absence of a central planner. Addressing this issue through trade and internalizing externalities such as emissions, land use, and biodiversity impacts related to crop cultivation could be a viable solution (67,68). The World Trade Organization may, for instance, consider incorporating climate change and environmental impacts into its future negotiations. So, while Camellia farms can certainly not be regarded as a universal solution for sustainable vegetable oil production, it is crucial to recognize the potential benefits and develop strategies to harness them (15). Our approach to identify priority areas for crop substitution with a limited proportion of the replacing areas is suitable for other crop replacement evaluations in the context of diversified vegetable oil production or cropping systems in general. Thus, this work can chart a realistic pathway beyond diversified vegetable oil production with ramifications for multiple agriculture-related sustainable development goals.

## Methods

We first performed a comprehensive life cycle assessment (LCA) of six major vegetable oil crops - groundnut, soybean, palm, rapeseed, sunflower, and olive for a spatially explicit assessment of environmental and economic effects. Spatially cover distribution (5 arc minute; 1/12°; ∼10-km resolution) of the six main vegetable oil crop-specific information were taken from the Global Agroecological Zones (GAEZ) v4.0 database of the Food and Agriculture Organization of the United Nations (FAO) (69). Owing to data limitations and variation in the use of vegetable oil crops, we do not consider industrial products, feed or animal meal and assume that their production remains constant and unaffected under crop replacement (37). We then combined global vegetable oil crop distribution datasets with a Camellia global habitat suitability assessment, and production capacity modelling to enhance global vegetable oil production efficiency through optimized vegetable oil crops land allocation. Long-term climatic factors for land suitability assessment of Camellia oleifera cultivation and average monthly precipitation data for water use calculation of all crops were taken from the University of East Anglia’s Climate Research Unit CRU TS Version 4 (70). Growing stages, planting dates, and crop coefficients were adapted from previous studies and FAO Penman-Monteith *ET*_0_ (37,71,72,73). We did not allow for increasing the overall land area and oil yield loss from the replacement by Camellia cultivation locally. We visualize our methods in Fig. S1 and provide details below. A summary of all datasets and sources is provided in Table S24.

### Mapping global vegetable oil environmental and economic impacts

The environmental impact results are calculated through an LCA according to ISO norms (ISO14040, 2006; ISO14044, 2006) (74,75). For each crop’s vegetable oil system, we include the following life-cycle processes: the production of inputs (such as fertilizers and electricity), transport of the inputs to the plantation, and crop harvest in the agricultural stage and vegetable oil production in the industrial stage to calculate the GHG emission for producing 1-liter oil. For water and land use, we mainly consider the stage of crop production, since insignificant amounts of water and land are consumed outside the production phase (76). Limited data availability prevented us from extending our analysis to all production stages along the value chain. Furthermore, we also calculated the quantity of pesticide use for producing 1-liter oil and farmer income for per hectare land of Camellia production and all other vegetable oil crops considered here (i.e., groundnut, palm, rapeseed, soybean, sunflower, and olive). All the agricultural input data were collected by conducting a meta-analysis of 285 studies with 837 observations categorized as sub-national and national studies, covering 10,500 regional or farm-level inventories globally (Extended Section 1.2).

#### GHG emission

For the GHG emissions from land use change, we consider the above and below-ground C stock change, forest burning, and organic soil burning, but excluding the leaching, runoff, and induced non-CO_2_ emissions *sensu* Poor and Nemechek (16, 77). Specifically, we excluded the CO_2_ emissions from direct land use change of forest into cropping systems for groundnut, olives, rapeseed, and sunflower crops, as the total percentage of current crop area transformed from forest are on average less than 1% globally and over two decades (78) (Table S9). Moreover, the small fraction of forest related to land use change is difficult to address with available maps globally. We then used the FAO statistical database to estimate CH_4_ and N_2_O emissions from forest burning, and CO_2_, CH_4,_ and N_2_O emissions from organic soil burning at the country level for all six vegetable oil crops (Table S11, Table S12). As Camellia trees are not currently grown on a large scale globally, we used a global-scale Terrestrial Ecosystem Model (79) to quantify the potential impact of growing Camellia on land use change and carbon emissions. The agricultural inputs for the seven types of vegetable oil crops mainly include planting seeds or nurseries, fertilizers, agricultural film, pesticides, diesel, and electric energy consumed by agricultural machinery or water irrigation. For the oil processing stage, we collected the carbon emission from the published papers by referring to the flow of 1 liter of (refined) oil (Table S14). We estimate their GHG emissions through emission factors and results are expressed in kg CO_2_e kg^-1^ for crop fruit and kg CO_2_e liter^-1^ for vegetable oil, with the global warming potential (GWP) of CH_4_ and N_2_O being 34 and 298 kg CO_2_ e kg^-1^, respectively.

#### Water use

We use the CROPWAT 8.0 model and the FAO Penman-Monteith equation to calculate green water and blue water footprints (71,80), with considering the crop coefficient and potential evapotranspiration (Equation 1). Here, the green water footprint represents the amount of rainwater and soil moisture consumed by crops through evaporation or transpiration without contributing to surface runoff. The blue water footprint accounts for the surface water and groundwater consumed during the growth of crops. The grey water footprint also was calculated to represent the water consumption to dilute pollutant concentrations from fertilizer applications to reach natural background concentrations or to meet ambient water quality standards.

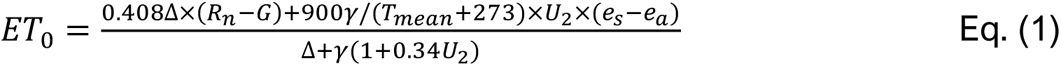

where *ET*_0_ is the potential evapotranspiration; Δ is saturated water pressure curve slope (kPa/℃); *R*;_*n*_ is ground surface radiation of crop canopy surface (MJ/(m*d)); *G* Soil heat flux (MJ/(m2·d)); *γ* is the wet and dry constant (KPa/℃); *T*_*mean*_ is the daily average temperature at 2 meters (℃); *U*_2_ is the wind speed at 2 meters (m/s); *e*_*s*_ is the saturated water pressure (Kpa); *e*_*a*_ is the actual water pressure (Kpa). The calculation process of Δ, *R*_*n*_, *γ*, *T*_*mean*_, and *U*_2_ are referred to from "FAO Irrigation and Drainage Paper, No. 56, Crop Evapotranspiration" (71) and the details are explained in Extended Section 3.2.3.

#### Land use

By taking the difference between temporary and permanent crops into consideration, we divided the vegetable oil crops into two main types namely temporary crops (groundnut, rapeseed, soybean, and sunflower) and permanent crops or orchard crops (Camellia, palm, and olive). These temporary and permanent oil crops have different cropping duration, growth forms, and life cycle characteristics which influence land use (15). Land use for each vegetable oil crop was calculated from occupation time and inverse crop yields in each region, with actual yield for the current six types of vegetable oil crops and three scenarios of potential yield for Camellia. Our land use estimates for each vegetable oil are higher than those reported in Poore and Nemecek (16), because we report the global average land use intensity based on our detailed regional results.

#### Pesticide use

Unlike the life cycle assessment required to estimate the processed-based GHG emission, crop water use, and land use, pesticide use is assessed directly based on meta-analysis and official statistical data, with studies being weighted in three ways: 1) within-region country weights; 2) between-region country weights; and 3) between-country weights to enable one geography reflecting another, where data was unavailable. For more details see Extended Section 1.2.3. Here, the pesticide use excludes minerals used for pest management, such as Sulphur or Copper, to make the amount of substance and the amount of active ingredients comparable (Table S2).

#### Farmer income

The crop-specific total production was calculated based on specific rainfed or irrigated yield and harvested areas. For the price of the six vegetable oil crops considered here, we used the average value from 2017 to 2021 obtained from FAOSTAT to get the producer prices in each country (USD Tonne^−1^) (Table S23) (2). As currently Camellia is mainly planted in China, the farmer incomes coefficient information was taken from the National Development and Reform Commission (81) and producer prices of Camellia seed were set as 19.74 RMB/Kg (100 USD = 660 RMB) (82). The farmer’s net profit varies based on the agricultural and labor inputs, such as the pesticide and fertilizer use intensity. Hence, for each agricultural commodity selected, we assumed a 20% margin of profit to obtain the opportunity cost for each vegetable oil crop’s fruits, as a likely conservative (that is, high) estimate based on profit margins in the USA (non-small farms) (83), some EU countries (84) and Canada (85). This simplified assumption results from limited data availability and variable profit margins, in particular with farm sizes (86).

### The potential distribution and yield supply of Camellia cultivation

#### Land suitability assessment

To identify the potential of land distribution for planting Camellia, we selected 11 indicators based on a comprehensive literature review from 3 aspects of climatic, topographic, and soil factors from multiple databases to model the spatially explicit distribution of suitable areas. For a full description of indicator selection, modelling details, and data sources, please see Extended Section 2. As the weight of each indicator varies in published papers, we used a “random forest” machine learning algorithm to identify the weight for each indicator of current Camellia planting observations. Random forests are robust to overfitting and variable importance when modelling crop distributions and predicting the yields (87, 88). The decision to use a random-forest approach was supported by its superior performance metrics compared to other machine learning approaches (Fig. S9).

Beyond the natural environmental viability that we considered from the land suitability assessment, we further identified five additional constraint factors (land use type, terrestrial and inland protected areas, world roads areas, intact forest landscapes, and global key biodiversity areas) that are likely to limit the suitable areas for large-scale Camellia cultivation development due to ecological protection or social constraints. For the land use constraints, we ultimately considered excluding the land cover with rainfed, irrigated, or post-flooding cropland, urban areas, water bodies, and permanent snow or ice. After obtaining these spatially explicit data layers of five constraint factors, we resampled each at a 0.083° resolution to match with the 11 graphic data layers. We first processed the 11 indicators layer as described in Table S16 to classify each grid into four classes, not suitable, moderately suitable, suitable, or very suitable. For the five additional constraint factors, we assume that some areas (for example, the areas within the protected areas, the key biodiversity areas, or the intact forest) will be completely unsuitable for planting Camellia. These five layers were further processed to assign double-valued with 0 and 1, where 0 represents not suitable and 1 represents suitable. These layers were then multiplied together to create the overall global map of land suitability shown in Fig. S11. Field verification in Extended Section 2.4 with an 88.20% cross-validation ratio shows the high reliability of our land suitability assessment result.

#### iGAEZ modelling on potential yield

To simulate crop yields globally for land suitable areas of Camellia, we developed a new approach called iGAEZ (improved Global Agro-Ecological Zones model) based on the GAEZ model (69). While GAEZ does provide evaluation of the biophysical limitations and production potentials of land, iGAEZ provides realistic crop yield combinations considering the whole spectrum of climatic and agricultural conditions. iGAEZ model generates the optimal crop parameter of the whole growth cycle to provide the best realistic crop yield combinations. Doing this, the improved model takes regionally differentiated climatic – social - economic status and crop condition into account. The details of each procedure in the iGAEZ model are detailed in Extended Section 3 with the results of the photosynthetic potential yield, light and temperature potential yield, climatic potential yield, and natural comprehensive potential yield, respectively (Fig. S13).

#### 5.2.3 Scenario design

We develop three scenarios in this study, namely, a conservative scenario (S1), where the Camellia yield potential stays close to business-as-usual and reflects actual yield. In the strategic improvement scenario (S2), we assume implementation of advanced cultivation techniques and optimized input use to achieve a 25% increase in Camellia yield over current levels. In a technology-policy-enabling scenario (S3), both technology advances and policy improve to promote diversified vegetable oil production with woody and environmental oilseed yield. Every scenario is contingent upon the one before. Specifically, in S1, estimates of yield growth were set as 50% of the final natural comprehensive potential yield at any spatial scale for best indicating near the actual yield. The actual yields for Camellia *seeds* are mainly collected from China Agriculture Outlook (89) and the 2019 China Forestry and Grassland Statistic yearbook (90). The current actual yield records of Camellia seed in China for Anhui, Hunan, and Guangxi Province are 6.7, 7.8 and 6.5 t/ha, respectively (91,92), which are still much lower than their yield potentials (that is, 14.4 t/ha) but close to half of the yield potential. In S2, we made 75% of the potential yield as an attainable or exploitable yield ceiling of Camellia. Due to diminishing profits from investing in additional production inputs and labor as yields approach the potential yield ceiling, present farm yields tend to plateau when they reach 75–80% of potential yield, also known as the exploitable yield ceiling (93,94). In S3, we assume a 100% potential yield of Camellia. This assumes that technology-oriented policies are implemented and yield improvement through technologies make 100% potential yield attainable.

### Spatial prioritization of Camellia crop-replacing model

We applied a multicriteria optimization framework (95,96) to identify global priority areas for replacement from current vegetable oil crops to Camellia plantations and to quantify trade-offs and synergies across five objectives, including GHG emission, water use, land use, pesticide use, and farmer income. To do so, we: (i) estimated the current six main types of vegetable oil crops planted worldwide and their producing environmental impacts and farmer income; (ii) estimated potential long-term replacing benefits by Camellia plantation across five objectives; (iii) implemented a multicriteria optimization algorithm based on linear programming to show the spatial prioritization ranking for Camellia planting areas; and (vi) simulated different global replacement scenarios based on each objective or comprehensive consideration under different Camellia yield potential. Our study is global with a focus on all terrestrial regions suitable for production of the six main types of vegetable oil crop from −80° to 80° latitude and from −180° to 180° degrees longitude.

#### Trade-offs among five objectives for sustainable vegetable oil production

To estimate the potential benefits of replacement from current vegetable oil crops to Camellia plantation, we set our five objectives when implementing the crop replacement to achieve a sustainable vegetable oil production system, that is, (1) maximize GHG emission reductions, (2) maximize water use reductions, (3) maximize land use reductions, (4) maximize pesticide use reductions, and (5) maximize farmer income increasing.

We first used the total oil production or the total harvested areas for each oil crop to calculate the current total environmental and economic impacts based on the previous results per liter of oil or hectare area in Section 1.3. For each grid, the current (2015) vegetable oil production from each crop *z* of rainfed (*OilPro*_*Cur*,*R*;*ai*,*i*,*z*_) and irrigated (*OilPro*_*Cur*,*Irr*,*i*,*z*_) were calculated as Equation 2.

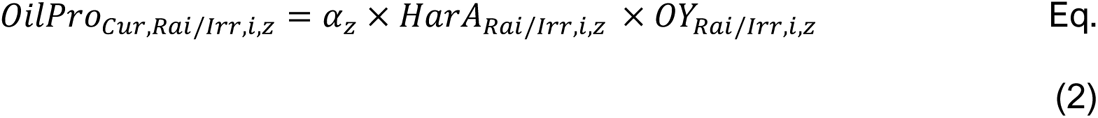

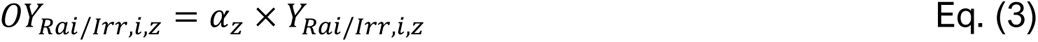

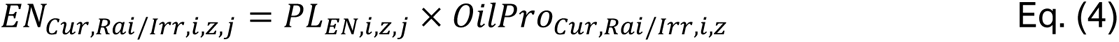

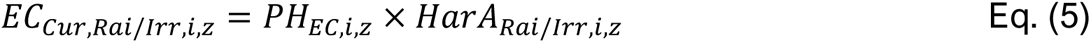

where *HarA* is the harvested area (ha) for each vegetable oil crop in grid *i*; *z* is the six types of vegetable oil crops with the value from 1, 2, 3, …, 6 referring to groundnut, soybean, palm, rapeseed, sunflower, and olive, respectively; *α*_*z*_ is the oil extraction ratio for different crops to produce 1 liter (refined) oil (Table S6). *OY* is the oil yield (kg/ha) by considering the crop fruits yield *Y* and oil extraction ratio *α*_*z*_; the subscripts *R*;*ai* and *Irr* represent two cropping systems in rainfed and irrigated, respectively; *PL*_*EN*,*i*,*z*,*j*_ is the environmental impact of producing per liter oil and *j* is the value of 1,2,3, and 4, representing the environmental indicators with GHG emission, water use, land use, and pesticide use, respectively. *PH*_*EC*,*i*,*z*_ is the economic impact with the indicator of net farmer income per hectare. *EN*_*Cur*,*Rai*/*Irr*,*i*,*z*,*j*_ and *EC*_*Cur*,*Rai*/*Irr*,*i*,*z*_ are the total environmental and economic impact in grid *i*. Then, we estimated the benefits of Camellia planting replacement, which can be expressed as:

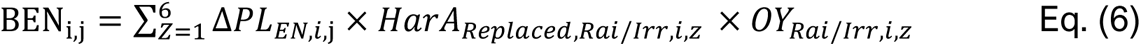

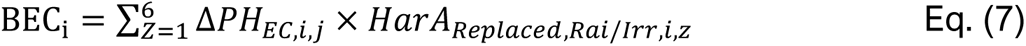

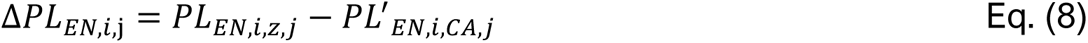

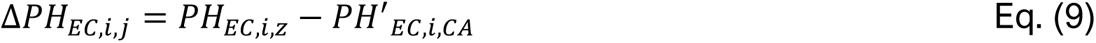

where BEN_i,j_ is the total replacing benefit of environmental indicator of *j* and BEC_i_ is the total replacement economic benefit. Especially, after crop replacement, it will possibly require additional transportation and transaction costs, which farmers do not bear. We acknowledge that our model has the limitations of not considering the additional costs and assumption of the constant harvested area under crop replacement. Thus, we made a very conservative estimation of replacing benefits by using the oil yield of original crops instead of Camellia in Eq. (6) to offset partial impacts as the crop-replacing constraint of higher oil yield compared to original crops. *PL*^′^_*EN*,*i*,*CA*,*j*_ is the environmental impact of producing per liter Camellia oil under three potential yield scenarios; *PH*^′^_*EC*,*i*,*CA*_ is the net farmer income per hectare of Camellia plantation. Δ*PL*_*EN*,*i*,j_ and Δ*PH*_*EC*,*i*,*j*_ are the differences between the current crop and Camellia per unit of the environmental and economic impact. *HarA*_*Replaced*,*Rai*/*Irr*,*i*,*z*_ is the current harvested area (ha) of replaced locations.

#### Crop-replacing constraints

There is a need to determine the crop replacement constraints to ensure successful implementation of crop replacement. The first constraint in Equation 10 limits the location of the vegetable oil crop to be replaced, which depends on the oil yield of Camellia (OY_i,CA_). The requirement of higher or equal to the current vegetable oil yield is to ensure no oil yield loss. In the second constraint for scale, *A*_*i*_ is the total percentage of the current six types of vegetable oil crops harvested areas to be replaced by Camellia. Based on recommendations from the literature (95), the proportion of the areas replaced or fallowed is usually limited to a maximum value of ΔCL = 20% to ensure global food market stability. In the study, the maximum value of ΔCL = 23.5%, which is our estimated proportion of all suitable areas for planting Camellia within the current vegetable oil production areas to the total six main vegetable oil harvested areas. Here, we assumed that the replaced harvested areas would remain the same with the current crops once identifying replacement, which is for maintaining defined crop-specific production and vegetable oil volumes (88). Furthermore, the current crop-specific producing area will not expand to minimize the global vegetable oil-producing land extent and directly avoid global burden-shifting, especially for the land use pressure. The third constraint was implemented based on pursuing different objectives. If we consider maximizing additional environmental impact reductions, it requires Δ*PL*_*EN*,*i*,j_ ≥ 0 meaning the environmental impact of producing per liter Camellia oil is lower than current vegetable oil. If under the comprehensive consideration of all five objectives, it requires both Δ*PL*_*EN*,*i*,j_ ≥ 0 per liter oil and Δ*PH*_*EC*,*i*,*j*_ ≥ 0 per planting hactare area.

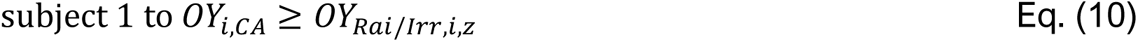

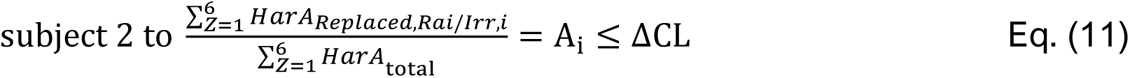

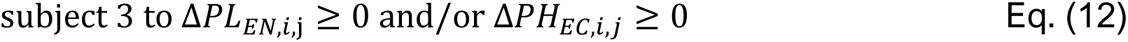

#### Spatial prioritization ranking and scenarios setting

To identify the priority areas for Camellia replacement to the current vegetable oil crops, we ran a multicriteria optimization algorithm based on an objective function that defines the amount of area to be replaced in each planning unit to maximize benefits. By modelling 108 scenarios of replacement prioritization (3 potential yield levels of Camellia × 6 types of main vegetable oil crops of groundnut, soybean, palm, rapeseed, sunflower, and olive × 6 targets by trade-offs among each five objective and/or the comprehensive consideration of all), we aim to interplay with land potentially spared from low vegetable oil production efficiency to achieve different replacing targets on environmental and/or economic benefits. There are 100 priority levels in each scenario divided as the equal grid number with the spatial layout of targeted replacing areas and then ranked as the high benefits of each five objectives and/or the comprehensive. All spatial prioritization ranking maps, unless otherwise noted, were the result of aggregating the nested sets of priorities for total benefit formulations by replacing six vegetable oil crops. For the comprehensive consideration of all five objectives, an equal weighting was chosen to understand how the five components trade off and to focus on the layout and replacing scale of priority on a macro scale.

## Supporting information

Text Section 1-3; Figures S1-S34; Table S1-S24

## CRediT authorship contribution statement

Conceptualization - SZ, FC, TCW; Formal Analysis - SZ, CZ; Data Curation - SZ, KFD; Writing – Original Draft Preparation - SZ, TCW; Investigation - SZ; Writing – Review & Editing - All authors; Supervision – TCW

## Acknowledgements

Funding was provided by a Westlake University start-up fund to TCW.

## Appendix A. Supporting information

Supplementary data associated with this article can be found in the online version at

