## Supplementary material for "Diversified global vegetable oil production can mitigate climate change and achieve socio-environmental co-benefits": Text Section 1-3; Figures S1-S34; Table S1-S24

**This file includes:**

Section 1: Life cycle assessment of seven vegetable oil crops

Section 2: Global land suitability assessment of *Camellia oleifera* cultivation

Section 3: Projection of *Camellia oleifera* production capacity

Figure S1 to S34

Table S1 to S24

References

### Table S1 to S23:

**Table S1.** Fatty acid and bioactive content of different varieties of Camellia oil (1-5)

**Table S2.** 837 unique records of agricultural input inventories covering ~10,500 regional or farm level input information in 43 countries

**Table S3.** Coefficient of carbon footprint of raw materials inputs in edible vegetable oil crops production

**Table S4.** Country specific electricity emission factors GWP100 cc fb conversion factors

**Table S5.** Agricultural inputs of Camellia oleifera plantation in the entire lifecycle information (80 years)

**Table S6.** Processing information for producing 1 liter oil of seven types of vegetable oil crops

**Table S7.** Life cycle inventory model-input and output operational data for Camellia sinensis nursery

**Table S8.** Parameters values and variance-covariance matrix of $N_2O$ emission model

**Table S9.** The current (2009-11 avg.) area of six vegetable oil crops transformed from forest (amortized over 20 years)

**Table S10.** $CO_2$ emissions from land use change

**Table S12.** $CO_2$, $CH_4$ and $N_2O$ emissions from burning of organic soils

**Table S13.** The biomass regression models for Camellia oleifera tree species

**Table S14.** The greenhouse gas emissions in the processing stage of vegetable oils

**Table S15.** The crop coefficients (Kc) of oil crops for water use with the FAO Penman-Monteith ET0

**Table S16.** Land suitability assessment system of Camellia oleifera cultivation

**Table S17.** 1,263 records for the weight identification in land suitability assessment system of Camellia oleifera cultivation

**Table S18.** 1,958 field sample sites of Camellia cultivation

**Table S19.** The value of parameters for estimating photosynthetic potential productivity

**Table S20.** Classification and weight of land effective coefficient indicators

**Table S21.** Comparison of fruit characteristics of four Camellia oleifera clones

**Table S22.** The total replacing areas for each vegetable oil crops under different objectives across scenario

**Table S23.** The statistical data of the average value from 2017 to 2021 obtained from FAOSTAT to get the producer prices (USD/Ton)

**Section 1: Life cycle assessment of seven vegetable oil crops**

The global food systems are facing the pressing challenge of providing sustenance for a growing global population while minimizing environmental damage and resource depletion (6,7,8,9). Vegetable oil, as one of the primary energy sources in human diets, plays an indispensable role in the current food system (10,11). Over the past decade, the demand for vegetable oils has surged by 43.8%, driven by rapid population growth and economic expansion worldwide (12,13). This increased consumption of edible oils has consequently spurred a 36.2% growth in the cultivation of oil-bearing crops (14). However, expanding the production and processing of vegetable oil crops is likely to exacerbate environmental pressures, particularly in terms of GHG emission, water consumption (blue, green, and gray water use), and land use (15,16). Here, we used the life cycle assessment method to calculate the current six main types of vegetable oil crops' environmental impact in per liter oil, which include groundnut oil, palm oil, rapeseed oil, soybean oil, olives oil, and sunflower oil. Furthermore, based on the results of land suitability and potential yield modelling of global *Camellia oleifera* production, we estimated the corresponding environmental impact at multi-yield level.

### 1.1 System boundary and functional unit

The environmental impact results are calculated through an LCA according to ISO norms (ISO14040, 2006; ISO14044, 2006) (17,18). The system boundary is the whole agricultural and industrial production “from cradle to farm gate”, including the tree seeding sowing, cultivation, and harvesting process of crops, and all the processes of vegetable oil production (Fig. S2). The system boundary includes upstream processes (the production and transportation of agricultural input materials), core agricultural production procedures including crop cultivation (from seeds or tree saplings sowing, water irrigation, application of chemical and fertilizer to crop harvesting), and vegetable oil production in the industrial stage. However, the downstream processes such as vegetable oil distribution, transport, and food consumption were not considered in our study. For the functional unit (FU), the agricultural phase is calculated based on 1 ha of the cultivated land surface, and the oil refining phase takes 1 liter of vegetable oil production as quantitative analysis for all relevant inputs and outputs of the environmental impacts. This can be the quantitative standard for decision-makers and researchers to make comparisons between different vegetable oil crops on footprint and then to formulate countermeasures and further studies conduction.

### 1.2 Inventory and data source

#### 1.2.1 Meta-analysis approach for data collection of the current six main types of vegetable oil in agricultural phase

Data collection for building the inputs and management inventory is the key to conducting the life cycle assessment. However, there is very limited statistical information available on agricultural inputs on the upstream of life cycle by crops in different regions, such as the pesticides, diesel, films, and irrigation electricity consumption data. Thus, we conduct a meta-analysis approach that can fill this gap by converting the collected information in a single observation into standard inputs across regions at a global level.

**(1) Study inclusion criteria**

We initiated a literature search using the following 11 criteria to identify studies:

**a.** Literature is included in peer-reviewed journals, or are PhD thesis and published in Web of Science (WOS) or China National Knowledge Infrastructure (CNKI), International Organization for Standardization (ISO) compliant reports, Life Cycle Assessment (LCA) databases, or conference proceedings with clear data and methods;

**b.** Literature is published between 2000 and July 2022 in print or online version;

**c.** Report results not presented in another included study;

**d.** Assess commercially farmed products;

**e.** For the seven oil crops, data from the published literature should report real yield and inventory data, but not simulated;

**f.** LCA or similar methodology was used;

**g.** The environmental impacts were calculated aligning with our system boundary or can provide sufficient inventory data to recalculate;

**h.** The environmental impacts were calculated aligning with our functional units, or can make recalculation possible;

**i.** Calculate GHG emissions, characterized under 100-year factors of IPCC Fifth Assessment Report with climate-carbon feedbacks, or can make recharacterization possible;

**j.** Calculate on- and off-farm land use, or make calculation possible;

**k.** The literature included is limited to those with specific site information or clear geographic data in farms, regions, or countries.

**(2) Aggregating results**

Results were included as a separate observation (line of the database) when they:

**a.** Represented different practices or food production systems, e.g., input level, rotation, cultivar

**b.** Represented significantly different geographies, e.g., regions or countries.

Otherwise, results were averaged into a single observation with standard deviations calculated across countries. Results were averaged across years to ensure the independence of each observation.

**(3) Literature research and systematic review**

A comprehensive approach was used by: (1) searching for publications using the terms "life cycle analysis" OR "life cycle assessment" OR "GHG emissions" AND the crop name OR corresponding relevant oil product name, in Google Scholar, WOS, or CNKI; (2) following references within publications and citations to publications; (3) identifying LCA conference proceedings; (4) and identifying online LCA datasets.

Based on literature research and systematic review form the published papers or databases, our meta-analyses of life cycle assessments aggregate and synthesize data from individual LCAs to provide estimates of the environmental impact of vegetable oil products. Previous analyses compared the impacts of food commodities, which contain vegetable oil products such as soybean oil, and olive oil, from the literature published between 2000 and June 2016. Here, we used agricultural inputs and food processing data available from Poore and Nemecek (2018) (19), which is being converted into an online environmental database named HESTIA (20). Then, we paired the same recalculation methods (19) and updated the latest data from the literature published after 2016 to construct a new dataset for understanding the environmental impact focusing specifically on vegetable oil products.

**(4) Data Extraction**

By conducting a meta-analysis approach strict on our 11 inclusion criteria, we collected 837 unique records of agricultural input inventories, covering ~10,500 regional or farm-level input information in 43 countries (Table S2, Fig. S3). The geographical coordinate data information of each record was obtained by the following three methods: 1) Observations' site information was directly acquired. In all studies, the names of sampling crop, production system types, geographical coordinate data, and number of farms assessed and related life cycle inventory (LCI) information were mentioned in detail; 2) In the second method, the observations' names were retrieved using their respective geographical coordinates. The latitude and longitude coordinates of sampling sites were acquired by Geosharp (Version 2.1, https://www.udparty.com). Systematic deviations of geographical coordinate data from different studies were transformed using coordinate transformation software, and 3) The third method included digitalization of observations' site images. A spatial distribution diagram of site information was extracted from these reports using ArcGIS (Version 10.1, http://www.esri.com) and Getdata Graph Digitizer (Version 2.25, http://www.getdatagraph-digitizer.com).

Of these included 837 unique records in 285 studies, supplemented with 272 records at national level and 565 at sub-national, showing a descending order of soybean (255) > rapeseed (232) > olives (131) > palm (93) > sunflower (71) > groundnut (55). Observations are concentrated in North America, Europe, Oceania, China, The United States, and Brazil, but limited in most areas of Africa, proving the necessity of study weights that enable one geography to reflect another.

#### 1.2.2 Data collection of C*amellia oleifera* oil in agricultural phase

The agricultural inputs data of *Camellia oleifera* includes two aspects: (1) **Foreground data** refers to primary relevant data of the *Camellia oleifera* planting system, including nursery, land preparation, planting, processing, harvesting, material transportation, and other agricultural inputs data. Foreground data are mainly collected from the China Statistical Yearbook, China Rural Statistics Yearbook, China Forestry and Grassland Statistics Yearbook, and Compilation of cost-benefit information on agricultural products in China (21,22,23,24). (2) **Background data** refers to production information of agricultural raw materials used in the *Camellia oleifera* planting process, such as plastic film, fertilizers, pesticides, electricity, and diesel. These data are collected from the Ecoinvent database and published literature (Table S3) (19,25-37). The country-specific electricity emission factors are shown in Table S4, cited by Brander et al. (2011) (38). Finally, as shown in Table S5, the environmental impact assessment scope of the agricultural stage is displayed, taking *Camellia oleifera* plantation in Hunan Province as an example, providing a data list of *Camellia oleifera* planting's entire lifecycle information (80 years) within a 1-hectare land area.

The water requirement of *Camellia oleifera* is calculated based on precipitation data, crop coefficients, and the crop growth period. Precipitation data is obtained from daily measurements at meteorological stations worldwide. *Camellia oleifera* is an evergreen tree species, and the root system is not well developed in the first five years. During the early growth stages, they are sensitive to both waterlogging and drought. Therefore, to ensure normal bud germination and flowering, no irrigation is considered during the spring irrigation and dormancy period at the end of March. By the end of October, the *Camellia oleifera* has matured, with different levels of water surplus or deficit having little impact on the growth and quality of the fruits. Thus, we consider no specific requirement of irrigation for the whole stage of *Camellia oleifera* plantation. The monthly crop coefficient $K_c$ of *Camellia oleifera* is obtained from Wang et al (Fig. S4) (39). Throughout the various growth stages of *Camellia oleifera* trees, influenced by the comprehensive effects of climate conditions and the biological characteristics, the reference crop potential evapotranspiration ($ET_0$) shows a unimodal change, starting from increasing and then decreasing. The peak period occurs in July and August, during which the fruit and leaf area of *Camellia oleifera* trees are at their maximum, exhibiting vigorous growth and high rates of evaporation and transpiration. The crop coefficient ($K_c$) shows a bimodal pattern throughout the whole year, with high values mainly occurring in June, August, and September, corresponding to the two peak water demand periods ("fruit expansion period" and "ripening period").

(1) During the "flowering and fruiting period" (January to April), the coverage of *Camellia oleifera* plantations is relatively low, and water consumption is mainly needed through evaporation. Therefore, both the reference crop potential evapotranspiration ($ET_0$) and the crop coefficient $K_c$ are low during this period.

(2) During the "fruit expansion period" (May to September), the fruits require a significant amount of water, and the nursery needs to continue proper management, including the removal of shading structures and timely fertilization, resulting in a peak water demand for *Camellia oleifera* trees.

(3) During the "abundant fruit harvest period" (October to December), strict water control is necessary to prevent fruit cracking. The water requirement is extended, and the water consumption of *Camellia oleifera* trees decreases as the leaves are harvested in October. Additionally, the evaporation between trees also decreases compared to the previous stage. The monthly averages of $ET_0$ and $K_c$ decrease during this stage.

As shown in Figure S4, the water consumption of *Camellia oleifera* trees from the "flowering and fruiting period" to the "abundant fruit harvest period" follows a similar seasonal pattern at both experimental sites. Thus, we determined the average values of

$K_{c-ini}$, $K_{c-mid}$, and $K_{c-end}$ to be 0.31, 0.99, and 0.66, respectively, for the months' mean value of January to April, May to September, and October to December.

#### 1.2.3 Weights on vegetable oil crops

**(1) Within-region in country weights**

The present global Köppen-Geiger climate classification maps (40) classified climate into five main classes at 1 km resolution were used to group regions in the country by similarity. Study representativeness within a region in a country was determined first if the agricultural input observations were obtained from global level dataset such as on the groundwater used for irrigation (41), the World of Organic Agriculture (42), or from national or regional statistical yearbooks or censuses (24), or the results reported by the published papers in a sub-national production system (43).

**(2) Between-region in country weights**

For those regions without observations, agricultural input inventory was weighted by the region-level reference and the total number of farms assessed within a country. Study observations were split into four groups based on the difference of region-level reference (- = national; C1 = region or province level; C2 = sub-regional level; S = point location; A = average of point locations). We set the weight of different study areas' levels in descending order of national (30%) > C1 (25%) > C2 (20%) > S (15%)> A (10%). In particular, if the values reported by the author didn't give specific information, we assumed the number of evaluated farms equals one for calculating the share of total farms.

**(3) Between-country weights**

As the uneven distribution of study observations, we first determined whether the crop production in the countries where we collected observations exceeds 75% of the total global production, showing a descending order of soybean (91.77%) > palm (91.55%) > rapeseed (77.02%) > olives (66.14%)> groundnut (60.12%)> sunflower (51.87%).

For the crops of soybean, palm, and rapeseed, where ≥75% of global production was represented by observations, and the crop of *camellia oleifera* currently mainly planted in China, the identification of agricultural input inventory across different regions in the country was weighted following the rules of “(2) Between- region in country weights”.

Based on field experience in low-yielding regions, nutrients, and water, rather than genetics, can hinder yields from increasing (44,45). Thus, for the crop of olives, groundnut, and sunflower, where <75% of global production was represented, macro-nutrient input levels (46) and global Köppen-Geiger climate classification (40), were used to create a 2×5 matrix in different regions across countries in high-low macro-nutrient input and main climatic zones. Specifically, if the national information on macro-nutrient input level is not available, the Human Development Index (HDI) was utilized as an alternative metric (19,47).

#### 1.2.4 Oil refining phase

In the oil refining phase, we take 1 liter vegetable oil production as a quantitative analysis for the environmental impacts. Vegetable oils are complex mixtures consisting of different fatty acids, sterols, phospholipids, and triacylglycerols (48). The characteristics of these fatty acids, such as chain length, degree of saturation, and branching, influence the oil density (49). Generally, vegetable oils with a higher proportion of saturated fatty acids tend to have a higher density than those with a higher proportion of unsaturated fatty acids (50,51). We determined the fresh fruits required to produce one liter of vegetable oil by considering the density of different plant oils and the corresponding oil extraction rate. The density (Kg per liter at 20℃) shows a descending order with soybean oil (0.93) > sunflower oil (0.92) = rapeseed oil (0.92) > palm oil (0.89) > olive oil (0.86) > groundnut oil (0.80) = *Camellia oleifera* oil (0.80) (19,52). For the oil extraction rate, we get the mean value for different oil crops from the literature review, as shown in Table S6 (8,23,53-63). Finally, the weight of fresh fruits required to produce one-liter vegetable oil for different crops show shows a descending order with *Camellia oleifera* oil (6.96 kg) > soybean oil (5.17 kg) > olive oil (4.74 kg) > palm oil (4.68 kg) > rapeseed oil (2.42 kg) > groundnut oil (2.40 kg) > sunflower oil (1.99 kg).

More specially, we introduced detailed information on *Camellia oleifera* oil process and refining procedures. Currently, the production processes of *Camellia oleifera* oil mainly include solvent extraction, hot pressing, and cold pressing.

① Solvent extracted *Camellia oleifera* oil: Chemical leaching extraction mainly uses No. 6 light petrol as a solvent to fully soak the raw materials, and then extract oil at high temperature. This method has a high oil extraction ratio but with the disadvantage of being contaminated by chemical solvents and residues. It is difficult to completely remove solvents from edible vegetable oil, and the solvents used are mostly light gasoline. If the quality is impure, it may contain toxic compounds such as benzene and polycyclic aromatic hydrocarbons, which can pose health risks for consumers in the long term.

② Hot-pressed *Camellia oleifera* oil: Hot pressing is the extraction of oilseed crops after frying and roasting (at temperatures higher than 120-130°C), which results in an extraordinarily fragrant odor, darker color, and higher yield but fewer residues and easier to preserve. The disadvantage is that the crude oil extracted from high-temperature processed oilseeds has a dark color and increased acid value. Therefore, the crude oil must be refined before consumption. Additionally, high-temperature pressing causes significant loss of bioactive substances, such as vitamin E tocopherols and carotenoids, resulting in a waste of resources.

③ Cold-pressed *Camellia oleifera* oil: Cold pressing is conducted at temperatures below 60°C to retain the nutrient components as intact as possible. Cold-pressed oil does not require additives and can be stored for a long time, making it more valuable in the market. The final product of cold-pressed oil preserves the natural flavor and color of the oilseeds, as well as the bioactive substances, such as vitamin E with anti-aging functions and phytosterols with skin-enhancing effects and metabolic functions. Cold-pressed oil retains the original taste and offers a healthier choice for a quality lifestyle. With the technology development, the combination of low-temperature cold pressing and twin-screw pressing technology has significantly reduced the residual content and markedly improved the oil yield from *Camellia oleifera* seeds. This method has become highly favored in factory production, as evidenced by products such as Xiangchun *Camellia oleifera* Oil, Dakang Era *Camellia oleifera* Oil, and Thousand Island *Camellia oleifera* Oil. Although residual oil can be extracted from seed cakes can be extracted by solvent leaching method, the oil content is relatively low. This study primarily evaluates the environmental impact of using the low-temperature cold pressing method to produce *Camellia oleifera* oil. The specific process flow is as follows (Figure S5) (64,65,66):

(1) Oil extraction: Fresh *Camellia oleifera* fruits are first subjected to preliminary impurity screening. The fruits are then dried (by either sun drying or air drying) and mechanically shucked to obtain seeds with moisture content below 10%-12%. After

secondary impurity removal, the seeds are processed by machine to separate the kernels from the shells, with the shell content in the kernels controlled at 12%-15%. Currently, the shucking equipment mainly includes centrifugal impact and hammer types, both of which can be used for dry and wet processing. The processed dry *Camellia oleifera* seeds are then transported to the oil press by spiral lifting and pressed to ensure that the chamber temperature does not exceed 80°C during pressing. The entire production process takes place at ambient temperature, with proteins in seeds remaining non-denatured and the kernels remaining soft. The crude oil obtained after pressing is filtered through a centrifugal oil filter to obtain the pressed virgin *Camellia oleifera* oil. This process of producing *Camellia oleifera* oil can retain the unique flavor and trace nutrients of *Camellia oleifera* oil while having minimal contamination as no organic solvents are involved.

(2) Oil refining: To meet the commercial standards of vegetable oil, it is necessary to refine the pressed virgin *Camellia oleifera* oil, which is commonly known as the "six extractions" process, including degumming, deacidification, dehydration, decolorization, degreasing, and deodorization.

1) Degumming. The colloidal substances in *Camellia oleifera* oil primarily refer to a mixture of phospholipids, proteinaceous colloids, and other impurities. Phospholipids can affect deodorization and the decolorization process during steam treatment, as well as chelate with metal ions, leading to increased oxidation and impacting oil stability. The degumming of crude oil is essential for removing all colloidal substances and other impurities from *Camellia oleifera* oil, while also generating valuable by-products. Degumming methods mainly include acid degumming, hydration degumming, and biotechnological degumming. By leveraging the principles of the hydrophilicity of colloid-soluble impurities and the transformation of non-hydrated colloids into hydrated forms in the medium, the colloid-soluble impurities are hydrated, expanded, coagulated, and separated. Here, we evaluate the chemical input for acid degumming, with citric acid or phosphoric acid commonly chosen as degumming agents. The specific operational procedure involves adding 0.1% phosphoric acid based on the oil mass at room temperature and rapidly stirring for 40 minutes.

2) Deacidification. The purpose of deacidification is to neutralize the free fatty acids in crude oil by adding alkali. Deacidification methods mainly include alkali refining, membrane separation, esterification deacidification, and supercritical $CO_2$

deacidification, among which alkali refining is the most commonly used. The initial refining temperature is set at 30°C, and 0.2% excess sodium hydroxide solution is added at a stirring speed of 60 r/min. Then, the temperature is raised to 60°C while stirring at 30 r/min, and the mixture is allowed to stand for 8-10 h, during which the free fatty acids precipitate out as salts that can dissolve in water. The mixture is then washed twice with hot water at 5%-10% of the oil mass, and the hot water temperature is slightly higher than the oil temperature during washing.

3) Dehydration. After deacidification, *Camellia oleifera* oil contains tea saponin and requires water washing to remove it. After tea saponin removal, the dehydrating process is conducted under vacuum conditions to achieve optimal subsequent decolorization effects. The optimal temperature for vacuum dehydration is 110°C, with a vacuum pressure of 1.3 kPa. Stirring at a speed of 60 r/min, the dehydration process takes approximately 40-50 minutes until there is no water mist in the dehydration vessel.

4) Decolorization. Color is one of the important criteria of good *Camellia oleifera* oil, and it is a necessary inspection item in the process of oil processing, storage, and sales, serving as an external manifestation of product quality. The main purpose of decolorization is to remove pigments, residual pesticides, trace metals, residual trace soap particles, phospholipids, glycerides, and polycyclic aromatic hydrocarbons from the oil. There are adsorption and chemical methods for decolorization, with adsorption being a common method for *Camellia oleifera* oil. Typically, the addition of 2% by mass of activated white clay or 0.5% by mass of activated carbon for 0.5 hours is chosen. Under vacuum conditions, effective decolorization of *Camellia oleifera* oil can be achieved by stirring and heating after adding adsorbent.

5) Degreasing also known as dewaxing. *Camellia oleifera* oil contains long-chain triglycerides and waxes, which are cyclic hydrocarbon impurities. Under low-temperature conditions, these impurities can suspend and precipitate from the oil over time, reducing transparency and directly affecting the appearance of *Camellia oleifera* oil. The degreasing filtration of *Camellia oleifera* oil is carried out under low-temperature conditions using crystallization, crystallization tanks, and filtration machines, utilizing the height difference for self-flow.

6) Deodorization. The principle of deodorization relies on the difference in volatility between odor-causing substances and triglycerides. Under high-temperature and high-vacuum conditions, a process utilizing water vapor is employed to remove the odor-causing substances. Deodorization not only eliminates odor-causing substances in

the oil, improving the flavor of the food but also enhances the smoke point of the oil. *Camellia oleifera* oil undergoes deodorization treatment at a vacuum level of 260 kPa and a temperature of 180°C. The steam consumption is approximately 3% to 8% of the weight of the *Camellia oleifera* oil.

### 1.3 The life cycle impact assessment

#### 1.3.1 GHG emissions

The system boundary of this study is defined using a "cradle-to-gate" approach for carbon emissions (6). This boundary includes the upstream production processes of agricultural inputs (such as fertilizers and pesticides), which account for emissions from the extraction, transportation, and manufacturing of fossil fuels and raw materials. The core production processes consist of two parts: agricultural cultivation processes (including crop sowing, irrigation, fertilization, and harvesting.) and industrial processing processes. Other downstream processes, such as the distribution and consumption of various vegetable oils, are not considered. We estimate the GHG emissions through emission factors and results are expressed in the unit of t $CO_2e$, with global warming potential (GWP) values of 34 kg $CO_2e$ $kg^{-1}$ and 298 kg $CO_2e$ $kg^{-1}$ for $CH_4$ and $N_2O$, respectively (67). Detailed estimates are presented below.

##### 1.3.1.1 GHG emissions during the production and transportation of agricultural inputs

GHG emissions from agricultural inputs calculated in our analysis include seed or nursery, fertilizers, plastic film, pesticides, diesel fuel, and electricity. Quantities of different types of raw materials inputs of each crop are from the published literature based on a meta-analysis approach as described above (Table S2, please see section 1.2.1). Detailed information on input and direct emission factors is presented below (listed in Table S3). Especially, as no systematic life cycle assessment of *Camellia oleifera* oil has been conducted yet, the carbon emission factors for the seed or nursery stages were calculated by referring to the study conducted by Xu et al (2021) (Table S7) (25). For the electricity coefficient of carbon footprint, we collected country-specific electricity emission factors based on alternative data available from the IEA for the power plant and grid model (Table S4) (38).

##### 1.3.1.2 Manure application to crop plantations

Fertilizer application during the crop cultivation stage includes organic and chemical fertilizers. For the chemical fertilizer, we consider four types including N, P,

K, and compound fertilizer shown in Table S3. For the organic fertilizer, we use the manure management data by a global meta-analysis combined with the coefficient of carbon footprint to estimate the manure application to crop plantations.

1.3.1.3 $N_2O$ emissions from nitrogen fertilizer

The direct and indirect $N_2O$ emissions from in-field N fertilizer application were estimated using the developed 5-arc-minute crop-specific estimates of total $N_2O$ emissions from synthetic N fertilizer for the year 2000 and animal manure N application to crop plantations for the year 2010 (30). Animal manure N application was calculated as described above (Please see section 1.3.1.2). For the synthetic N fertilizer inputs, we do not account for N inputs from atmospheric N deposition onto soil, soil mineralization, crop residues, and non-manure organic N additions. We use a global spatial N fertilizer application rates dataset compiled by Potter et al (2010) (68), which estimates crop-specific synthetic fertilizer application rates circa 2000 (1997–2003) at 0.5° resolution in latitude × longitude grid cell.

(1) Direct $N_2O$ emissions

$N_2O$ is generated by the microbially-mediated transformation of N in soils during the process of nitrification and de-nitrification (69), leading to spatial-temporal variation of emission rates that can be modified by diverse vegetative conditions, soil, and climate (70). The most important control factor on soil $N_2O$ fluxes is the reactive N availability, thus the elevated $N_2O$ emissions in a "direct" way are highly associated with the total amount of N fertilizer application (71). In our study, we used a newly developed model (72) to estimate $N_2O$ emissions, which is an updated version of the non-linear "NL-N-RR" model ((NLNRR indicates a non-linear (NL) nitrogen effect (N) random intercept (R) random effect (R) model) by Philibert et al (69). By using random parameters and an exponential model to combine $N_2O$ emissions with N fertilizer application rate, this model performed much better than linear models and exponential models with fixed parameters (73) and was developed from a larger and more recent data set of field experiments compiled by Shcherbak et al (74). The $N_2O$ emission model is based on the following equation (72):

$$N_2O_{\ x,D-N} = \mathrm{EXP}\ (\alpha_0 + \alpha_1 X + \beta Z) + \varepsilon \qquad (1)$$

where $N_2O_{\ x,D-N}$ denotes the direct $N_2O$ emission for crop $x$ (kg $N_2O$-N $ha^{-1}$ $yr^{-1}$); $X$ denotes the applied N dose (kg N $ha^{-1}$ $yr^{-1}$), here, we use the sum of synthetic and manure N fertilizer application dose as model input. We assumed a maximum combined synthetic + manure N application dose of 700 kg N ha$^{-1}$ (72). $Z$ is a binary variable to account for reduced $N_2O$ emissions from flooded rice (74), whose value is equal to "1" if the crop is flooded rice and "0" otherwise. Here, we focus on calculating the vegetable oil crops' direct $N_2O$ emission and thus $Z$=0. β denotes a corresponding "discount factor" parameter, in our analysis, $\beta Z = 0$. The random terms $\alpha_0$, $\alpha_1$, and ε are assumed to be independent and normally distributed, which are presented in equation (1) - (3) based on Philibert et al (69):

$$\alpha_0 \sim N(\mu_0, \sigma_0^2) \quad (2)$$

where $\alpha_0$ denotes the logarithm of location-specific background emission; $\mu_0$ denotes the log mean background emissions; $\sigma_0$ denotes standard deviations showing the variability of $\alpha_0$ across site-years.

$$\alpha_1 \sim N(\mu_1, \sigma_1^2) \quad (3)$$

where $\alpha_1$ denotes the logarithm of location-specific applied N effect; $\mu_1$ denotes the log mean applied N effect; $\sigma_1$ denotes standard deviations showing the variability of $\alpha_1$ across site-years.

$$\varepsilon \sim N(0, \tau^2) \quad (4)$$

where ε denotes the residual error term; $\tau$ denotes the standard deviation of ε. The estimated parameter values of the $N_2O$ emission model are listed in Table S8 (67).

(2) Indirect $N_2O$ emissions

These indirect emissions of $N_2O$ are formed from N volatilization as $NO_X$ and $NH_3$ and redeposition onto water and land surfaces. In addition, in locations where the soil's water holding capacity is less than water input, $NO_3$ may be leached into groundwater and water bodies from managed soils (75). We calculated the indirect $N_2O$ emissions based on the following equations:

$$N_2O_{\ x,ID-N} = \left(F_{x,SN} \times EF_{SN-ATD} + F_{x,ON} \times EF_{ON-ATD}\right) \times 1.0\% + N_2O_{\ x,L-N} \quad (5)$$

where $N_2O_{\ x,ID-N}$ denotes the indirect $N_2O$ emission for crop $x$ (kg $N_2O$-N ha$^{-1}$ yr$^{-1}$); $F_{x,SN}$ represents the annual amount of synthetic fertilizer N applied to soils (kg N ha$^{-1}$ yr$^{-1}$); $EF_{SN-ATD}$ represents a fraction of synthetic fertilizer N that volatilizes as $NH_3$ and $NO_x$, equal to 11% (30); $F_{x,ON}$ represents the amount of manure N applied to soils, and we do not consider the compost and other organic N applied to soils; $EF_{ON-ATD}$ represents a fraction of applied organic N fertilizer materials that volatilizes as $NH_3$ and $NO_x$, equal to 21% (30); 1.0% represents the emission factor for $N_2O$ emissions from atmospheric deposition of N on soils and water surfaces (30).

$$N_2O_{\ x,L-N} = \left(F_{x,SN} + F_{x,ON}\right) \times EF_{L-N} \times 1.1\% \tag{6}$$

where $N_2O_{\ x,L-N}$ represents the $N_2O$ emission from leaching and runoff for crop $x$ (kg $N_2O$-N $ha^{-1}$ $yr^{-1}$); $EF_{L-N}$ represents the fraction of all N added to/mineralized in soils in regions where leaching/runoff occurs, equal to 24% (30); 1.1% represents the emission factor for $N_2O$ emissions from N leaching and runoff (30).

Conversion of $N_2O$–N emissions to $N_2O$ emissions for reporting purposes is performed by using the following equation:

$$N_2O = N_2O\text{–}N \times 44/28 \tag{7}$$

1.3.1.4 GHG emissions from land use change

(1) Land use change carbon stock and burn of current six vegetable oil crops

For the GHG emissions from land use change, we mainly consider the above and below-ground C stock change, forest burning, and organic soil burning, but excluding the leaching, runoff, and induced non-$CO_2$ emissions (76). By defining a specific vegetable oil crop's expansion translates into losses from another land class, we followed the country data adapting from the 'LUC Impact tool' in Poor and Nemecek. (2018) (19) and Blonk Consultants (2014) (77) to show the current (2009-11 avg.) area of six oil crops transformed from forest (Table S9) and then amortized the carbon emissions over 20 years in PAS2050-1 model (Table S10) (19,78). Compared to farm or sub-regional data, country data are more accurately to reflect the drivers of land use change which involves multiple actors (79). Specifically, we excluded the $CO_2$ emissions from direct land use change of forest into cropping systems for groundnut, olives, rapeseed, and sunflower crops, as the total percentage of current crop area transformed from forest are on average less than 1% globally and over two decades. Moreover, the small fraction of forest related to land use change is difficult to address with available maps globally. We then used the FAO data to estimate $CH_4$ and $N_2O$ emissions from forest burning, and $CO_2$, $CH_4$, and $N_2O$ emissions from organic soil burning at the country level for all six vegetable oil crops (Table S11, Table S12) (19,80).

Here, the 'LUC Impact tool' is an Excel tool, which provides a predefined way of calculating greenhouse gas emissions from direct land use change. The tool has three basic functionalities, based on three different approaches related to what data is available for the user.

a) Country known & land use unknown: this approach is described in the PAS 2050-1 published by BSI (78) and is made operational in the tool using various FAO and IPCC data sources. The calculation is based on country-level statistics of the expansion and contraction of forestland, grassland, annual cropland, and perennial cropland in FAOSTAT (80). The land use change of a selected crop is based on country-level statistics on the relative expansion of the selected crop (80).

b) Country & land use unknown: The 'average' land use change GHG emissions for a selected crop are determined by taking the weighted average of all producing countries, based on cultivated area. The GHG emissions for each relevant country are determined through the methodology as described in 'country know & land use unknown'.

c) Country & land use known: In case the country and both the current and the previous land use is known, the carbon stock change of a selected crop is calculated using IPCC defaults and methodology.

The 'LUC Impact tool' has grown in past years based on the PAS 2050 and specifically the PAS 2050-1 frameworks. This basic methodology is now widely referenced in LCA guidelines, such as the Product Environmental Footprint guidelines & Envifood protocol. Due to the data availability, we recognize that total deforestation in the Forest Resources Assessment from remote sensing estimation is higher than an annual inventory of multiple land classes in FAO, which made an underestimates agriculturally induced land use change emissions, but still benefits by reconciling to FAO data (19,81).

(2) Potential Camellia oleifera cultivation on land use change and carbon emission

As a perennial woody plant, *Camellia oleifera* trees have the potential for carbon sequestration through two processes: $CO_2$ absorption during photosynthesis and the fixation of soil organic carbon. The carbon dioxide absorbed by trees can be quantified by measuring their biomass, which is crucial for assessing carbon sinks in artificial tree plantation ecosystems (82). Since *Camellia oleifera* trees are not currently grown on a large scale globally, we used a global-scale Terrestrial Ecosystem Model to quantify the potential impact of growing *Camellia oleifera* on globally suitable land on land use change and carbon emissions.

**Terrestrial Ecosystem Model.** The Terrestrial Ecosystem Model (TEM) is a process-based simulator that estimates carbon and nitrogen fluxes and stocks in terrestrial ecosystems on a monthly time step, using spatial information such as climate, soil, vegetation, and land use (83). In terms of fluxes, net primary production (NPP) represents the biomass produced by ecosystems, commonly used to calculate the harvestable biomass in agricultural ecosystems. net ecosystem production (NEP) represents the net organic carbon content in the ecosystem and serves as a comprehensive indicator of net carbon accumulation. We improved and parameterized the TEM to quantify the carbon dynamics in *Camellia oleifera* ecosystems. Calibration of the TEM was conducted using driving data, and several rate-limiting parameters for biogeochemical processes were obtained through parameterization, including gross primary productivity (GPP), autotrophic respiration, and heterotrophic respiration.

**Impact of Camellia oleifera cultivation on land use change and carbon emission.** We compared the net primary productivity (NPP) of global existing land cover crops with *Camellia oleifera* trees if replaced current vegetable oil crops within suitable areas. The NPP of oilseed crops was estimated based on their corresponding economic yields by using the following equation (84):

$$NPP_i = \frac{\mathrm{EY_i \times D_i \times C \times (RS_i + 1)}}{K \times \mathrm{HI_i}} \quad (8)$$

$$\mathrm{EY_i} = \frac{PY}{\mathrm{f_i}} \quad (9)$$

where $NPP_i$ represents the net primary productivity of *Camellia oleifera* trees. K is the conversion coefficient setting to 100. $EY$ denotes the economic yield calculated based on the predicted crop yield ($PY$) (More details are in Section 3.2). $\mathrm{f_i}$ represents the conversion coefficient for oil production from fresh *Camellia oleifera* fruits, determined as 6.96 (More details are in Section 1.2.4). $\mathrm{HI_i}$ refers to the harvest index, which measures the proportion of above-ground biomass to economic yield. The harvest index for oilseed crops is generally low, ranging from 0.3 to 0.4. Here, we use the mean value of 0.35. $\mathrm{D_i}$ represents the dry weight coefficient of the economic yield ($EY$), which is determined based on the oil content of *Camellia oleifera* seeds, set at 35.65% (More details are in Section 3.3). $C$ represents the carbon content in the biomass of the *Camellia oleifera* tree species, with a commonly used value of 0.5, meaning the conversion from biomass to forest carbon storage (85,86,87). $\mathrm{RS_i}$ indicates the root-to-shoot ratio, representing the ratio of below-ground biomass to above-ground biomass, estimated by the regression model between basal diameter ($D$) and biomass of *Camellia oleifera* tree (Table S13) (88,89,90).

The basal diameter ($D$) of *Camellia oleifera* trees typically ranges from 3.7 to 15.2 cm and varies depending on the tree's age and growth rate. Here, we determined the total biomass of the *Camellia oleifera* plantation based on Lin et al (2014) (91). The total biomass was estimated to be 35.76 t • $hm^{-2}$, with the stem biomass being the highest at 12.16 t • $hm^{-2}$, accounting for 34.00% of the total biomass. The order of organ biomass from largest to smallest is as follows: trunk > branch > root > leaf > fruit, and the root biomass was calculated to be 7.80 t • $hm^{-2}$. The root-to-shoot ratio $RS_i$ was determined as 0.2712 by using the formula $RS_i$ = 7.80 / (35.76-7.80).

The land suitability assessment for *Camellia oleifera* shows that the most suitable areas are in forests, wastelands, and bare lands. It means that the carbon storage of the ecosystem will be altered after being formed by replacing or expanding the *Camellia oleifera* plantation. We calculated the difference in net ecosystem productivity ($NEP_\Delta$, t/$hm^2$) caused by replacing from *Camellia oleifera* plantation in the existing terrestrial ecosystem. The formula for calculating $NEP_\Delta$ is as follows:

$$NEP_\Delta = NEP_o - \left(NPP_i - R_{h,i}\right) \tag{10}$$

$$R_{h,i} = \mathrm{Average}(\mathrm{S1{:}S1958}) \tag{11}$$

where $NEP_\Delta > 0$ represents $CO_2$ emissions from the terrestrial ecosystem to the atmosphere after replacing from Camellia plantation, while the opposite indicates $CO_2$ absorption. $NEP_o$ represents the net ecosystem productivity of the existing terrestrial ecosystems such as forests, wastelands, and bare lands. $NEP_o$ indicates the difference between net primary productivity (NPP) and heterotrophic respiration ($R_h$). When $NEP_o$ is greater than 0, it signifies that the ecosystem acts as a $CO_2$ sink, indicating $CO_2$ absorption by the ecosystem from the atmosphere. Conversely, if $NEP_o$ is negative, it represents a $CO_2$ source, indicating $CO_2$ emissions from the ecosystem to the atmosphere. The global daily Net ecosystem productivity dataset (2017-2019) was generated using a Boreal Ecosystem Productivity Simulator model with vegetation parameters (leaf area index, land cover type, and clumping index), remote sensing data, meteorological data, and atmospheric $CO_2$ concentration (92). The dataset covers all vegetated land areas globally with a spatial resolution of 0.072727° × 0.072727°, which was interpolated to 0.038° × 0.038° using bilinear interpolation (93). $R_{h,i}$ represents the average soil heterotrophic respiration for 1,958 *Camellia oleifera* sample sites worldwide (Figure S6), which was obtained from GBIF and extracted by using a point query analysis to determine the soil heterotrophic respiration.

1.3.1.5 GHG emissions during vegetable oil processing phase

For the current six types of vegetable oil processing information, we directly collected the mean value of greenhouse gas emissions per kilogram and combined the oil density to make it comparison in the unit of $kgCO_2eq$ per liter, showing a descending order with palm oil (1.05) > olive oil (1.28) > soybean oil (0.30) > sunflower oil (0.20) > rapeseed oil (0.19) > groundnut oil (0.84) (Table S14) (8,54-60,63,94). For the *Camellia oleifera* oil, we mainly considered the GHG emissions from electricity, sodium hydroxide, phosphoric acid, activated carbon, and steam used in the oil refining phase (More details are in Section 1.2.4).

#### 1.3.2 Water use

Refer to "The Water Footprint Manual" (95), we consider the crop green, blue, and gray water footprint in the crop plantation stage. Here, the green water footprint represents the amount of rainwater and soil moisture consumed by crops through evaporation or transpiration without contributing to surface runoff. It reflects the utilization of natural water resources that do not directly affect water scarcity. The blue water footprint accounts for the surface water and groundwater consumed during the growth of crops. This component quantifies the water resources directly withdrawn from rivers, lakes, reservoirs, or aquifers for irrigation purposes. The gray water footprint is associated with the water consumption required to dilute pollutants present in the crops to reach the natural background concentration or meet environmental water quality standards. This component takes into consideration the potential pollution caused by agricultural activities. In particular, we do not consider water use outside the production phase of the crop life cycle because the amount is small (34,96). The calculation process is as follows:

$$WF_{proc} = (WF_{green} + WF_{blue} + WF_{gray})/Y \quad (12)$$

where $WF_{proc}$ is the total water footprint per unit of yield during crop growth ($m^3/t$); $WF_{green}$, $WF_{blue}$, and $WF_{gray}$ denote the green, blue, and grey water footprints of the crop ($m^3/yr$), respectively; and Y is the total crop yield (t). Specifically, crop blue water footprint is the amount of evapotranspiration used for field irrigation (95). Therefore, we calculated the consumptive water use of rainfed and irrigated crops separately, where rainfed agriculture has a water footprint of green and grey, while irrigated agriculture has a water footprint of three types of water. agriculture has a water footprint of green, blue, and grey.

1.3.2.1 $WF_{green}$ and $WF_{blue}$

The green and blue water footprint of the crop is equal to the value of the green and blue water consumption of the crop multiplied by the crop total harvested areas, as follows:

$$WF_{green} = CWU_{green} \times A \tag{13}$$

$$WF_{blue} = CWU_{blue} \times A \tag{14}$$

$$CWU_{green} = 10\sum_{d=1}^{lgp} ET_{green} \tag{15}$$

$$CWU_{blue} = 10\sum_{d=1}^{lgp} ET_{blue} \tag{16}$$

where $CWU_{green}$ and $CWU_{blue}$ is the crop green water consumption and blue water consumption ($m^3$); $A$ represents the harvested area ($hm^2$). The coefficient of 10 is used to convert water depth (mm) into the amount of water ($m^3 \cdot hm^{-2}$) per unit land area. $lgp$ represents the length of the growing period in days.

The length of growing periods varies greatly among different crops and has a significant impact on crop water consumption. As long-term perennial crops, such as *Camellia oleifera*, palm, and olive trees, transpire throughout the year. Therefore, to calculate the variation in transpiration of perennial crops over the entire growing cycle, we should estimate the annual average transpiration over its entire life cycle. *Camellia oleifera* have a long lifespan, typically reaching 70-80 years and providing yields from the fifth to sixth year onwards (97). Here, we use the water consumption over 80 years divided by the total yield over 75 years to obtain the water footprint per unit of yield. For palm trees, the crop water requirement of oil palm is estimated based on soil water readily available over 25 years; 5 years for the starting period and 20 years for an established period (98); For olives, the growing phase, from the 1st to the 4th years and the production phase from the 5th to the 50th years (99). Given the availability of long-term crop yield data, we calculate the total green water and blue water consumption for perennial crops by multiplying them directly with the ratio coefficient of their production period to the growing period. Specifically, we use the coefficients of 80/75, 25/20, and 50/46 for these crops of *Camellia oleifera*, palm, and olive trees, respectively.

$ET_{green}$ and $ET_{blue}$ refer to crop green water evapotranspiration and blue water evapotranspiration (mm/yr), respectively, and are calculated as follows:

$$ET_{green} = \min\,(ET_c, P_{eff}) \tag{17}$$

$$ET_{blue} = \max(0, ET_c - P_{eff}) \tag{18}$$

where $ET_c$ is the crop's actual evapotranspiration (mm/yr); $P_{eff}$ is the effective precipitation available to crops (mm/yr).

$$P_{eff} = \begin{cases} \frac{P(125-0.6P)}{125} & (P < 250/3mm/yr) \\ 125/3 + 0.1 \times P & (P \geq 250/3mm/yr) \end{cases} \tag{19}$$

where $P$ (mm/yr) represents the total precipitation. The crop's actual evapotranspiration ($ET_c$) during the growth period is calculated based on the reference crop potential evapotranspiration ($ET_0$) and the crop coefficient ($K_c$) (Table S15, see Section 3.2.3) (35).

1.3.2.2 $WF_{gray}$

The grey water footprint of the crop ($WF_{gray}$) was calculated as follows:

$$WF_{gray} = [(\alpha \times AR)/(C_{max} - C_{nat})] \times A \tag{20}$$

where $\alpha$ represents the leaching rate, which is the proportion of the pollution entering the water to the total amount of chemical substances applied. As water can simultaneously dilute multiple pollutants, the water requirement for diluting pollutants is determined by the pollutant with the largest required dilution water volume. Considering the availability and representativeness of data, the water requirement for diluting leached nitrogen is used as a characterization. The leaching nitrogen is typically 5% to 15% of the nitrogen fertilizer application, and here, $\alpha$ is set to 10% (100). $AR$ represents the total amount of fertilizer applied per hectare of land (kg/hm$^2$); $C_{max}$ is the maximum allowable concentration of pollutants to achieve the corresponding water quality standards, with the dilution standard set at a nitrogen concentration of no more than 10 mg per liter of drinking water, $C_{max}$= 0.01 kg/m$^3$; $C_{nat}$ represents the natural background concentration of pollutants, with the natural background concentration of the receiving water body determined in an uncontaminated environment as $C_{nat}$=0 kg/m$^3$.

#### 1.3.3 Land use

In terms of land use for each vegetable oil crop, we mainly consider the stage of crop production, since an insignificant amount of land use is consumed outside the production phase of the crop life cycle (34). Inverse yield and crop occupation time were used to calculate the environmental impact of land use.

Fallow requirements increase occupation time, but multiple cropping or intercropping decreases it. The minimum fallow requirements for crops were determined by dividing the maximum monthly growing area (MMGA) by the cropland extent (CEMIRCA) (101,102). MMGA represents the maximum area needed for crop rotation at a specific location. CEMIRCA is the area where arable crops, including fallow, are cultivated (103). The fallow requirements were determined by considering the commercial lifespans and literature data, taking into account the variations in the lifecycles of permanent crops and orchard crops (More details are in Poor and Nemecek. (2018) (19)).

Multiple cropping was determined by calculating the ratio of the maximum monthly growing area (MMGA) to the area harvested (AH) (102). AH represents the number of times a crop is harvested in a year (e.g., for double cropping, MMGA/AH = 2). Land use, calculated based on yield, can be adjusted by multiplying it with these ratios to align with the global cropland extent from FAOSTAT, as shown in Equation (21) and Equation (22). Therefore, the land use for each vegetable oil crop includes land occupation for seed or nursery production, arable land for crop cultivation, and land utilization for fallow requirements specific to each region.

Land use for producing 1 kg of vegetable oil product is mainly decided by the cropping duration of different vegetable oil crops. Thus, we divided the vegetable oil crops into two main types, namely temporary crops (groundnut, rapeseed, soybean, and sunflower) and permanent crops or orchard crops (*Camellia oleifera*, palm, and olive). After getting the land use to produce 1 kg of product, we then multiply with the weight of fresh fruits required to produce one liter of vegetable oil for different crops to get the total land use per liter of oil.

For temporary vegetable oil crops, land use was calculated as follows:

$$Land\ use = \frac{10{,}000}{yield} \times \frac{Seed+yield}{yield} \times \frac{Crop\ duation}{365} \times \frac{Rotation\ duation}{Cultivated\ duation} \tag{21}$$

where $yield$ and $Seed$ are expressed in kg ha$^{-1}$ and are based on the same marketable weight basis, where a 15% moisture post-field loss is taken into account. The $duation$ is measured in days. $Land\ use$ is defined as the area occupied to produce 1 kg of product, in m$^2$·year.

For permanent or orchard vegetable oil crops, land use was calculated as follows:

$$Land\ use = \frac{10{,}000}{yield} \times \frac{Cultivated\ duation}{Bearing\ duation} \times Nursery \times \frac{Rotation\ duation}{Cultivated\ duation} \quad (22)$$

$$Nursery = 1 + \frac{Nursery\ duation/365}{Sampling\ yield} \times \frac{Orchard\ density}{Cultivated\ duation} \quad (23)$$

where $yield$ refers to the period when the orchard bears marketed fruit ($Bearing\ duation$), while $Cultivated\ duation$ represents the period from orchard establishment to removal. The non-bearing period after establishment is the difference between $Bearing\ duation$ and $Cultivated\ duation$. Here, we assumed 5, 4, and 3 years for *Camellia oleifera*, palm, and olive, respectively. The fallow period after orchard removal and before replanting is $Rotation\ duation / Cultivated\ duation$. The nursery period is more important than the seed for the permanent or orchard vegetable oil crops. $Nursery\ duation$ is the time from planting seedlings to the nursery trees (in days); $Sampling\ yield$ is the number of marketable saplings produced per hectare per year; and $Orchard\ density$ is the number of trees required for 1 ha land.

**Section 2: Global land suitability assessment of *Camellia oleifera* cultivation**

*Camellia oleifera* plantations in China have been increasing continuously to support the high demand for producing eatery products and daily necessities (104), even expanding into some areas that were never grown before due to the non-suitable climate conditions (105,106). However, the expansion of crop cultivations, including for *Camellia oleifera*, has direct impacts on land use potential that also affects the effective yield. In the case of China, land suitability assessment of *Camellia oleifera* has been studied but usually at specific sites or focused on a single parameter such as soil pH values, precipitation, temperature, etc., due to data limitation and low-resolution dataset (107,108,109,110). The selection of suitable land for specific crops can decrease the environmental impacts, especially for land use, and increase the potential yield. Hence, this study aims to assess the suitability for global Camellia oleifera plantations by incorporating with GIS-based suitability procedures and machine learning methods. We propose to construct an indicator system for assessing the suitability of *Camellia oleifera* plantations by considering three major categories of natural environmental factors, namely meteorological conditions, soil conditions, and topography. This indicator system comprises a total of 11 factors, which were derived from long-term time series data spanning from 2010 to 2020. To enhance the accuracy of the evaluation model, we incorporated global ecological conservation policies and adjusted the model based on land resource types favorable for *Camellia oleifera* cultivation. The refined evaluation model enables fine-scale zoning and grading of the global suitability for *Camellia oleifera* cultivation, thereby providing a scientific basis for the introduction and expansion of *Camellia oleifera* and the planning layout of global industries. The findings could contribute to the development of evidence-based strategies for *Camellia oleifera* cultivation and global vegetable oil-related industry planning. Furthermore, expanding the *Camellia oleifera* plantations into diversification of vegetable oil production systems and agricultural systems, in general, can provide important buffers for extreme climate and unexpected geopolitical events. As such, we urge policy makers to seize opportunities to expand diversified vegetable oil production system and advance on the agenda of a sustainable food systems transformation based on diversified systems.

### 2.1 Land suitability indicators selection

The growth of *Camellia oleifera* exhibits zonal patterns, which are influenced by the natural attributes of land suitability and environmental factors such as climate conditions, geological features, and soil characteristics. In this study, we propose an indicator system for evaluating the suitability of *Camellia oleifera* cultivation based on the principles of stability, dominance, and comprehensiveness in indicator selection. The indicator system, as presented in Table S16, encompasses three key aspects: climatic conditions, topographic features, and soil characteristics, all of which play significant roles in shaping the favorable environment for *Camellia oleifera* growth. By integrating these factors, we aim to provide a comprehensive understanding of the natural attributes that contribute to the suitability of *Camellia oleifera* cultivation.

(1) Climatic conditions.

*Camellia oleifera* prefers warm and humid climates, with temperature, frost-free period, and annual rainfall as factors that directly affect the flowering period, fruit setting, and fruit plumpness. Among them, temperature is the key factor affecting the flowering period and fruit-set rate, and unsuitable temperatures can affect insect pollination and flower opening. The base temperature, which refers to the minimum, optimum, and maximum temperatures that allow plant growth, is the dominant factor affecting the whole life cycle growth and development of *Camellia oleifera* (111,112). Low temperatures not only affect the growth and development of *Camellia oleifera* but also have a decisive effect on its flowering period and oil accumulation.

① When the annual average temperature is below 13°C, it will affect the normal flowering and reduce the fruit set. ② High temperatures in January help to reduce frost damage to young fruits and increase the yield. When the temperature reaches above 10°C, it is beneficial for the swelling of the ovary and the formation of young fruits. Once extreme temperatures occur (below -10°C), it will directly affect the growth of *Camellia oleifera*. ③ October to December is the flowering period of *Camellia oleifera*, and the average temperature between 14-18°C is most suitable. If the temperature is too low, it will cause flowers and fruit to drop. ④ Whether heat can be fully utilized also depends on the length of the frost-free period. July to September is the period when oil accumulates in the seeds of *Camellia oleifera*. Generally, the frost-free period required by *Camellia oleifera* should be more than 200 days, but not less than 150 days, to ensure that young fruits are not damaged by frost and safely overwinter. ⑤ In addition, although *Camellia oleifera* is drought-resistant, continuous drought will reduce the survival rate of seedlings and cannot ensure the production of *Camellia oleifera* seedlings (113). *Camellia oleifera* trees generally require more than 600 mm of rainfall, and no enough rainfall will affect the growth of fruit trees. During the flowering period, the rainfall should not be too high, otherwise it will affect pollination (114). Therefore, following the impact of *Camellia oleifera* habitat conditions and bioclimatic factors, we selected five dominant climate factors: Annual average temperature (℃), Average temperature in January (℃), October to December Average temperature (℃), Frost-free days (d), and Average annual precipitation (mm).

(2) Topographic features.

Topographic factors such as elevation, slope, and aspect influence the growth and development of *Camellia oleifera* by affecting light conditions and soil nutrient status. In terms of fruit characteristics, elevation also plays an important role, as lower elevations tend to have larger fruits with higher seed and kernel extraction rates (115,116). Slope gradient directly affects soil conservation and fertility, with slopes less than 30° being suitable for cultivating *Camellia oleifera* in the forest, while mountainous areas and valleys should have a width greater than 50 meters (117). Different aspects will have an impact on the light exposure of *Camellia oleifera*. During critical physiological stages such as flowering, low temperatures, and insufficient light can affect the fruit setting rate. Optimal aspects for sufficient sunlight are east-facing slopes (67.5° to 112.5°), southeast-facing slopes (112.5° to 157.5°), south-facing slopes (157.5° to 202.5°), and southwest-facing slopes (202.5° to 247.5°) (118).

(3) Soil characteristics.

Land attributes and their composition, especially soil, serve as the carrier of mineral nutrients and water necessary for crop growth. Under favorable climatic conditions, the natural production potential of crops can be realized by overcoming land limitations. For *Camellia oleifera* cultivation, the soil has two main impacts: soil acidity and soil fertility. *Camellia oleifera* is an acid-loving species, and its root system has a buffering capacity for the acidity of sap. This is because the root cells of *Camellia oleifera* lack slightly alkaline phosphates and can only buffer with organic salts that are slightly acidic. Additionally, afforestation of *Camellia oleifera* on soils with a total nitrogen content greater than 0.08% and total organic carbon content greater than 0.75% is beneficial for fruit development and oil content in seeds (119).

The data sources for the 11 indicators required for assessing the suitability of *Camellia oleifera* are as follows: (1) Meteorological data is obtained from the UK National Centre for Atmospheric Science (NCAS) using the CRU TS climate dataset, which provides monthly meteorological data covering land surfaces from 2011 to 2020 (120). (2) Elevation, slope, and aspect terrain factors are extracted from the Digital

Elevation Model (DEM) data (https://globalmaps.github.io/el.html). (3) Soil data is collected from the Harmonized World Soil Database (HWSD) constructed by the Food and Agriculture Organization (FAO) and the International Institute for Applied Systems Analysis (IIASA). The classification attributes are obtained by linking with HWSD_SMU, and the soil classification system used is FAO-90 in a WGS84 projection coordinate system. In particular, all datasets have a 0.083° resolution except the CRU TS climate dataset, which has a 0.5° resolution. For consistency, the CRU TS climate dataset was resampled to 0.083° using bilinear interpolation.

Based on spatial analysis techniques in ArcGIS 10.6, a single-factor suitability evaluation was conducted to visually express the suitability level zoning of each grid unit. There are spatial variations in the evaluation of suitable zones for *Camellia oleifera*, especially for three temperature indicators, a. Annual average temperature/°C, b. The average temperature in January/°C, and c. October to December average temperature/°C, which are mainly located in South America, Africa, Oceania, as well as the southern parts of Asia and North America (Figure S7). The suitable zone for indicator e. Average annual precipitation/mm is found in the southeastern parts of North America, central parts of Africa, southern and southeastern parts of Asia, but only along the eastern and northern coastal regions of Australia. The single-factor indicators f, g, and k do not exhibit significant restrictions on *Camellia oleifera* cultivation globally, as most regions fall within the suitable cultivation zone. Indicators i and j show similar spatial patterns of suitability, with very suitable regions appearing in northern Asia, most parts of Europe, northern and southern Africa, central Asia, and western Australia. However, large areas of unsuitable regions are observed in central Asia and western Australia. It should be noted that the single-factor evaluation model only considers the effect of a certain factor under the condition that other influencing factors remain relatively constant, and thus cannot fully reflect the actual impact of climate factors on crop growth (121). Therefore, to better reflect the habitat characteristics of *Camellia oleifera* and its suitability on a global scale, a comprehensive evaluation should be conducted by combining different indicator weights based on the single-factor evaluation.

### 2.2 Weight identification

When collecting these 11 indicators among climatic, topographic, and soil conditions from published papers, we noticed that there are discrepancies in the weightings assigned to these indicators. To address this issue, we employed machine learning methods to determine the most appropriate weighting for each indicator. This approach was selected because it allows for the consideration of multiple factors and captures complex relationships between variables, which may not be apparent through traditional statistical analysis. Additionally, machine learning algorithms can be used to optimize the weighting scheme based on a given set of criteria, such as accuracy or precision, to improve model performance and the ability to adapt under changing conditions. In particular, we chose to use the machine learning method only to determine indicator weights instead of the geospatial suitability mapping to avoid overfitting caused by the model performing well on training data, but poorly on unknown data (22,123).

**Data collection and processing.** We first collected 29,641 station data from the National Oceanic and Atmospheric Administration (NOAA) with detailed geographic information (latitude and longitude). By following the National Development Plan for the Oil Tea Industry (2009-2020) issued in China, we classified counties suitable for *Camellia oleifera* cultivation into three levels: highly suitable cultivation area, moderately suitable cultivation area, and suitable cultivation area. Based on the geographic information of each station, we then further determined their corresponding suitability levels for *Camellia oleifera* cultivation. In addition, we marked the sample points collected from NOAA that showed no production in China during the past decade (2011-2021) as unsuitable cultivation areas. For each suitable level, we marked the point samples with labels 3, 2, 1, and 0 to represent a highly suitable cultivation area, moderately suitable cultivation area, suitable cultivation area, and not suitable cultivation area, respectively. Subsequently, by utilizing the "extract values to points" function with the latitude and longitude information in Arcgis, we obtained detailed values for 11 indicator layers for *Camellia oleifera* cultivation. Finally, after removing data points without suitability levels, duplicate points, and missing data entries, we retained a total of 1,263 records for weight identification in machine learning methods (Table S17).

**Indicators correlation.** After standardizing the 11 indicators, we examined the intercorrelations among them, using a range of values from -1 to 1 to visualize the relationships between the indicators to gain a comprehensive understanding of their associations. A correlation coefficient of 1 indicates a positive correlation, -1 indicates a negative correlation, and 0 indicates no correlation (Figure S8). Although the five climate indicators appear to be highly correlated, we still chose to include all of them in our land suitability assessment instead of selecting just one. Each indicator reflecting a distinct aspect of the local climate conditions can affect the suitability of *Camellia oleifera* cultivation. For example, temperature and precipitation are both critical factors for plant growth and development, but they can have different effects on *Camellia oleifera* production depending on their timing and intensity. Furthermore, including multiple indicators allows us to identify potential interactions or feedback loops that may not be evident when examining individual factors in isolation. For instance, high temperatures may increase water demand and exacerbate water stress in arid regions, which in turn can affect soil fertility and nutrient availability.

**Machine learning methods comparison.** We selected four different machine learning methods, namely multinomial logistic regression, support vector machine, random forest, and decision tree, to compare their performance in weight identification for predicting the suitability of *Camellia oleifera* cultivation. Here, multinomial logistic regression is a widely used statistical model that allows us to predict the probability of an observation belonging to multiple classes (124). It is suitable for our analysis as it can handle categorical outcomes. Support vector machine is a powerful algorithm that aims to find the best hyperplane to separate data points into different classes with a high degree of accuracy (125). Random forest is an ensemble learning method that combines multiple decision trees to make predictions. It is robust against overfitting and can handle both categorical and continuous variables effectively (126). The random feature selection and bootstrap aggregating techniques employed by random forest enhance its prediction accuracy. By using a tree-like model to make decisions based on multiple features, a decision tree is easy to understand and visualize, making it useful for identifying the most influential factors in predicting *Camellia oleifera* cultivation suitability (127). By comparing these four machine learning methods, we aim to evaluate their performance on weight identification in terms of prediction accuracy, computational efficiency, interpretability, and robustness. This comprehensive analysis will enable us to determine the most suitable approach for predicting *Camellia oleifera* cultivation suitability and provide valuable insights for decision-making in the agricultural sector.

Here, the dataset with a total of 1,263 records is divided into a 0.7 training set and a 0.3 testing set to effectively utilize the data for model training and evaluation. By allocating the majority of the data for training, we ensure that the model can learn the patterns and features of the data sufficiently. Simultaneously, reserving a portion of the data as a testing set allows us to assess the model's performance on unseen data, evaluating its generalization ability and prediction accuracy. This approach of splitting the dataset into training and testing sets helps prevent overfitting on the training set and enables an objective evaluation of the model's performance, providing a scientific basis for model selection and optimization (128).

By comparing the accuracy, precision, and recall of the four models on both train and test datasets, the random forest outperformed the multinomial logistic regression, support vector machine, and decision tree models (Figure S9). The high $R^2$ score of random forest ($R^2 = 0.914$) indicates its superior ability to explain the variance in the data and capture the underlying relationships between the input features and the *Camellia oleifera* cultivation suitability more effectively. In terms of accuracy, the random forest achieved the highest value among the four models. This means that it accurately predicted the oil tea cultivation suitability for a larger proportion of the samples in the testing set. The precision measures the proportion of correctly predicted positive cases among all predicted positive cases, while recall measures the proportion of correctly predicted positive cases among all actual positive cases. The high accuracy suggests that the Random Forest model provides reliable predictions for practical applications. Random Forest model exhibited higher precision and recall values compared to the other models, indicating its ability to correctly identify and classify *Camellia oleifera* cultivation suitability. The weight identified by random forest for each indicator shows a descending order with Average temperature in January/°C (0.16074526) > Average annual precipitation/mm (0.15297809) > Frost-free days/d (0.14447819) > October to December Average temperature /°C (0.14067698) > Annual average temperature /°C (0.1173547) > Soil pH (0.08795861) > Elevation/m (0.05662247) > Organic carbon % of weight (0.04990066) > Total N % of weight (0.04083468) > Slope/° (0.03195019) > Aspect (0.01650018). Then, the spatial analysis and map algebra modules were employed to perform a weighted overlay of individual indicator raster data, resulting in a preliminary comprehensive evaluation of *Camellia oleifera* cultivation suitability.

### 2.3 Incorporation of land use constraints

Using a combination of five datasets, namely global land use/cover, World Protected Areas Database, GRIP global road network, intact forest areas, and key biodiversity zones, we conducted spatial exclusion and refinement processes (Figure S10) to generate a suitability zoning map for global *Camellia oleifera* cultivation. This incorporation of land use constraints ensures the feasibility of *Camellia oleifera* cultivation worldwide.

Firstly, we utilized global 300m land use/cover data to exclude unsuitable regions such as croplands and urban areas, focusing only on potential areas for *Camellia oleifera* cultivation, such as sparse forests, shrublands, and non-forested lands (129). Secondly, the latest global terrestrial and inland protected areas data (as of November 2022) were obtained from the World Database on Protected Areas (WDPA), jointly published by the United Nations Environment Program (UNEP) and the International Union for Conservation of Nature (IUCN) (130). Thirdly, artificial impervious surfaces, particularly roads such as highways, major roads, and ferries, were considered detrimental to surface water infiltration and natural evapotranspiration processes. These factors can negatively impact crop growth, leading to increased heat and aridity around trees due to the presence of impermeable surfaces or road cover. Global road data was sourced from the Global Roads Inventory Project (GRIP) (www.globio.info), which compiled road information covering more than 21 million kilometers across 222 countries/regions (131). Lastly, the intact forest landscape is an unbroken expanse of natural ecosystems within the zone of current forest extent, showing no signs of significant human activity and large enough that all native biodiversity, including viable populations of wide-ranging species, could be maintained (132). Excluding these areas within the intact forest landscape and key biodiversity zones from *Camellia oleifera* cultivation ensures their preservation and minimizes the potential negative impacts on biodiversity and ecosystem services (133). By considering these five indicators in the spatial exclusion process, we finally generate a suitability zoning map that promotes sustainable *Camellia oleifera* cultivation practices globally.

### 2.4 Field verification

Field verification is crucial in the evaluation and validation of research findings and models, especially in the case of assessing the suitability and distribution of *Camellia oleifera* cultivation areas. While geospatial analysis provides valuable predictions on the spatial pattern, on-the-ground verification ensures the accuracy and reliability of our results. This further provides a basis for delineating the potential advantage areas for global *Camellia oleifera* production and formulating related industry policies.

To obtain the global distribution data of existing *Camellia oleifera* cultivation areas, we searched for the scientific name of *Camellia oleifera* in The Global Biodiversity Information Facility (https://www.gbif.org/) and the Chinese Virtual Herbarium (https://www.cvh.ac.cn/), excluding duplicate records (the same specimen in different herbariums and duplicate sampling at the same location and time), records without clear collection times or geographic location information (134,135). By using the georeferencing method of place name reversal to obtain and verify the latitude and longitude information of each record through Geosharp2.1 and Google Map, a total of 1,958 specimen data were finally screened (Table S18, Figure S6).

Our land suitability map shows that 70.9% of the area is distributed in subtropical and tropical regions, which are traditionally considered suitable habitats for *Camellia oleifera* cultivation (136). It is generally believed that there is no big difference in climate, soil conditions, etc., suitable for plant growth within the provincial scale. The field cross-validated result indicates that 88.20% of the existing *Camellia oleifera* cultivation samples are distributed within the range of our suitability assessment results or their corresponding provincial level range, demonstrating the high accuracy of our land suitability evaluation results. For the rest 11.80% of samples that are not within the range, upon verification, they are mostly distributed in some islands, such as the South China Sea Islands of China, Miyako Island, and Ishigaki Island of Japan. This discrepancy is caused by the lack of climate and topography data, as it did not cover these regions, only covering the vast majority of inland areas globally.

### Section 3: Projection of *Camellia oleifera* production capacity

Crop production increase depends on either an expansion in planting areas or an improvement in crop yield per unit area. Firstly, planting area refers to the actual land area where crops are planted, and the increase in planting area is mainly achieved by reducing fallow land, shortening the fallow period, increasing the replanting index, and cultivating more suitable uncultivated land. The main factors affecting crop yield per unit area include variety improvement and increased agricultural inputs such as fertilization and irrigation, which can be divided into three aspects: ① Breeding of superior varieties provides strong support for food security from the source (137). With the realization of CRISPR-Cas9 gene editing technology, newly engineered domesticated varieties can be directly used in cultivation, or can be hybridized with excellent strains to quickly introduce new traits without delay caused by wild germplasm utilization. ② Soil fertility is the ability of the soil to provide enough nutrients required for crop growth. Appropriate application of chemical fertilizers can increase crop yield and now the contribution rate of fertilizer to global grain yield is 50% to 60% (138,139). However, there is a decreasing marginal benefit of agricultural fertilizer use, and the contribution rate to yield increase decreases as the amount applied increases (140). ③ Agricultural irrigation can ensure stable and high agricultural yields by meeting the water requirements of crops. In addition to supplying water for crops, it can also regulate soil temperature, humidity, soil air, and nutrients, or even improve soil fertility and flush out saline-alkali soils (141). Under the improvement measures such as breeding of superior crop varieties, fertilization, and irrigation, the climatic potential yield for crops now is achievable. In addition, agroforestry are sensitive to climate change, and analyzing the impact of climate change on potential yield and simulating the corresponding spatiotemporal characteristics can be used to inform decision-making regarding land use, resource allocation, and agricultural policies, with the ultimate goal of ensuring food security and sustainable food production (142,143). The role of simulating the potential yield for crop production is to provide insights into how various factors, such as climate change, agricultural practices, and technological advancements, may impact crop yields in the future.

#### 3.1 Research progress of crop potential yield modelling

Crop potential yield refers to the maximum yield that can be obtained from a given crop under specific environmental and management conditions. It represents the upper limit of possible crop productivity, which is influenced by multiple factors such as soil fertility, climate, water availability, and agricultural practices. For the crop potential yield modelling, there are currently four main estimation methods as follows (144):

(1) Field trial yield method refers to obtaining crop yields through optimizing agricultural techniques and management measures in experimental fields selected based on representative agricultural cultivation conditions in a specific region (145). Since field experiments are conducted under natural soil and climatic conditions similar to those in actual field production, the results are usually considered directly applicable for demonstration and promotion in similar production conditions. However, due to differences in agricultural environments and inputs such as light, temperature, water, soil, and atmosphere across different regions of the world, the crop yields obtained through this method also vary, leading to limitations in reflecting the overall yield potential.

(2) The regional high-yield record value method refers to the record-high crop yields achieved by farmers under the guidance of experts through the implementation of various measures to enhance agricultural production, including the use of superior crop varieties and high-yield cultivation techniques developed by researchers. Compared to other methods, these record yields are closer to the maximum yield potential of the crops. However, these record yields can only represent the current highest yields in the region and cannot reflect the yield levels in other areas, as the regionally differentiated oil, climate, and other planting conditions (146).

(3) The High-yield farmer survey method involves extensive surveys and collection of crop yield data from different regions, climate types, and crop varieties. The potential values of crop yields in actual field conditions are determined based on the relatively high or highest yields observed among a large number of farmers (typically selecting the top 5% to 10% of yields) (147). However, due to constraints imposed by technological conditions such as crop variety improvement and mechanical harvesting, or limited input costs during actual cultivation by farmers, the yields obtained through the high-yield farmer survey method are usually lower than those obtained through the regional record yield method and crop production potential model simulations.

(4) Crop simulation model refers to combining agricultural meteorological data, crop cultivation management experience (such as crop varieties, sowing time, growing period, planting density, etc.), and other information, to dynamically simulate crop

growth and development and quantitatively estimate crop yield potential (148). This method can effectively consider the interaction among crop growth and development, agricultural climate and environmental changes, and cultivation management measures, and it can be used at the field, regional, national, and even global levels to help understand and predict crop potential yield. It is currently the most commonly used method for quantitatively analyzing food potential yield (149). However, due to the diversity of crop varieties and differences in field practices and planting measures, the choice of models, parameter settings, and scenario simulations can affect the results of crop potential yield. Currently, globally mature crop potential yield models can be classified into four categories.

① Empirical formula models, also known as comprehensive climate factor models, estimate the crop potential yield by considering empirical climate indicators' impacts and establishing correlations between crop yield data and meteorological observations based on historical statistical data. The empirical formula models mainly include the Miami model, the Gessner-Lieth model, and the Chikugo model. The Miami model, developed by German ecologist H. Lieth in 1972, was the first global primary productivity model for ecosystems (150). The reliability of the results estimated using this model ranges from 66% to 75%. However, these empirical formula models consider only a limited number of factors and are suitable for rough estimates of crop yields on a large scale.

② Crop growth process simulation models can simulate the entire process of crop sowing, germination, growth, and harvesting by integrating the physiological characteristics of crops, photosynthesis during growth, and external environmental conditions. Currently, crop growth process simulation models have been applied to various types of crops such as wheat, potatoes, and soybeans, and which are mainly including the Crop Environment Resource Synthesis model (151), Simple and Universal Crop growth Simulation model (152), Environmental Policy Integrated Climate model (153), and the GROPGRO model (154). These models have the advantage of considering multiple factors that influence crop yield and can accurately estimate specific crop yields in a particular region. However, they require high reliability in parameter acquisition, as inaccurate parameters can significantly reduce the reliability of the model's final predictions.

③ Machine Learning prediction method, a subset of Artificial Intelligence that focuses on learning, is a practical technique that may give superior yield prediction based on a variety of characteristics. Machine learning can identify patterns and correlations in datasets and uncover information. However, machine learning trained using datasets that describe the outcomes highly relied on previous experience. To tackle the issue at hand, it is essential to choose the appropriate algorithms, and both the algorithms and the supporting platforms must be able to handle the big volume of data (155).

④ Mechanistic models, also known as environmental factor stepwise correction models or potential attenuation models, study the attenuation effect of external environmental factors on crop productivity based on a stepwise correction approach. Specifically, starting from the light interception characteristics and photosynthetic efficiency of crops, these models calculate the photosynthetic potential yield, light and temperature potential yield, climatic potential yield, and soil potential yield to finally get the potential comprehensive production capacity of crops by stepwise "attenuation" calculation based on the principles of energy conversion and crop growth mechanism in terms of light, temperature, water, soil, and other factors. These models are recognized worldwide as the best method for simulating macro-scale cereal potential yield, such as the Wageningen model and the Global Agro-Ecological Zones (GAEZ) method. The GAEZ model is an agricultural ecological zoning model jointly developed by the Food and Agriculture Organization of the United Nations (FAO) and the International Institute for Applied Systems Analysis (IIASA) (156), which are extensively used in crop potential yield modelling. For example, in the Yangtze River basin, from 2010 to 2015, Liu et al estimated the rapeseed production potential on winter fallow fields by using the GAEZ model (157). Pu et al. (2019) provided a full overview of the GAEZ model's calculating techniques and validation (158), and then, Pu et al. (2020) followed the model principles and investigated the influence of climate change on prospective maize and paddy rice yields in Northeast China from 2015 to 2050 (159). The GAEZ model has shown excellent global-scale capability in predicting crop yields by comparing simulated yields with FAO statistics (160).

### 3.2 The improved GAEZ (iGAEZ) model

Here, we conducted a comprehensive comparative analysis and utilized an improved GAEZ model to assess the global potential yield of *Camellia oleifera* cultivation. This approach was combined with a global assessment of *Camellia oleifera* land suitability to determine the potential planting space for *Camellia oleifera*, while also revealing the interaction mechanisms between fresh fruits of *Camellia oleifera* yield and environmental conditions. The iGAEZ model generates the optimal crop parameter of the whole growth cycle to provide the best realistic crop yield combinations while taking regionally differentiated climatic – social - economic status and crop condition into account. The details of each procedure in the improved GAEZ model are as follows.

#### 3.2.1 The photosynthetic potential yield

The photosynthetic potential yield is defined as the maximum dry matter production that can be synthesized by the energy derived from plant photosynthetic efficiency and light resources, assuming that other factors affecting crop growth are in an ideal state or at their optimal ratio (161). The magnitude of photosynthetic potential yield depends on the photosynthetically active radiation (PAR), which represents the maximum solar energy per unit area that plants can convert through photosynthesis. The formula for calculating the photosynthetic potential (Y(Q), ($kg \cdot hm^{-2} \cdot a^{-1}$)) and the correction function f(Q) for the effective solar radiation coefficient are presented as follows:

$$Y(Q) = K \times f(Q) \tag{24}$$

$$f(Q) = \Omega\, \varepsilon\varphi\alpha(1-\rho)(1-\gamma)(1-\omega)E\frac{\sum Q_i}{q(1-\eta)(1-\xi)} \tag{25}$$

where K is the conversion coefficient, equal to 10,000 (162); $\sum Q_i$ represents the total annual surface solar radiation, unit in $MJ \cdot m^{-2}$; $\alpha$ is the absorptivity of leaf area for photosynthetically active radiation, calculated according to equations (26) and (27). $\varepsilon$ represents the proportion of photosynthetically active radiation in the total solar radiation available for photosynthesis, assumed to be 0.49 (161). $\varphi$ and $\rho$ are the average values for most subtropical hilly area crops (163). $\omega$ is the respiratory loss rate in plants, assumed to be 0.3 (164). $E$ is the economic coefficient or the crop harvest index, which indicates the ability of crops to convert organic matter into yield (143). For oil crops, the economic coefficient is generally lower, ranging from 0.3 to 0.4. Here, we take the median value for *Camellia oleifera*, $E$=0.35. $q$ is the energy content per unit of dry matter, taken as 17.20 KJ/g (143). $\eta$ is the moisture content of *Camellia oleifera* seed and determined according to the testing method specified in the *National Standard for Food Safety Determination of Moisture in Food* (GB 5009.3-2016). According to the requirements specified in *Camellia oleifera Seeds* (GB/T 37917-2019), the moisture content should not exceed 13.0% (165). Here, we take the upper limit value, η=13.0%. The remaining coefficients, including $\Omega$, $\gamma$, and $\xi$, are obtained from previous research on *Camellia oleifera* as reported in Table S19 (143,161-167).

Specially, the total surface solar radiation $Q$ is closely related to the extraterrestrial solar radiation and the sunshine hours, which is calculated by the following formula:

$$\mathrm{Q}=\mathrm{Q}_a \times (\mathrm{a} + \mathrm{bS}) \tag{26}$$

$$\mathrm{S}=\frac{n}{\mathrm{N}} \times 100\% \tag{27}$$

where $Q$ is the total surface solar radiation (MJ • $m^{-2}$) and is calculated by using the FAO Ångström-Prescott equation (168). $\mathrm{Q}_a$ is the extraterrestrial solar radiation, while a and b are empirical coefficients that vary by season and can be obtained using measured radiation and meteorological data through the least squares regression model (169). $S$ denotes the sunshine fraction, which is the ratio of actual sunshine duration hours ($n$) and maximum possible sunshine hours ($N$). Here, we used the EARTHDATA provided by NASA to select the M2TMNXRAD (or tavgM_2d_rad_Nx) dataset, which is a time-averaged 2-dimensional monthly mean data collection in Modern-Era Retrospective analysis for Research and Applications version 2 (MERRA-2) with spatial resolution of 0.5° × 0.625° (170). Although the resolution of the M2TMNXRAD dataset is lower than that of the Dataset of high-resolution (3 hours, 10 km) global surface solar radiation, the latter dataset is only updated to 2018 (171). Each pixel was resampled into four new cells to obtain a 5 arcmin resolution and multiplied by the number of days in the corresponding month to convert from daily to monthly values.

Considering the changes in reflectance and transmittance caused by variations in leaf area index ($L$), the crop canopy light absorption efficiency $\alpha$ throughout the entire growth period is represented by a linear function that increases with leaf area. The equation is as follows:

$$\alpha=0.83f(L) = 0.83 * \mathrm{L_i}/\mathrm{L_0} \tag{28}$$

where $f(L)$ represents the correction factor for the dynamic changes in crop leaf area. During the entire crop growth period, the crop canopy light absorption efficiency is a function of leaf area growth, given by $f(L) = \mathrm{L_i}/\mathrm{L_0}$. Here, $\mathrm{L_0}$ denotes the maximum leaf area index, and $\mathrm{L_i}$ represents the leaf area index at a specific time interval. It is difficult to achieve the optimal structure when $\mathrm{L_i}/\mathrm{L_0}$ equals 1. Due to limited reports on the climatic potential yield of *Camellia oleifera* trees, we adopted the average reference value from most crops such as rice, wheat, and maize, which determines $f(L)$ as 0.58 (172,173).

#### 3.2.2 Light-temperature potential yield

The photosynthetic potential yield represents the optimal growing environment for crops and indicates the maximum photosynthetic production under ideal conditions. However, achieving this optimal state in field cultivation is challenging. When soil nutrients, water availability, and agricultural practices are optimized, the yield per unit area is jointly determined by light and temperature. Therefore, considering that different crops have varying temperature requirements during their growth stages, we introduce a temperature correction function to adjust the photosynthetic potential yield Y(Q) and obtain the light-temperature potential yield. The calculation formula for the light-temperature potential yield (Y(T), kg • km$^{-2}$ • a$^{-1}$) is as follows:

$$\mathrm{Y(T)} = \mathrm{Y(Q)} \times \mathrm{f(T)} \tag{29}$$

$$\mathrm{f(T)} = \frac{(T-T_1)(T_2-T)^b}{(T_0-T_1)(T_2-T_0)^b} \tag{30}$$

$$\mathrm{b} = (T_2 - T_0)/(T_0 - T_1) \tag{31}$$

where f(T) represents the temperature correction function; $T$ denotes the average temperature during the growing period of *Camellia oleifera*. $T_0$, $T_1$, and $T_2$ are three critical temperatures within the whole growth period, representing the optimal temperature, minimum temperature, and maximum temperature, respectively. At the optimal temperature, crop growth and development are rapid. At the highest and lowest temperatures, crop growth ceases but the plants can still survive. However, further increases or decreases in temperature can cause varying degrees of harm to the crops, ultimately leading to death. Based on the characteristics of *Camellia oleifera*, the following critical temperature points have been determined: ① The optimal temperature ($T_0$) range for *Camellia oleifera* growth is between 19-21℃, within which the tree shoots grow rapidly, extending by an average of 1-2 centimeters per day. Here, we take the average value for $T_0$, which is determined as 20℃ (97). ② Although *Camellia oleifera* branches have strong resistance to low temperatures, especially the shrubby small-leaved varieties can tolerate temperatures as low as -10℃. However, from December to February, the average temperature generally drops below 10℃, causing the buds of *Camellia oleifera* to stop sprouting and enter a dormant state for winter. Occasionally, severe frosts occur, causing damage to *Camellia oleifera* seedlings, young trees, or less cold-resistant varieties. Thus, $T_1$ is determined as 10℃. ③ *Camellia oleifera* prefers warm temperatures. According to the comprehensive review conducted by Jiang et al., 83% of the literature indicates that the upper limit of extreme highest temperature suitable for *Camellia oleifera* growth is 40℃ (174). However, the upper limit of the average hottest month temperature suitable for growth is 31℃, a value supported by 100% of the peer-reviewed literature. Therefore, $T_2$ is determined as 31℃.

#### 3.2.3 Climatic potential yield

The climatic potential yield of crops depends on three factors: light, temperature, and water. The requirement for water is a fundamental necessity for plant growth, as it serves as a raw material for photosynthesis and participates in essential physiological processes related to the synthesis and transformation of plant organic matter. The lack of water severely limits the productivity of crops. Therefore, we introduce a water correction function based on the light-temperature potential yield Y(T) to obtain the climate potential yield (Y(C), $kg \cdot km^{-2} \cdot a^{-1}$). As *Camellia oleifera* is a drought-tolerant crop, relying on natural precipitation as a water source, the irrigation coefficient is determined to be zero. In this study, the water-use efficiency factor for *Camellia oleifera* cultivation is primarily calculated based on the water demand and actual evapotranspiration. The calculation formula is as follows:

$$\mathrm{Y(C)} = \mathrm{Y(T)} \times \mathrm{f(W)} \tag{32}$$

$$\mathrm{f(W)} = \begin{cases} 1 - K_y \times (1 - P/ET_c) & (0 \le P < ET_c) \\ 1 - \frac{P - ET_c}{3ET_c} & (ET_c \le P < 4ET_c) \\ 0 & (P \ge 4ET_c) \end{cases} \tag{33}$$

where f(W) represents the water use efficiency correction function, where $0 < \mathrm{f(W)} < 1$. $K_y$ is the crop yield response coefficient, considering that among various meteorological factors, precipitation has the greatest impact on the oil content of *Camellia oleifera* seeds (175). Adequate precipitation and water supply significantly improve the quality of seed oil, thus determining $K_y = 1$ (176). $P$ represents the total precipitation in units of mm/yr. The estimation of crop evapotranspiration follows the recommended crop water requirement method by the Food and Agriculture

Organization (FAO) of the United Nations. The crop's actual evapotranspiration ($ET_c$) during the growth period is the product of the reference crop potential evapotranspiration ($ET_0$) and the crop coefficient ($K_c$). The calculation formula is as follows:

$$ET_c = \sum_{i=1}^{\mathrm{gd}_j}(ET_{0,i} \times K_{c,i}) \tag{34}$$

where $ET_c$ represents crop evapotranspiration (mm/yr); $\mathrm{gd}_j$ is the growing length of *Camellia oleifera* in the different stages； $K_{c,i}$ is the daily average crop coefficient. The crop coefficient for *Camellia oleifera* ( $K_c$ ) is adopted from the standard crop coefficients of 84 crops recommended by the Food and Agriculture Organization (FAO-56) (169). And then, we combined the sentinel monitoring data from Wang et al to identify the monthly change pattern of the crop coefficient in different growth periods (see Section 1.2.2 and Figure S4) (39). $ET_{0,i}$ is the reference crop potential evapotranspiration (mm/day). $ET_0$ is related to various parameters such as the daily minimum and maximum temperatures, average relative humidity, wind speed, sunshine hours, and net radiation during the crop growth period. The CROPWAT 8.0 model is used in combination with meteorological data and the FAO Penman-Monteith equation to estimate $ET_0$ (170,177). The calculation process is described as follows:

$$ET_0 = \frac{0.408\Delta \times (R_n - G) + 900\gamma/(T_{mean} + 273) \times U_2 \times (e_s - e_a)}{\Delta + \gamma(1 + 0.34U_2)} \tag{35}$$

where $\Delta$ is the saturated water pressure curve slope (kPa/℃); $R_n$ is the ground surface radiation (MJ/(m$^2$*d)); $G$ is the soil heat flux (MJ/(m$^2$*d)), $G \approx 0$; $\gamma$ is the wet and dry constant (KPa/°C); $T_{mean}$ is the Daily average temperature at 2 meters (°C); $U_2$ is the wind speed at 2 meters (m/s); $e_s$ is the saturated water pressure (Kpa); $e_a$ is the Actual water pressure (Kpa); The detailed calculation of $\Delta$, $R_n$, $\gamma$, $T_{mean}$ and $U_2$ are described in the manual "FAO Irrigation and Drainage Paper, No. 56, Crop Evapotranspiration" (170), and can be estimated by using the software "$ET_0$ Calculator" (178). The specific calculation process is as follows:

**Parameter (1) $\Delta$:**

$$\Delta = \frac{4098 \times \left[0.6108 \times \exp\left(\frac{17.27T}{T+237.3}\right)\right]}{(T+237.3)^2} \tag{36}$$

where $T$ is the daily average temperature (°C).

**Parameter (2) $R_n$:**

$$R_n = R_{ns} - R_{nl} \tag{37}$$

where $R_n$ is the net radiation, which is the difference between the incoming short-wave radiation $R_{ns}$ and net outcoming long-wave radiation $R_{nl}$.

①The calculation process of $R_{ns}$ is described as follows:

$$R_{ns} = (1-\alpha)R_s \tag{38}$$

$$R_s = (a_s + b_s \frac{n}{N})R_a \tag{39}$$

$$N = \frac{24}{\pi} W_s \tag{40}$$

$$W_s = arccos[-tan\,(\varphi)tan\,(\delta)] \tag{41}$$

$$\delta = 0.408 sin\,(\frac{2\pi}{365}J - 1.39) \tag{42}$$

$$R_a = \frac{24(60)}{\pi} G_{sc} \times d_r[W_s\ sin(\varphi)\,sin(\delta) + cos(\varphi)\,cos(\delta)\,sin(W_s)] \tag{43}$$

$$d_r = 1 + 0.033\,cos\left(\frac{2\pi}{365}J\right) \tag{44}$$

where α =0.23 by taking the albedo of the reference crop in grassland; $R_s$ is the sun radiation; $a_s$ =0.25, $b_s$ =0.50; $n$ is the actual sunshine hours (h). $N$ is the maximum possible sunshine hours (h). $W_s$ is the angle at sunrise; φ is latitude (rad); δ is solar declination; $J$ is day order, $J = 1 - 365/366$. $R_a$ is the extraterrestrial radiation; $G_{sc}$ =0.0820, meaning the solar constant (MJ/($m^2$*min)); $d_r$ is the sun-earth average distance.

②The calculation process of $R_{nl}$ is described as follows:

$$R_{nl} = \delta\left[\frac{T_{max,k}^4 + T_{min,k}^4}{4}\right](0.34 - 0.14\sqrt{e_a})(1.35\frac{R_s}{R_{s0}} - 0.35) \tag{45}$$

$$\delta = 4.903 \times 10^{-9} \tag{46}$$

$$T_{max,k} = T_{min} + 272.15 \tag{47}$$

$$T_{min,k} = T_{min} - 272.15 \tag{48}$$

$$e_a = RH \times e_S \tag{49}$$

$$e_S = \frac{e(T_{max}) + e(T_{min})}{2} \tag{50}$$

$$e(T_{max}) = 0.6108 \times \exp\,(\frac{17.27T_{max}}{T_{max} + 237.3}) \tag{51}$$

$$R_{s0} = (a_s + b_s)R_a \tag{52}$$

where $\delta$ is Stephen Boltzmann constant ($M \cdot K^{-3} \cdot m^{-2} \cdot d^{-1}$); $e_a$ is the actual water pressure; $RH$ is the relative humidity; $e_S$ is the saturated water pressure.

**Parameter (3) $\gamma$:**

$$\gamma = 0.665 \times 10^{-3} P \quad (53)$$

**Parameter (4) $T_{mean}$:**

$$T_{mean} = \frac{T_{max}+T_{min}}{2} \quad (54)$$

**Parameter (5) $U_2$:**

$$U_2 = U_s \frac{4.87}{\ln(67.8s-5.42)} \quad (55)$$

where $U_2$ is the wind speed at 2 meters; $U_s$ is wind speed at 10 meters, $s$ =10. This is mainly because the wind speed data observed at meteorological stations worldwide is recorded at a height of 10 meters above the ground, while in agricultural climate analysis, the standard crop canopy height is generally considered to be at a height of 2 meters above the ground. Therefore, it is necessary to use a wind speed attenuation coefficient to convert the observed wind speed data at a height of 10 meters above the ground to wind speed data at a height of 2 meters (179,180).

#### 3.2.4 Natural comprehensive potential yield

The natural comprehensive potential yield (Y(L)) is estimated based on the climatic potential yield $Y(C)$ considering the soil correction function $f(S)$, which incorporates soil properties and composition to assess the potential yield due to land constraints. We selected four key indicators, including soil cation exchange capacity (cmol/kg), total organic carbon content (g/kg), slope, and elevation (m), as factors influencing the soil correction function. The formula for calculating the natural comprehensive potential yield (Y(L), kg • $hm^{-2}$ • $a^{-1}$) is as follows:

$$\mathrm{Y(L)} = \mathrm{Y(C)} \times \mathrm{f(S)} \quad (56)$$

$$\mathrm{f(S)} = \sum_{i=1}^{n} W_i A_i \quad (57)$$

where $0 < f(S) < 1$; $n$ = 4; $W_i$ is the weighting factor for the indicator $i$. We assume that each indicator is equally important, $W_i$=0.25. $A_i$ is the scoring value for the indicator $i$. Referring to Wang et al, it was determined to divide into 5 levels between 0.7 and 0.9 as the scoring of soil effective coefficient, which is shown in Table S20 (143).

### 3.3 Conversion from *Camellia oleifera* fresh fruits to seed for oil processing

As the projection of production capacity can only predict the weight of fresh fruits in *Camellia oleifera*, it is necessary to convert the value of fresh fruit to dry seed weight for vegetable oil processing. Specially, we select the following four *Camellia oleifera* clones' information to get the mean value of dry seed rate in fresh fruit as $R_c$= 35.65% (Table S21) (53). These four *Camellia oleifera* clones are "Ganwu 1", "Changlin 4", "Changlin 40", and "Ganyong 5", among which the first three clones are listed as mostly recommended varieties of *Camellia oleifera* (181). The fruit characteristics and yield of *Camellia oleifera* are not only controlled by genetic factors, but also significantly influenced by environmental conditions and management practices, and the fruits exhibit biennial bearing. Here, the average values of fruit characteristics of four *Camellia oleifera* clones show the result of over two consecutive years of monitoring to reflect the actual level. We use this relatively conservative estimation of dry seed rate in fresh fruit also for our subsequent analysis of environmental impacts under multi-yield potential scenarios simulation.

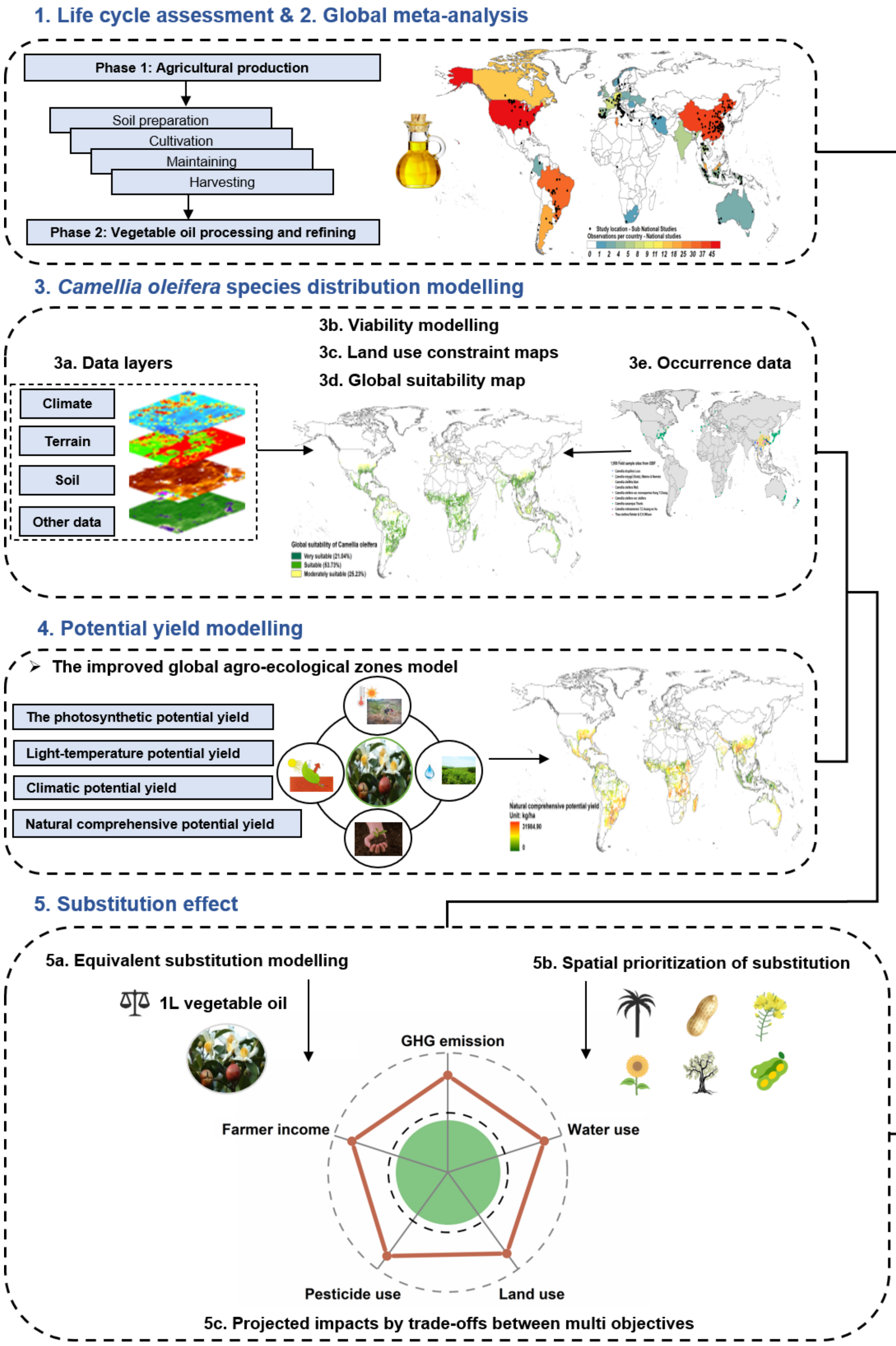


**Fig. S1.** Methodological roadmap.

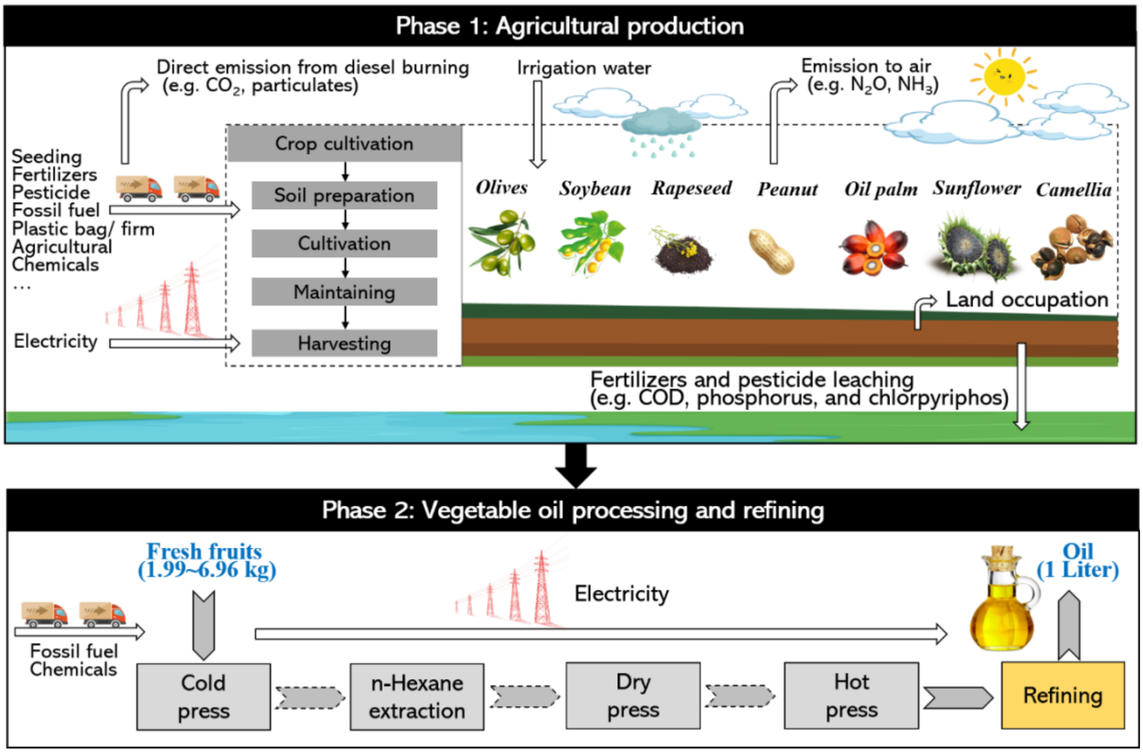


**Fig. S2.** System boundary of the life cycle assessment for seven vegetable oil crops.

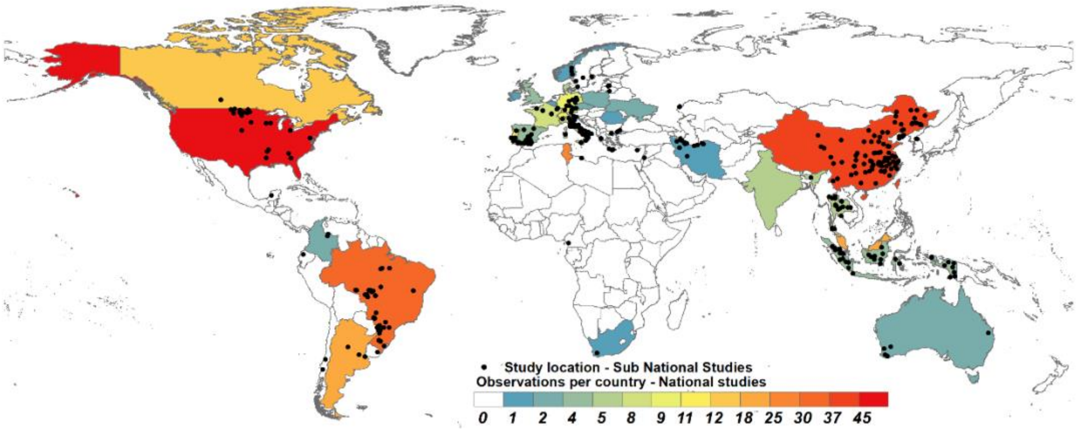


**Fig. S3.** Study locations of vegetable oil crops in global meta-analysis. Country coloring indicates the number of national-level studies per country (n observations = 272); black circles indicate the locations of subnational studies (n observations = 565).

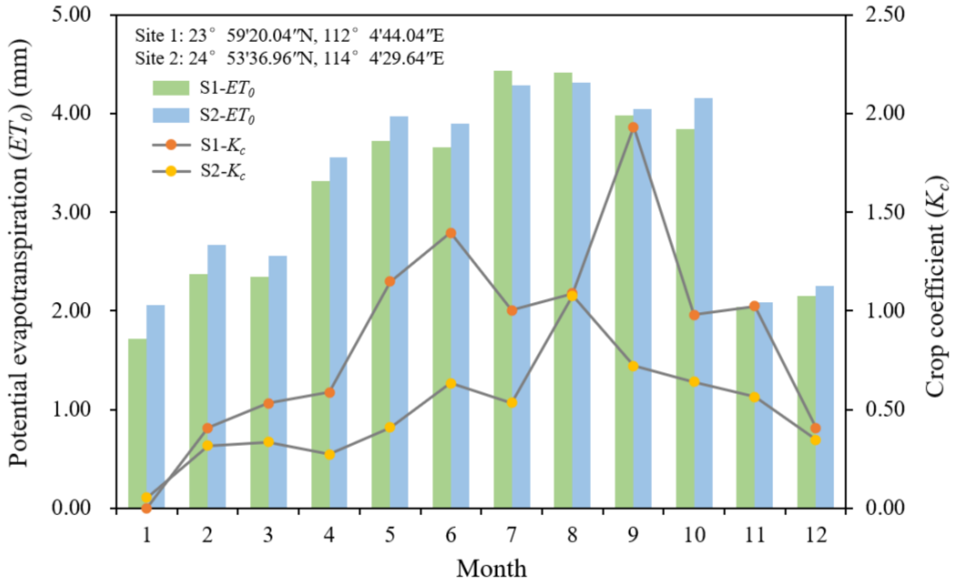


**Fig. S4.** Monthly variation of the reference crop potential evapotranspiration ($\boldsymbol{ET_0}$) and the crop coefficient ($\boldsymbol{K_c}$) for Camellia oleifera. Site 1: Huaji County, Zhaoqing City, Guangdong Province, China (23°59'20.04″N, 112°4'44.04″E). Site 2: Shixing County, Shaoguan City, Guangdong Province, China (24°53'36.96″N, 114°4'29.64″E).

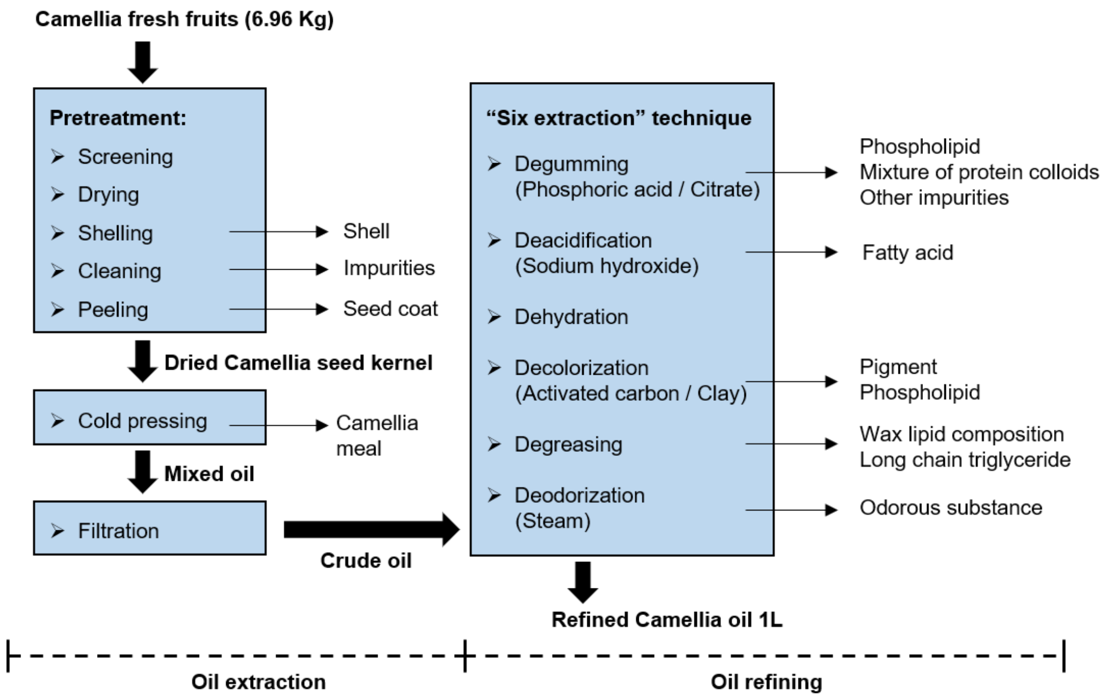


**Fig. S5.** Processing technique of *Camellia oleifera* oil production

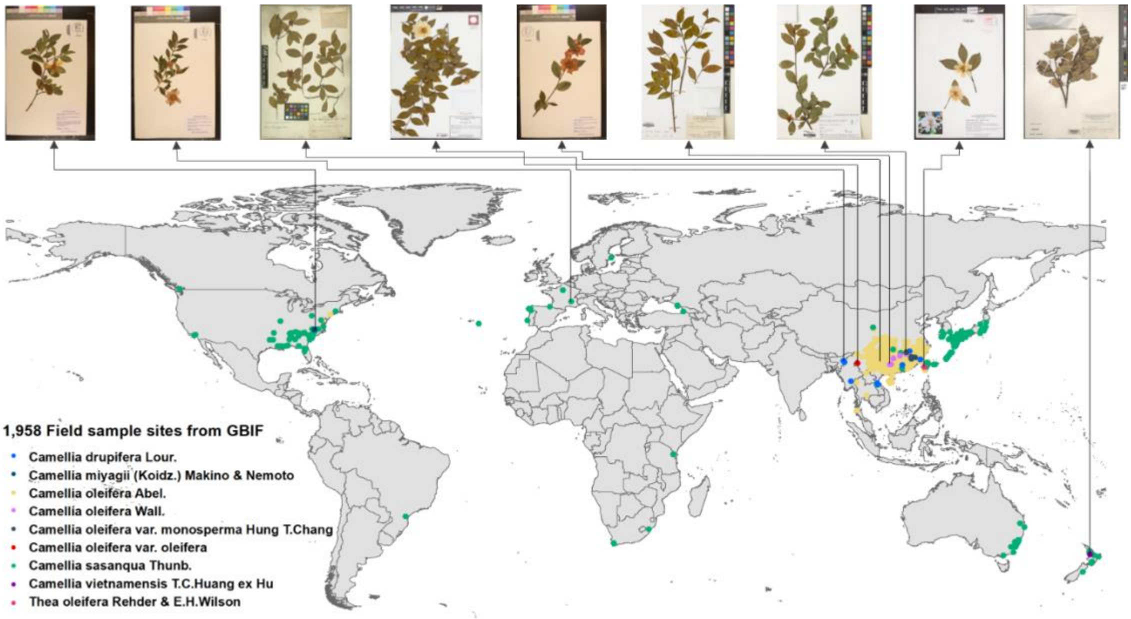


**Fig. S6.** Spatial distribution of all 1,958 field sample sites of *Camellia oleifera*.

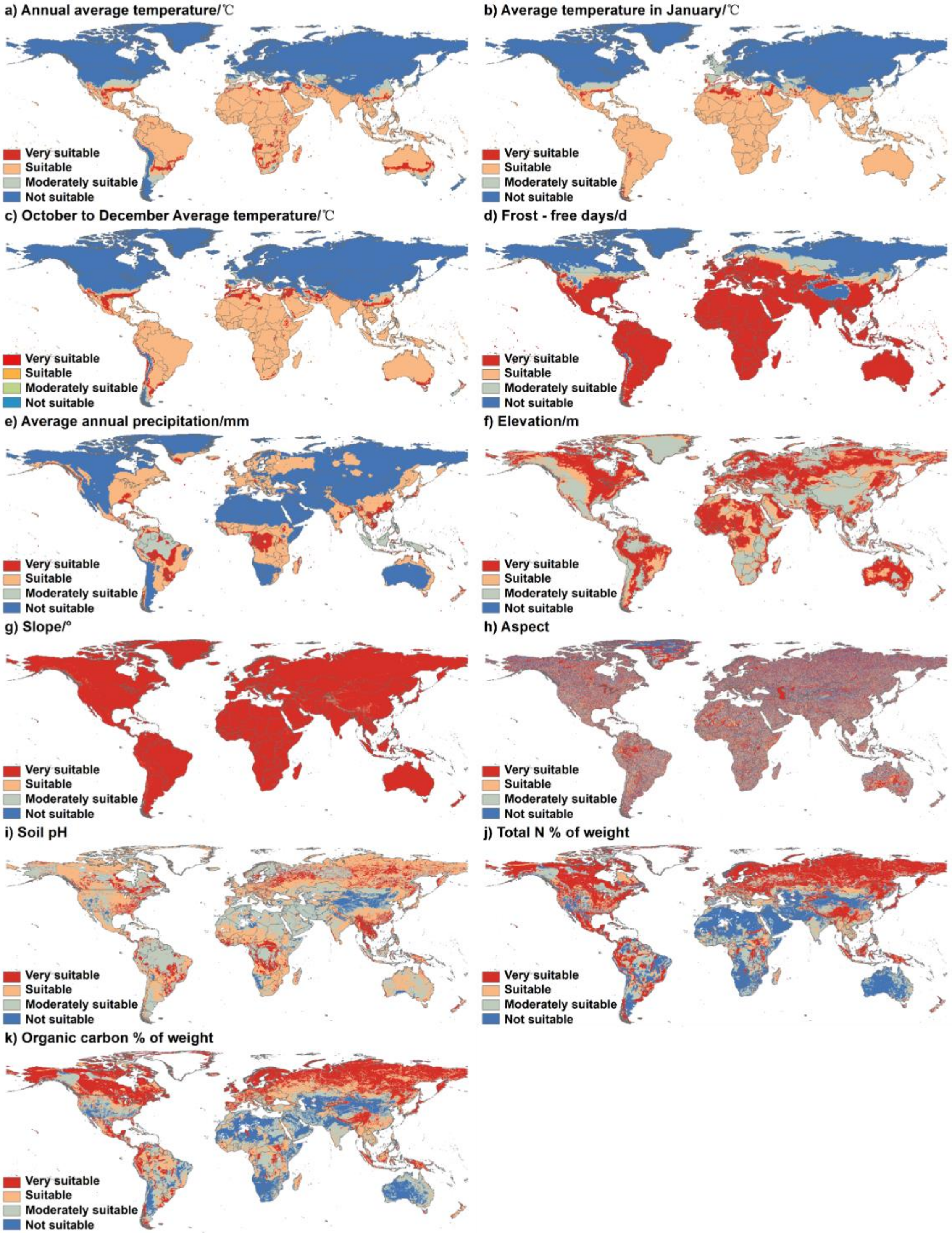


**Fig. S7.** Spatial distribution of each indicator in the suitability assessment of *Camellia oleifera* cultivation. Data sources a-e): Version 4 of CRU TS in https://crudata.uea.ac.uk/cru/data/hrg/; f-h): Harmonized World Soil Database (version 1.2) (FAO/IIASA, Rome/Laxenburg, 2012); and i-k): National Tibetan Plateau / Third Pole Environment Data Center in https://data.tpdc.ac.cn/.

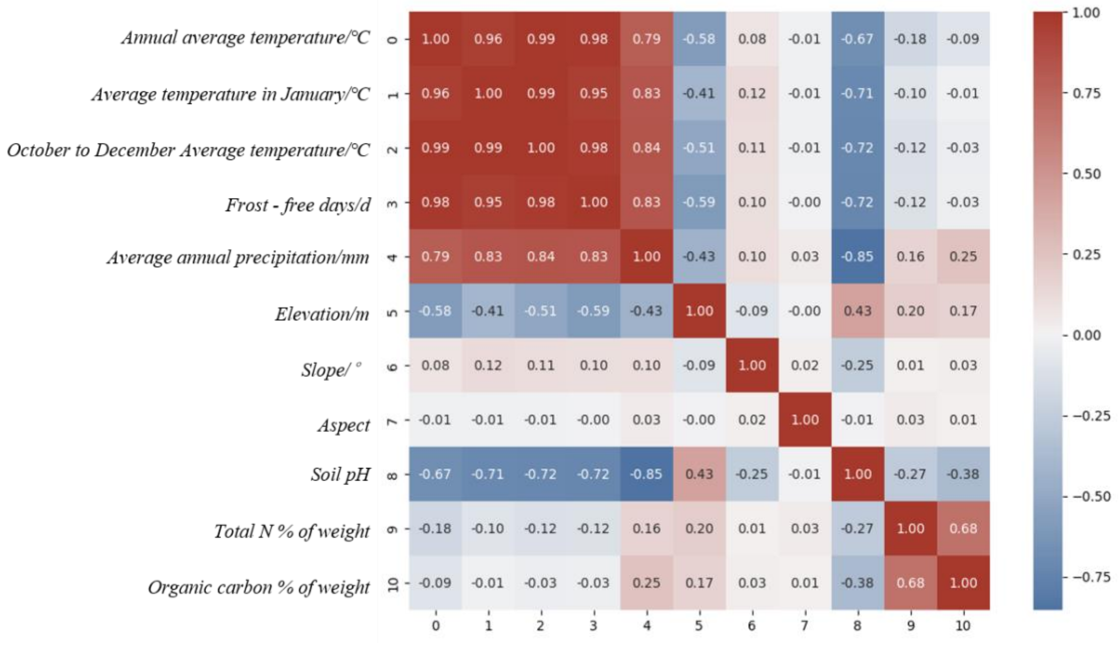


**Fig. S8.** The correlation matrix of each indicator in the *Camellia* suitability assessment.

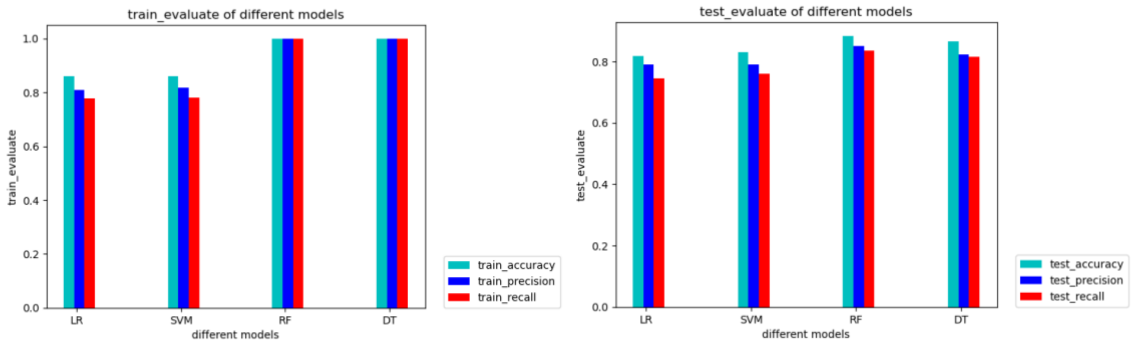


**Fig. S9.** Train and test evaluation of four models for weight identification

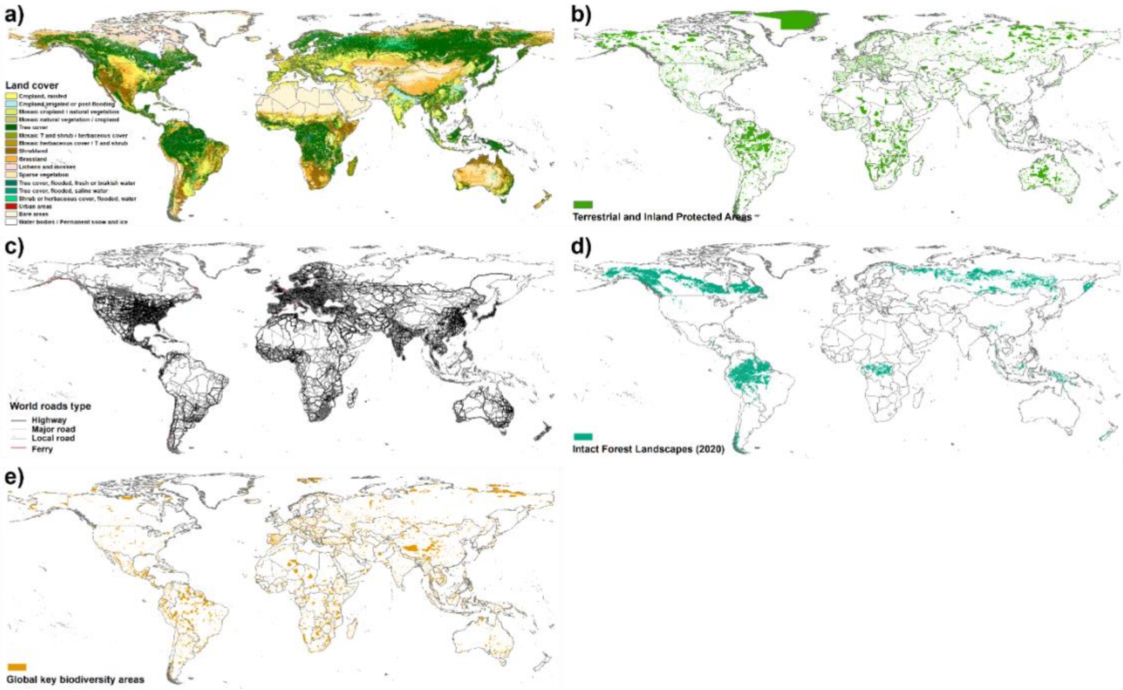


**Fig. S10.** Incorporated land use constraints: a) land cover; b) terrestrial and inland protected areas; c) world roads network; d) intact forest landscapes; and e) global key biodiversity areas

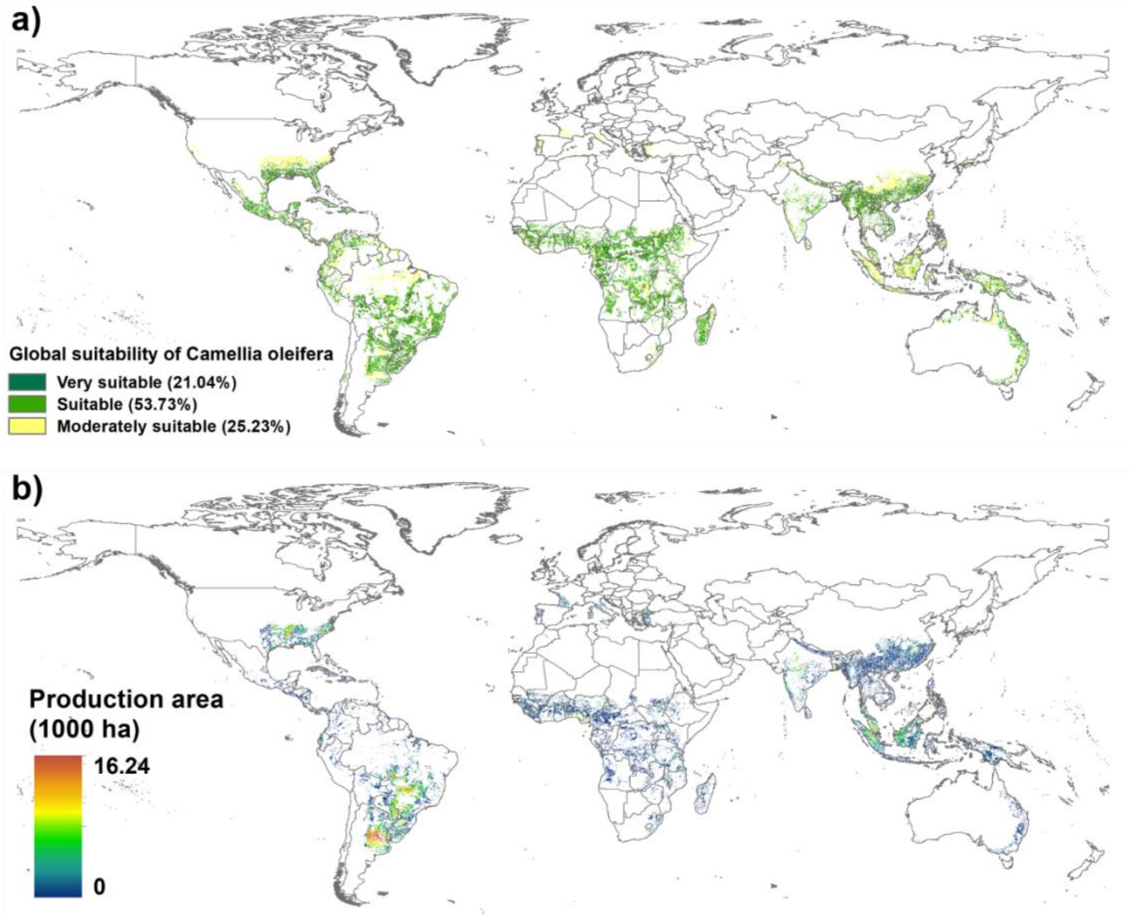


**Fig. S11.** Spatial suitability assessment of *Camellia oleifera* cultivation globally (a) and potential production areas if limited within current vegetable oil plantation areas (b)

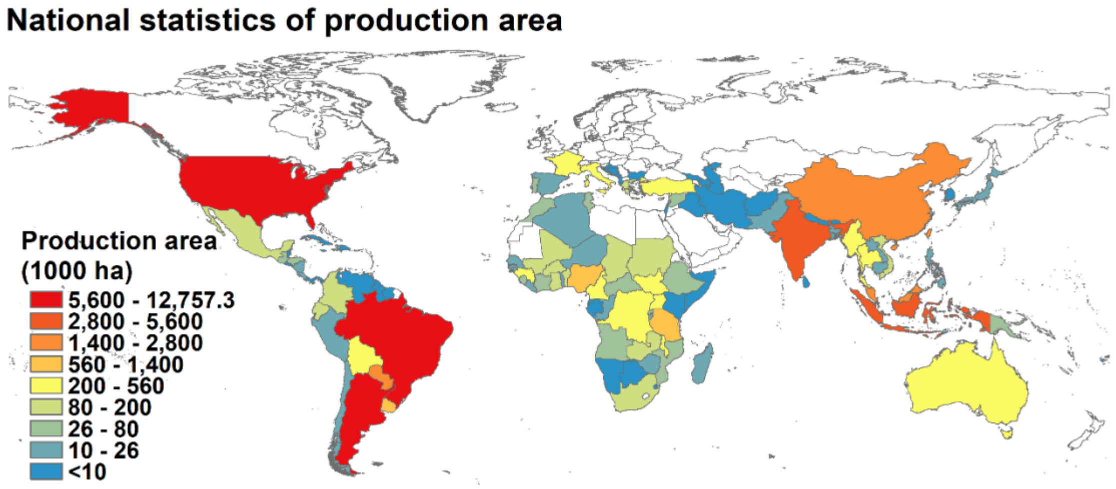


**Fig. S12.** Suitability areas of *Camellia oleifera* at country level if limited within current vegetable oil production areas

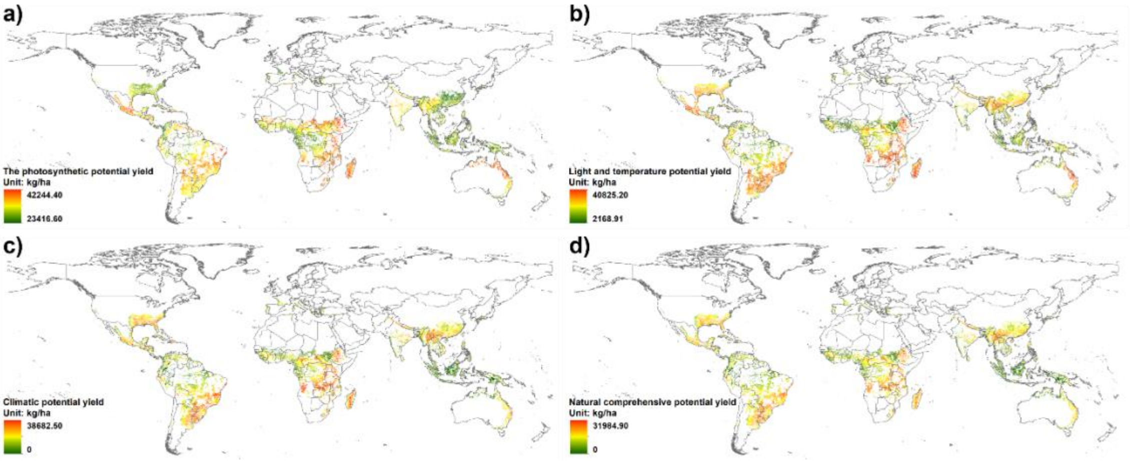


**Fig. S13.** Global spatial distribution of *Camellia oleifera* production capacity: a) photosynthetic potential yield; b) light and temperature potential yield; c) climatic potential yield; and d) natural comprehensive potential yield

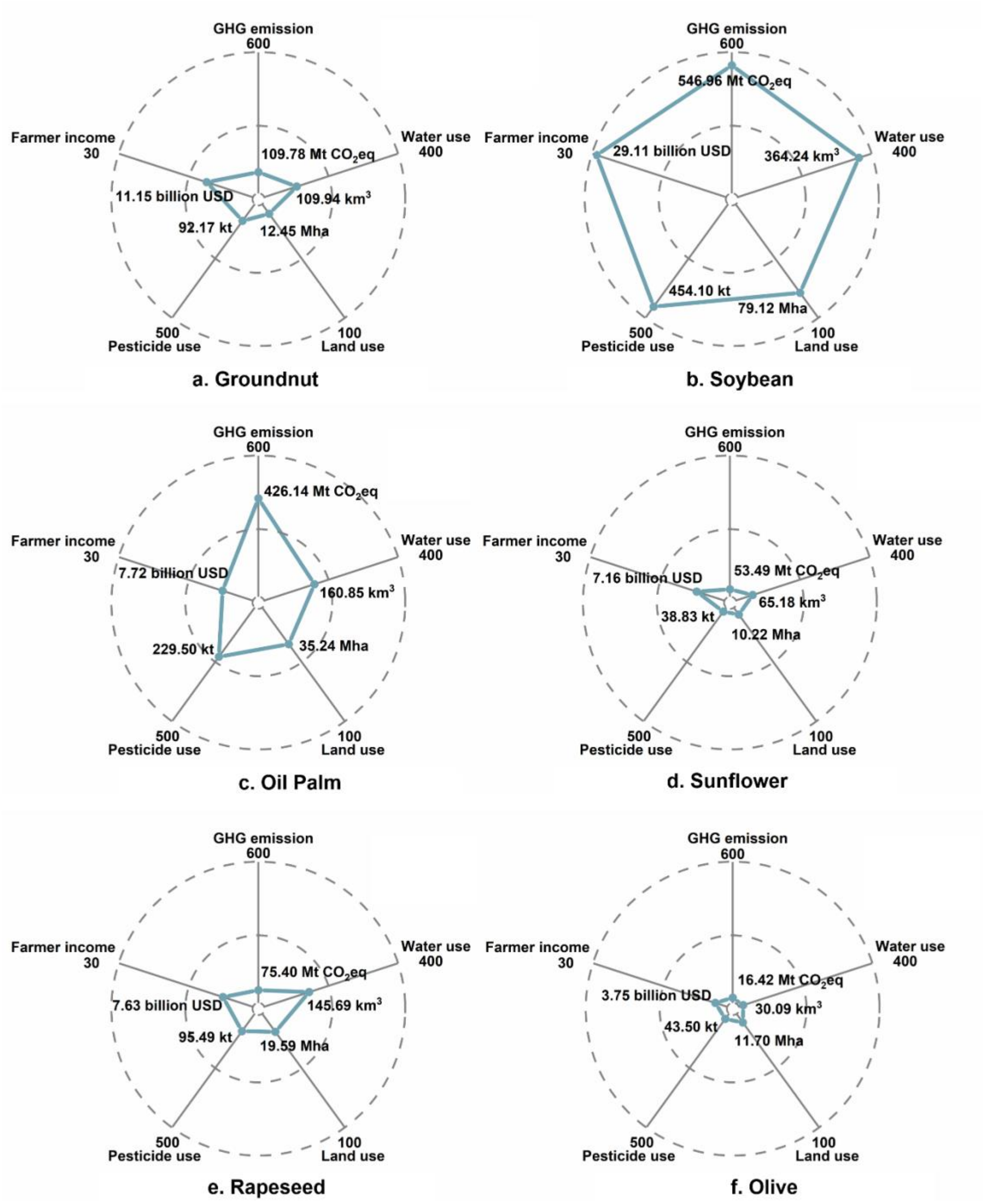


**Fig. S14.** Global environmental and socioeconomic effect of the six main vegetable oil crops per year: a) groundnut, b) soybean, c) oil palm, d) sunflower, e) rapeseed, and f) olive.

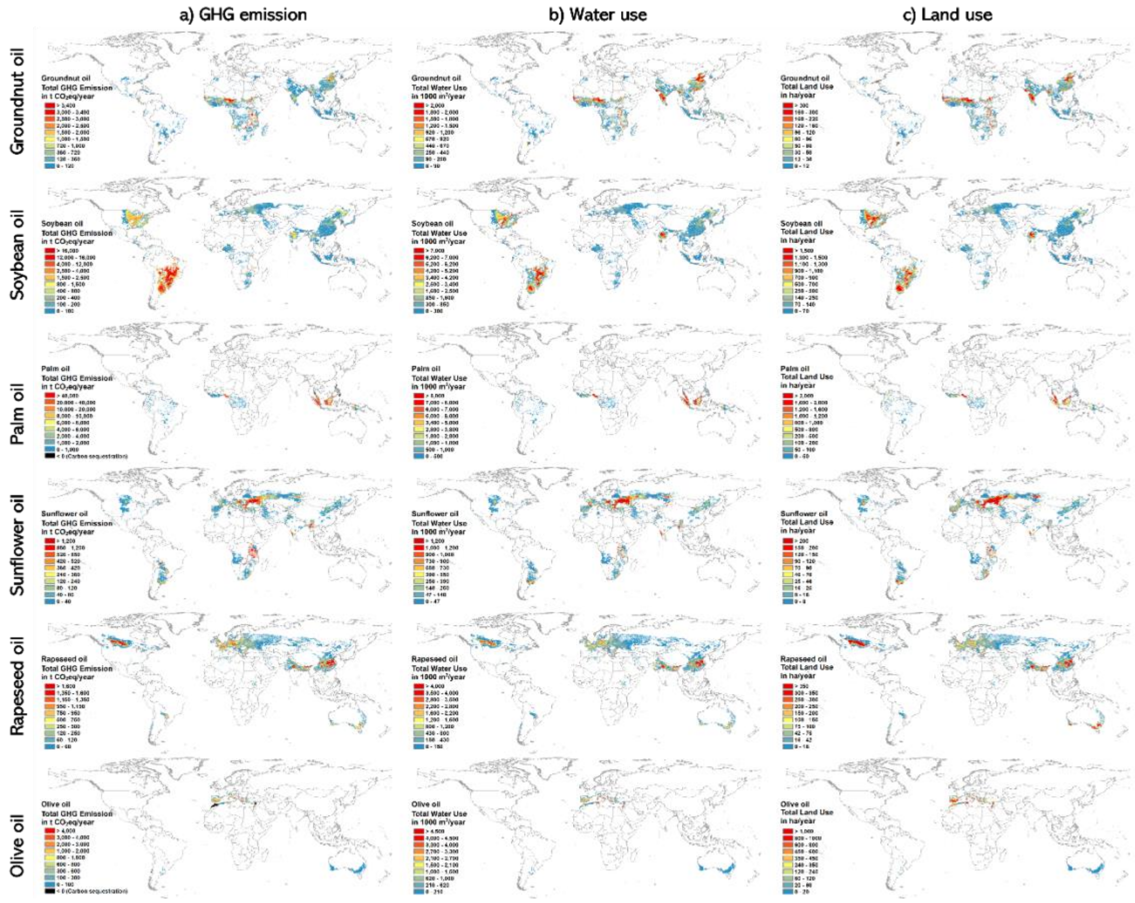


**Fig. S15.** Environmental impacts of six major vegetable oil crops: a) GHG emissions in t $CO_2$eq/year, b) water use in $km^3$/year, and c) land use in ha/year.

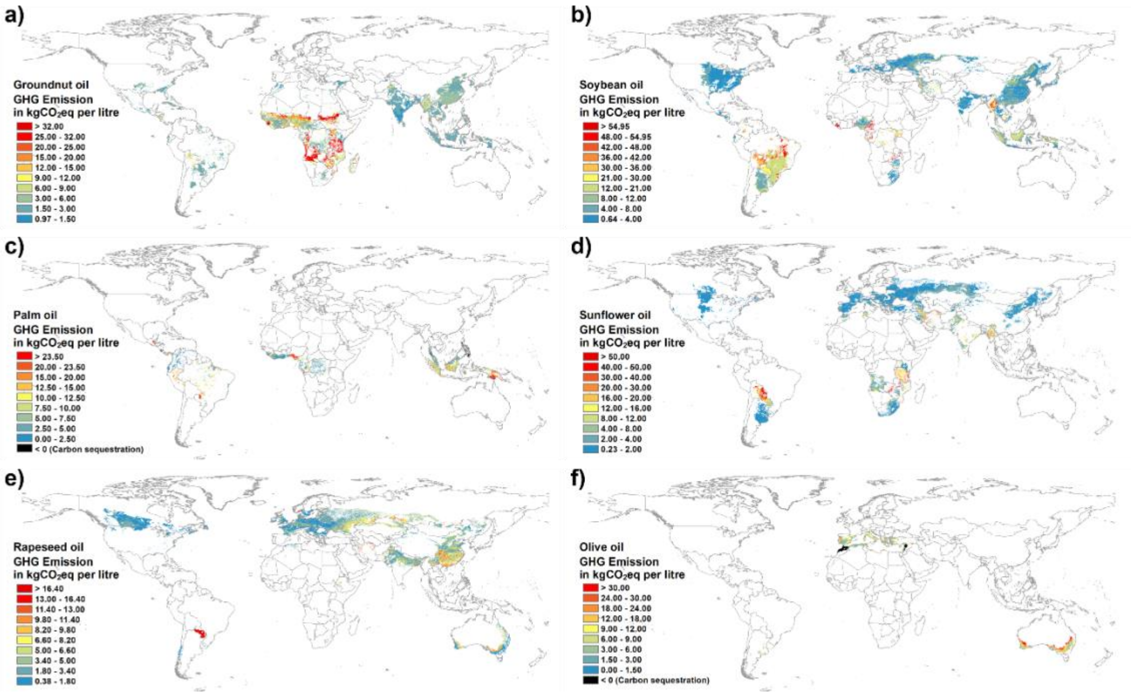


**Fig. S16.** GHG emission per liter vegetable oil across the current six types of crops: a) groundnut oil, b) soybean oil, c) palm oil, d) sunflower oil, e) rapeseed oil, and f) olive oil. The rank of global mean GHG emission intensity is followed by palm oil (9.7 kg $CO_2$ eq/L) > olive oil (8.5 kg $CO_2$ eq/L) > groundnut oil (8.4 kg $CO_2$ eq/L) > soybean oil (8.4 kg $CO_2$ eq/L) > rapeseed oil (4.0 kg $CO_2$ eq/L) > sunflower oil (3.5 kg $CO_2$ eq/L).

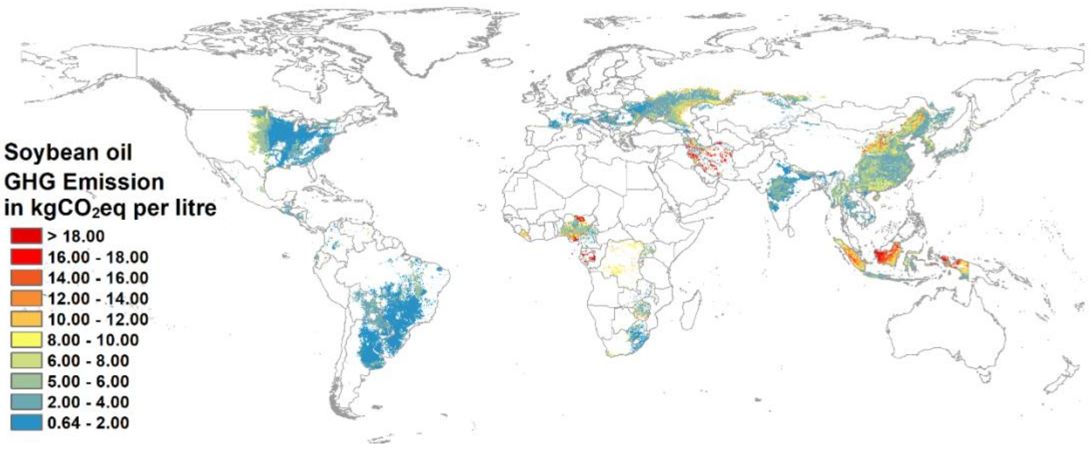


**Fig. S17.** GHG emission per liter soybean oil if excluding the stage of land use change (the above and below-ground C stock change, forest burning, and organic soil burning) (Global mean GHG emission intensity = 4.4 kg $CO_2$ eq/L)

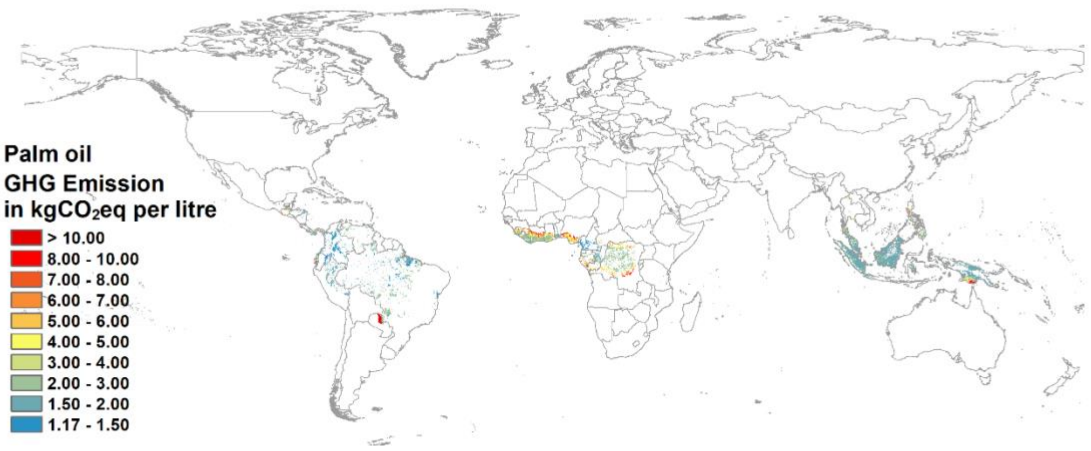


**Fig. S18.** GHG emission per liter palm oil if excluding the stage of land use change (the above and below-ground C stock change, forest burning, and organic soil burning) (Global mean GHG emission intensity = 2.7 kg $CO_2$ eq/L).

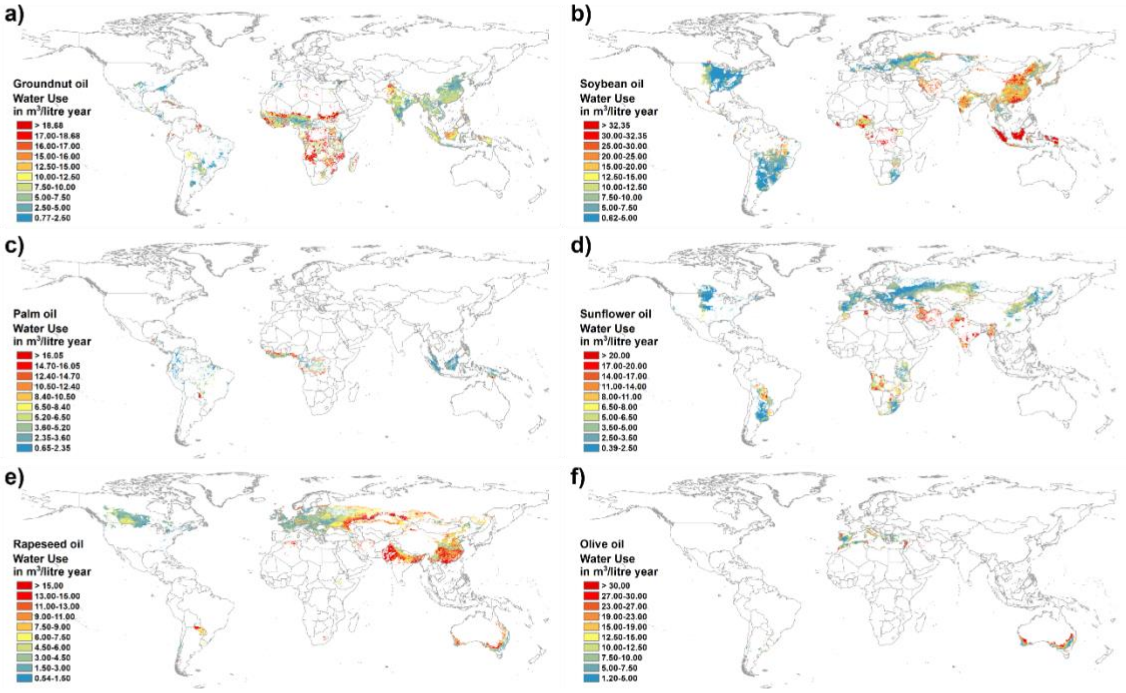


**Fig. S19.** Water use per liter vegetable oil across the current six types of crops: a) groundnut oil, b) soybean oil, c) palm oil, d) sunflower oil, e) rapeseed oil, and f) olive oil. The rank of global mean water use intensity is followed by olive oil (14.5 $m^3/L$) > soybean oil (13.9 $m^3/L$) > groundnut oil (9.7 $m^3/L$) > rapeseed oil (6.8 $m^3/L$) > sunflower oil (5.7 $m^3/L$) > palm oil (5.3 $m^3/L$).

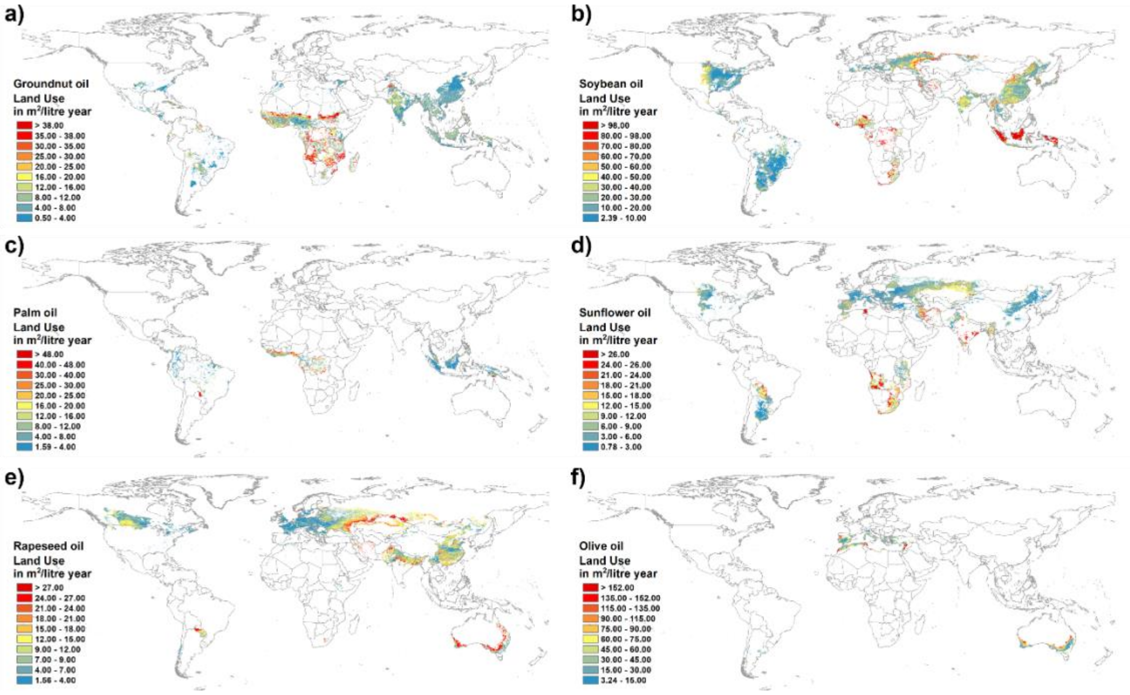


**Fig. S20.** Land use per liter vegetable oil across the current six types of crops: a) groundnut oil, b) soybean oil, c) palm oil, d) sunflower oil, e) rapeseed oil, and f) olive oil. The rank of global mean land use intensity is followed by sunflower oil (8.4 $m^2$/L) > rapeseed oil (10.4 $m^2$/L) > palm oil (11.0 $m^2$/L) > groundnut oil (11.9 $m^2$/L) > soybean oil (31.9 $m^2$/L) > olive oil (51.6 $m^2$/L).

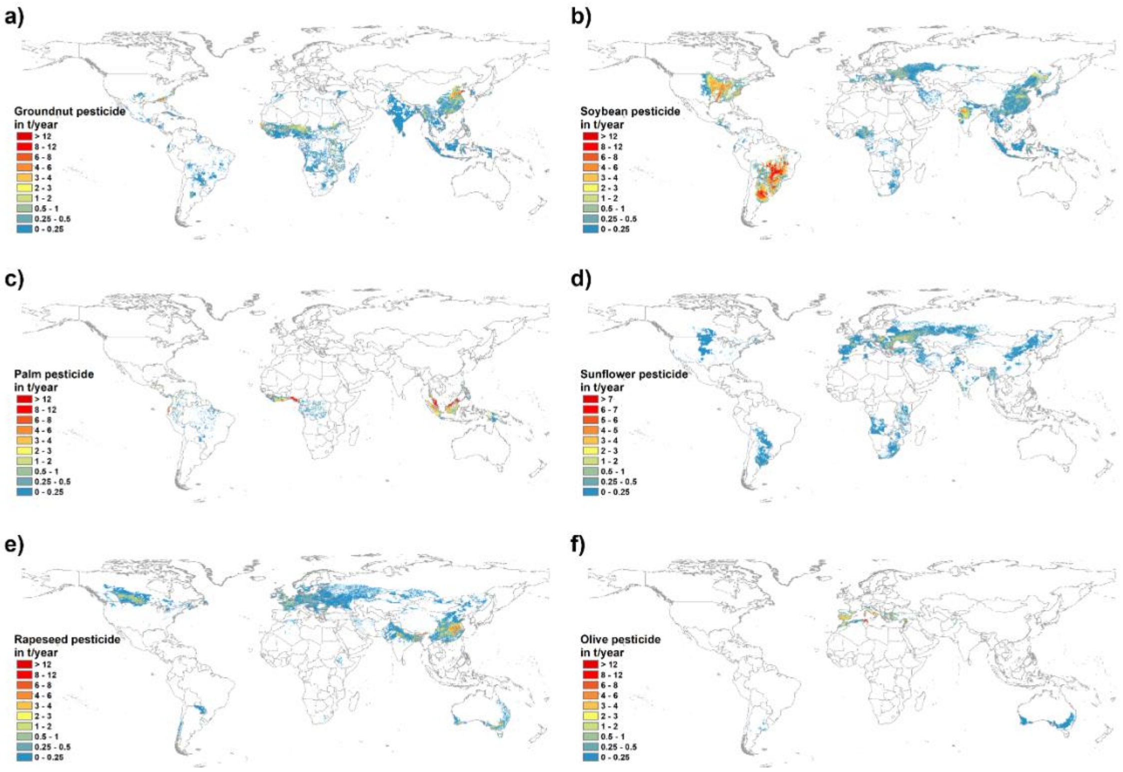


**Fig. S21.** Spatial variation of pesticide use (t/year) in major six vegetable oil crops at the global level: a) groundnut, b) soybean, c) palm, d) sunflower, e) rapeseed, and f) olive

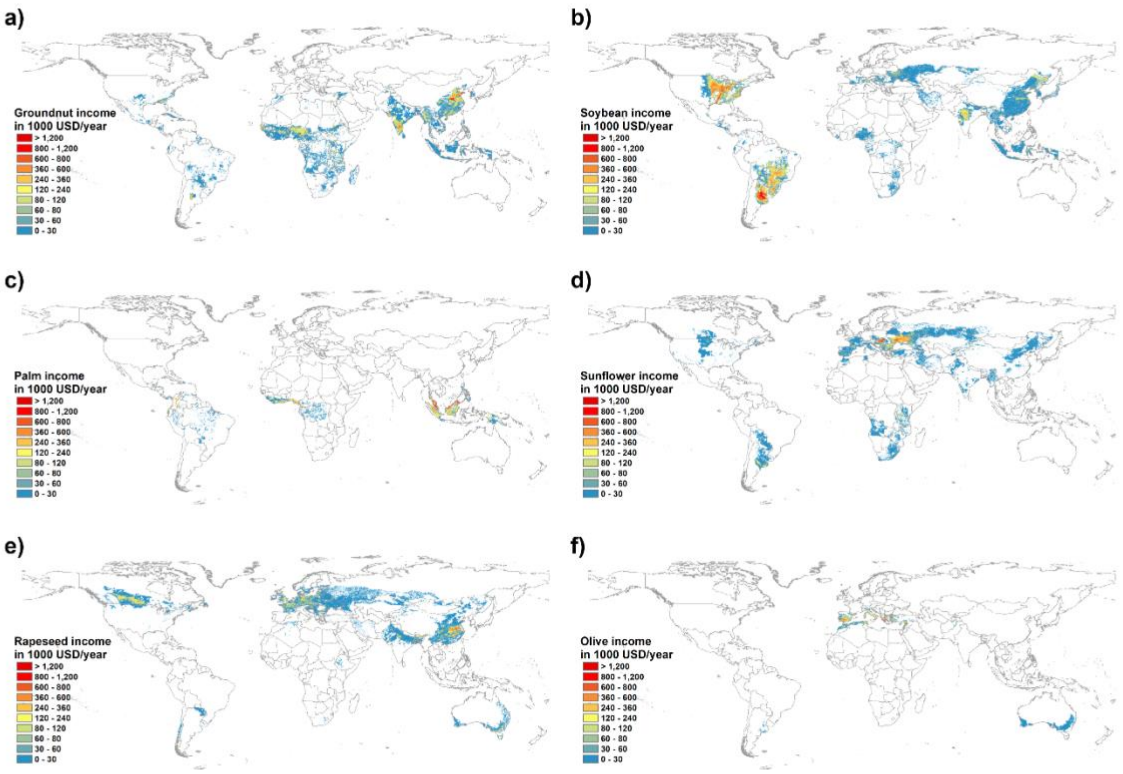


**Fig. S22.** Spatial variation of farmer income (USD/year) in major six vegetable oil crops at the global level: a) groundnut, b) soybean, c) palm, d) sunflower, e) rapeseed, and f) olive

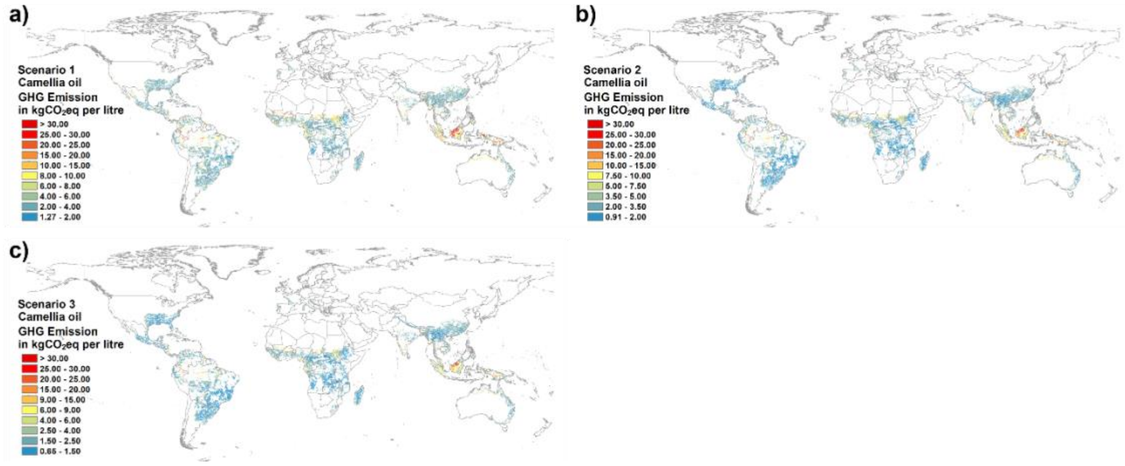


**Fig. S23.** GHG emission per liter *Camellia oleifera* oil under three yield projection scenarios: a) conservative yield scenario (S1; 50% potential yield); b) strategic yield improvement scenario (S2; 75% potential yield); and c) technology-policy-enabling yield scenario (S3; 100% potential yield)

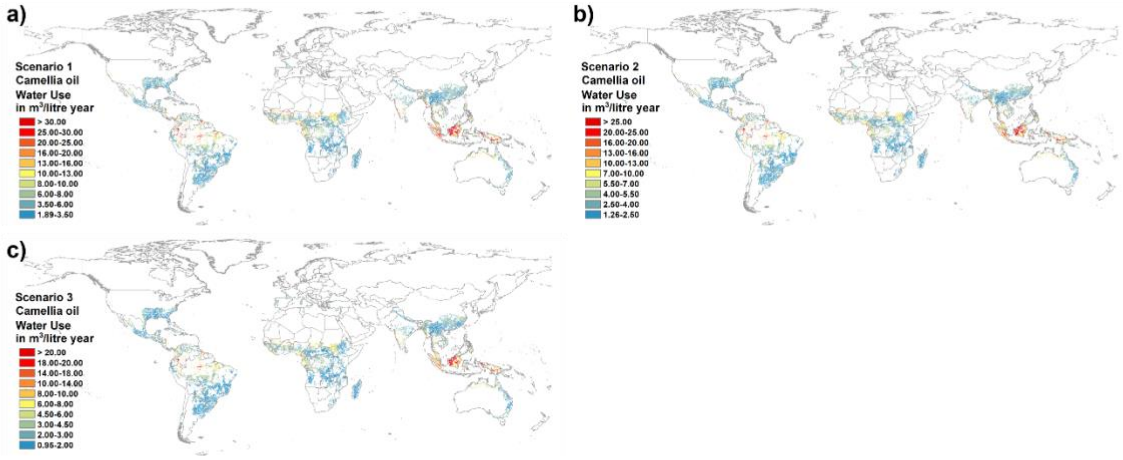


**Fig. S24.** Water use per liter *Camellia oleifera* oil: a) conservative yield scenario (S1; 50% potential yield); b) strategic yield improvement scenario (S2; 75% potential yield); and c) technology-policy-enabling yield scenario (S3; 100% potential yield)

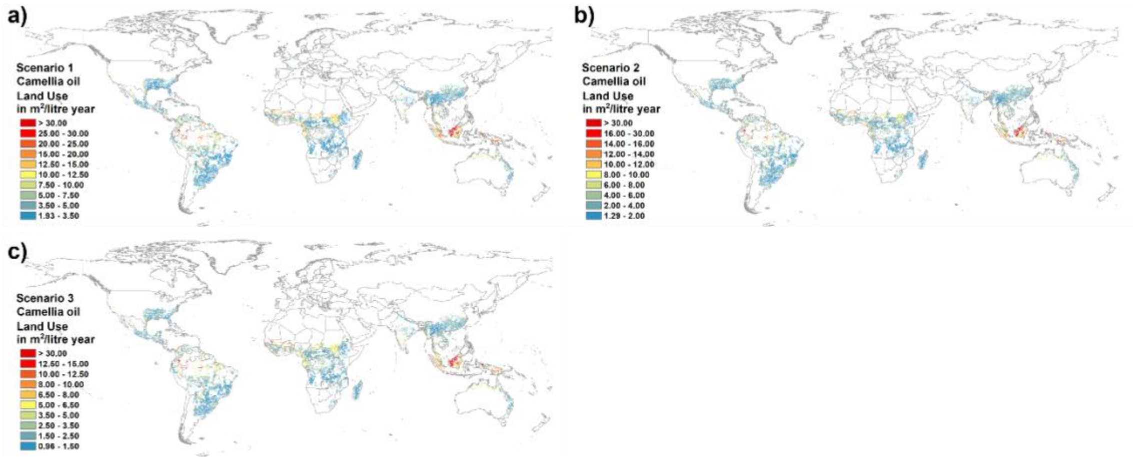


**Fig. S25.** Land use per liter *Camellia oleifera* oil: a) conservative yield scenario (S1; 50% potential yield); b) strategic yield improvement scenario (S2; 75% potential yield); and c) technology-policy-enabling yield scenario (S3; 100% potential yield)

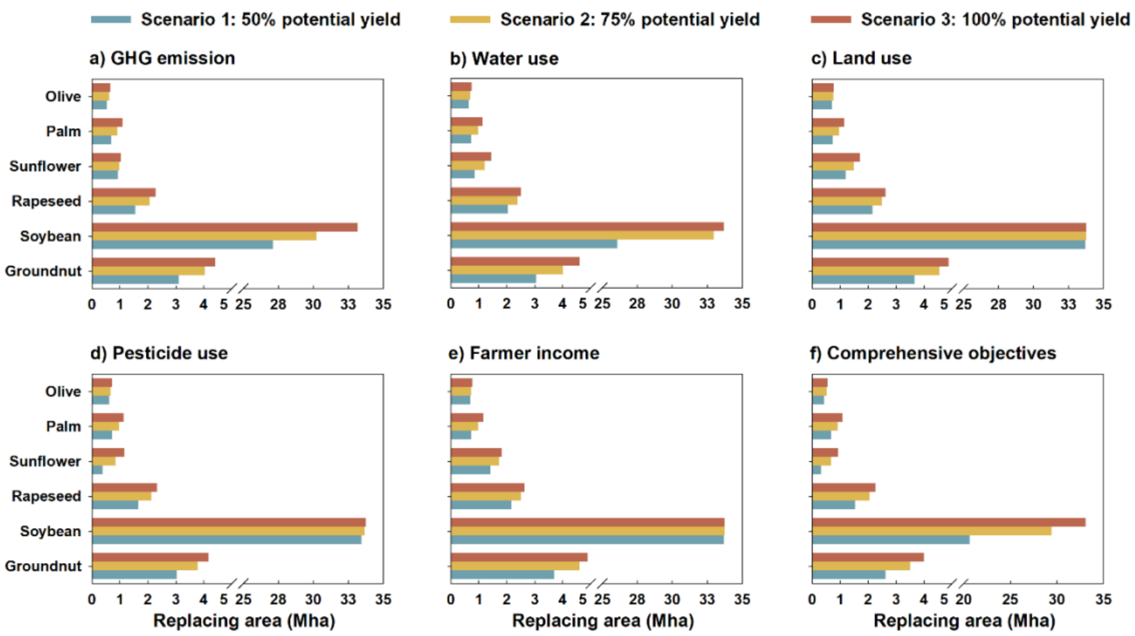


**Fig. S26.** The replacing areas of six vegetable oil crops under Camellia yields scenarios 1-3 for the five environmental improvement objectives individually and together.

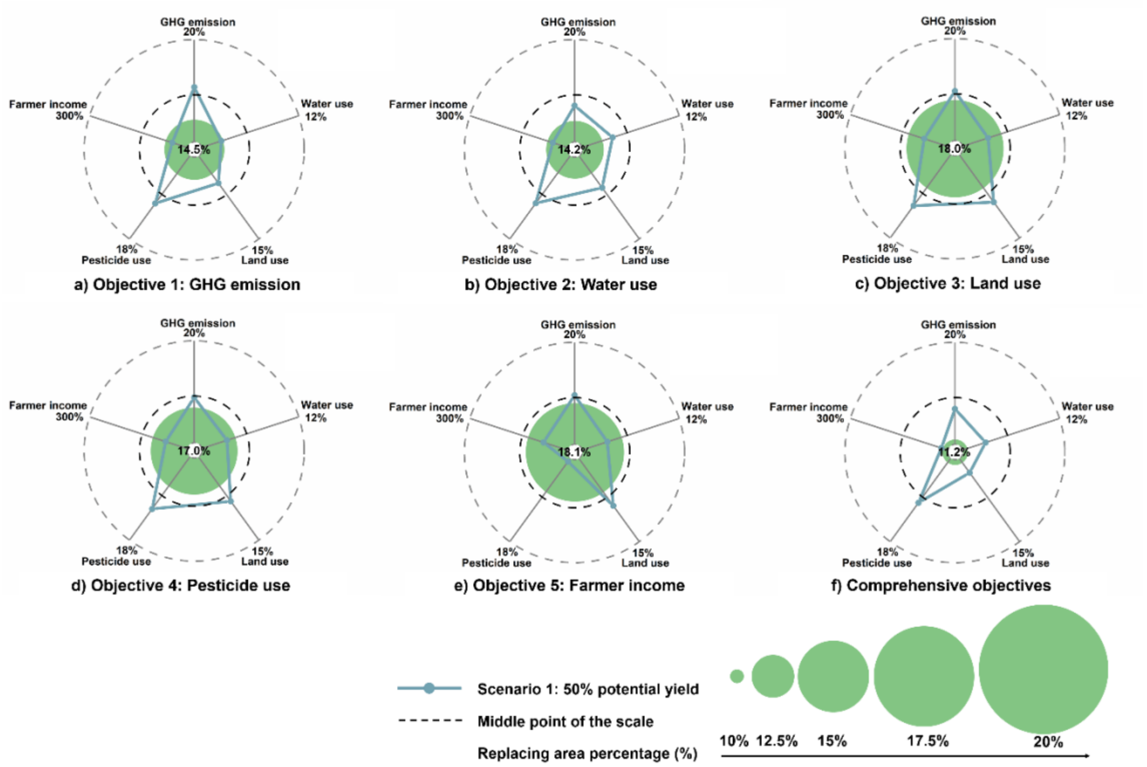


**Fig. S27.** Outcomes for *Camellia oleifera* under a conservative yield scenario (S1; 50% potential yield) to replace local vegetable oil crops according to different objectives: a) GHG emission; b) water use; c) land use; d) pesticide use; e) farmer income; and f) the comprehensive consideration of all. The percentages in the radar plot refer to the proportion of all five targets reached. The green circle is the total percentage of substituting global vegetable oil planting areas.

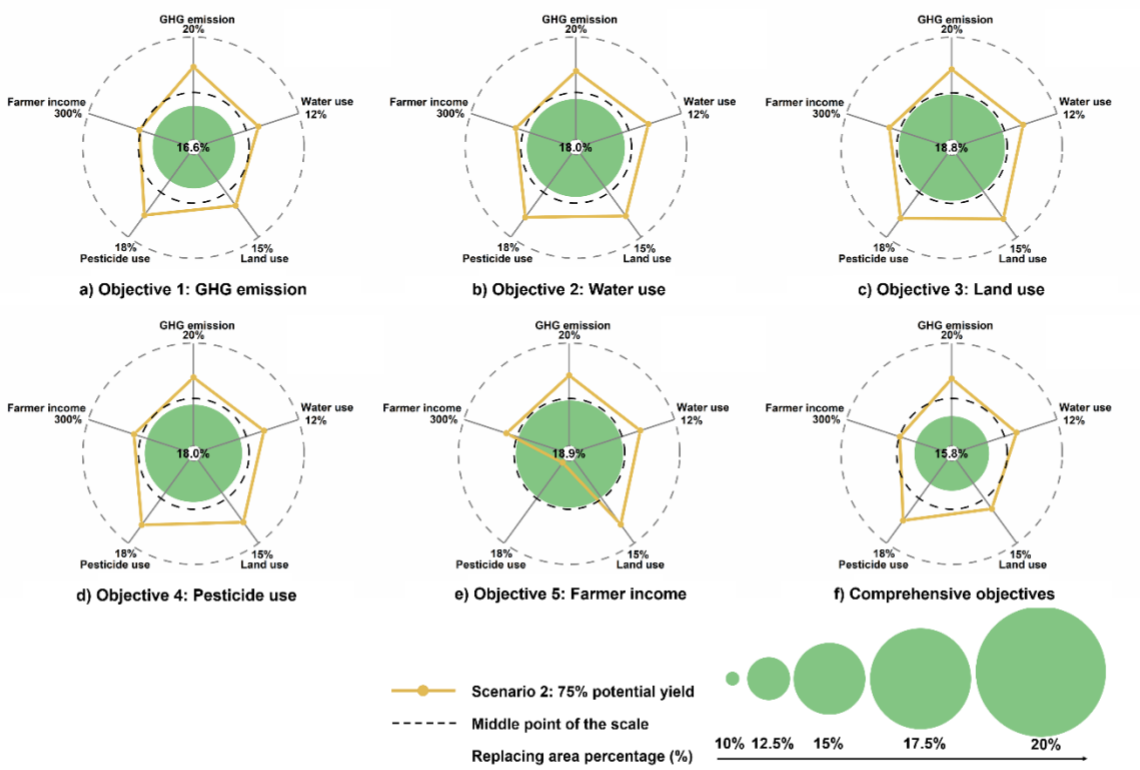


**Fig. S28.** Outcomes for *Camellia oleifera* under a strategic yield improvement scenario (S2; 75% potential yield) to replace local vegetable oil crops according to different objectives: a) GHG emission; b) water use; c) land use; d) pesticide use; e) farmer income; and f) the comprehensive consideration of all. The percentages in the radar plot refer to the proportion of all five targets reached. The green circle is the total percentage of substituting global vegetable oil planting areas.

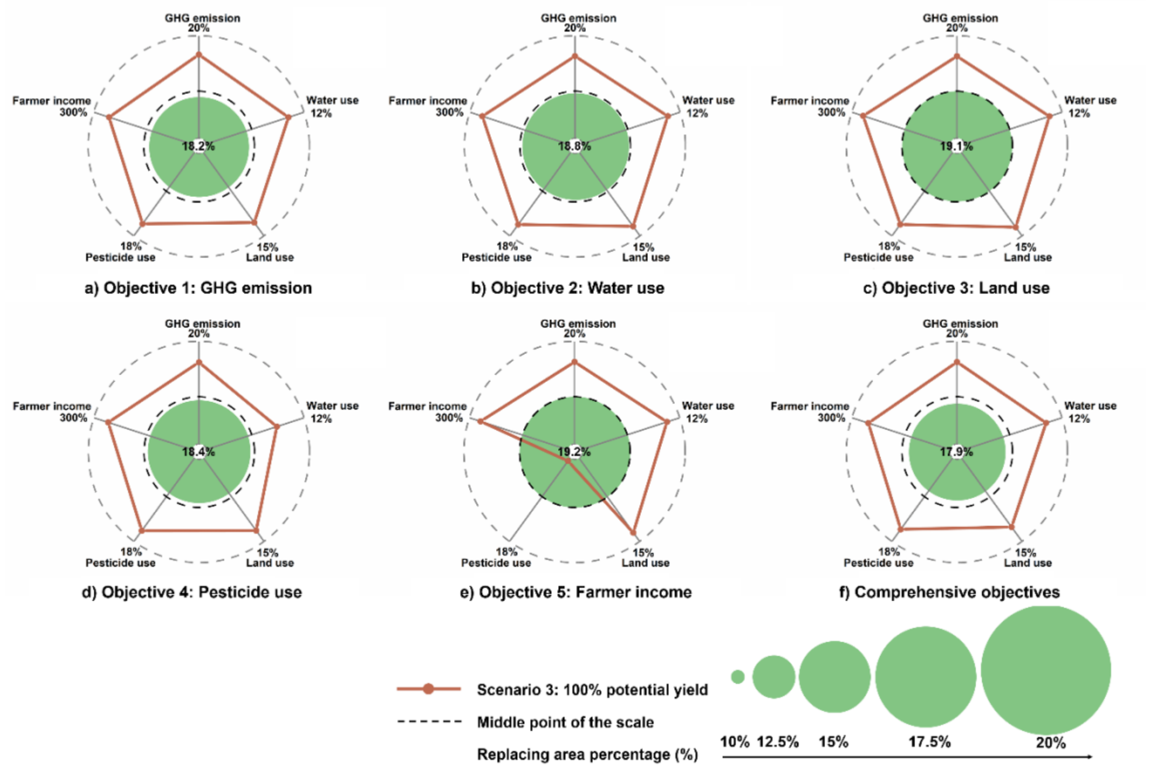


**Fig. S29.** Outcomes for *Camellia oleifera* under a technology-policy-enabling scenario (S3; 100% potential yield) to replace local vegetable oil crops according to different objectives: a) GHG emission; b) water use; c) land use; d) pesticide use; e) farmer income; and f) the comprehensive consideration of all. The percentages in the radar plot refer to the proportion of all five targets reached. The green circle is the total percentage of substituting global vegetable oil planting areas.

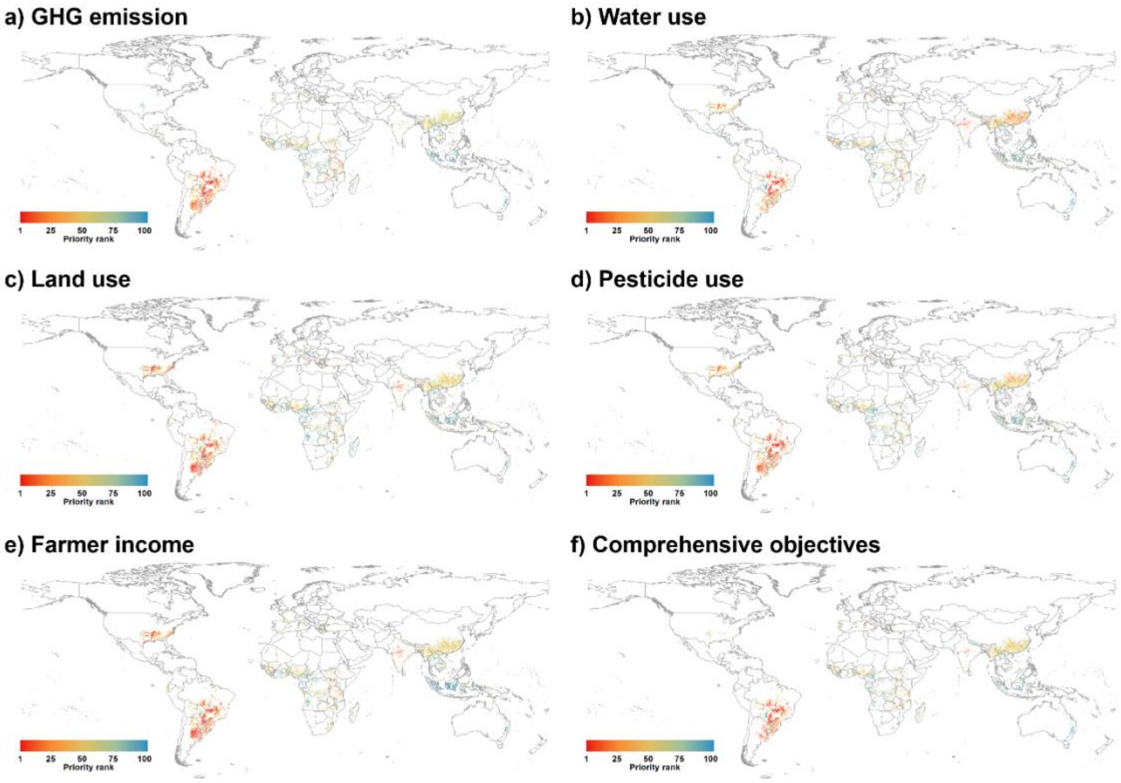


**Fig. S30.** Global priorities for *Camellia oleifera* under a conservative yield scenario (S1; 50% potential yield) to replace local vegetable oil crops according to five objectives: a) GHG emission; b) water use; c) land use; d) pesticide use; e) farmer income; and f) the comprehensive consideration of all objectives.

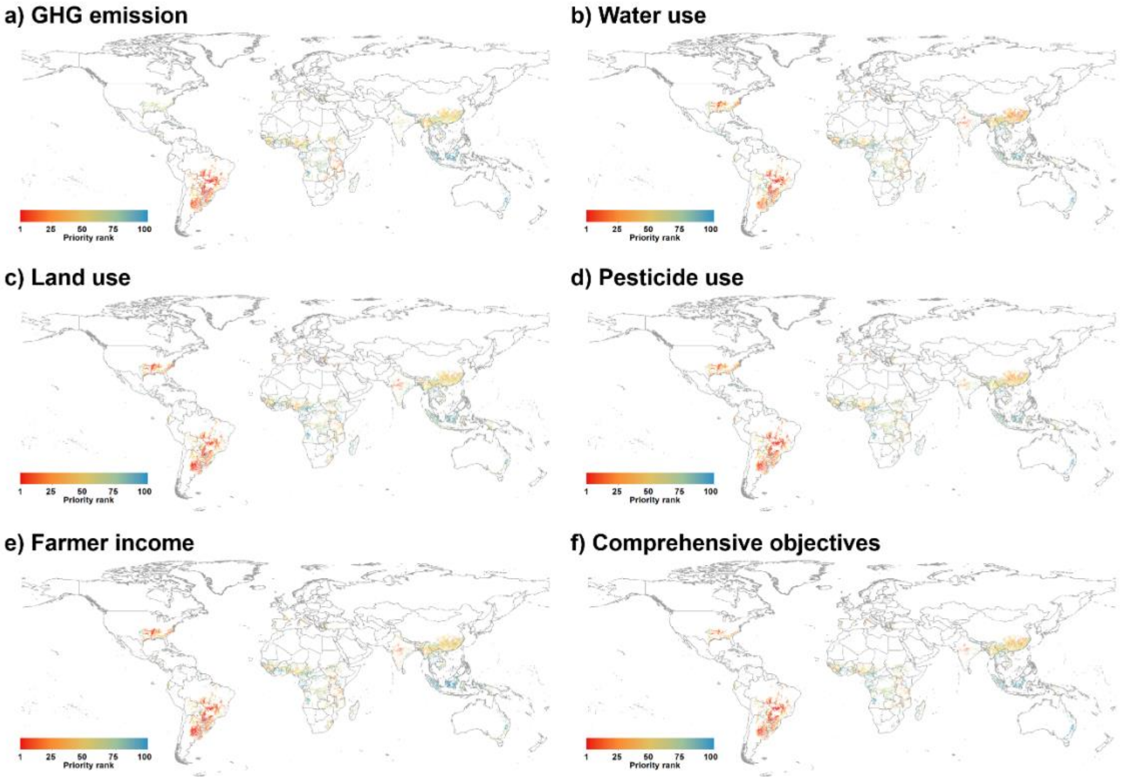


**Fig. S31.** Global priorities for *Camellia oleifera* under a strategic yield improvement scenario (S2; 75% potential yield) to replace local vegetable oil crops according to five objectives: a) GHG emission; b) water use; c) land use; d) pesticide use; e) farmer income; and f) the comprehensive consideration of all objectives.

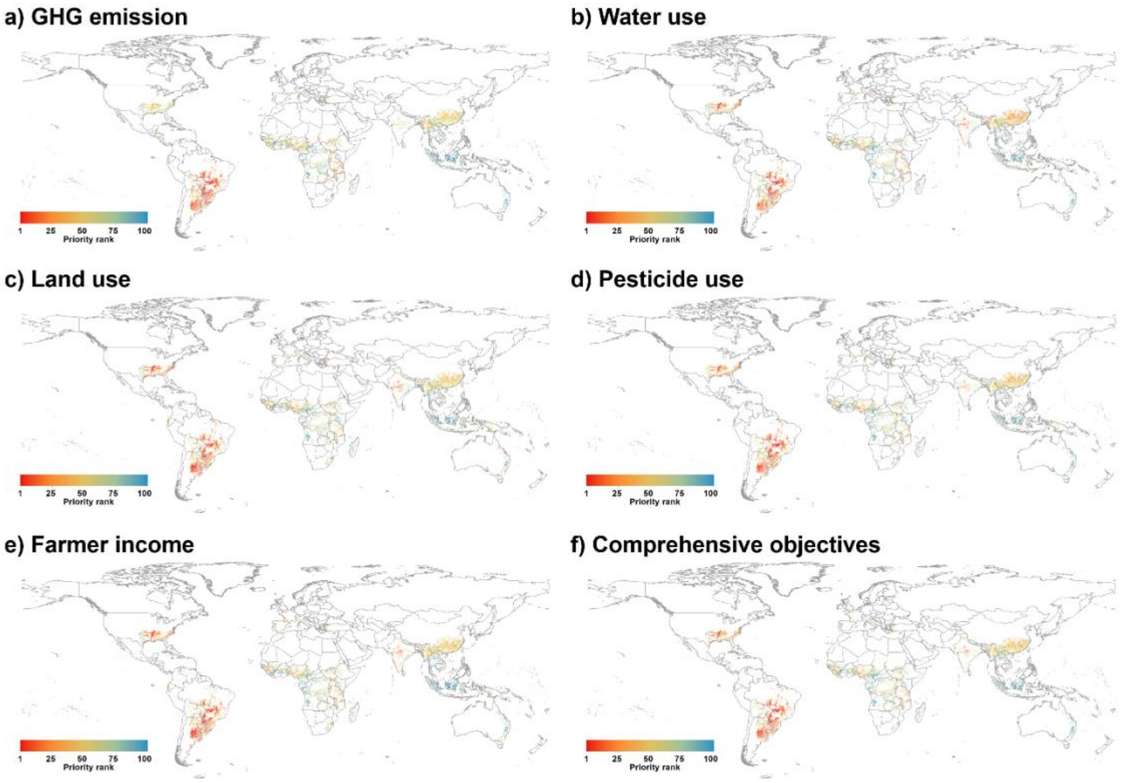


**Fig. S32.** Global priorities for *Camellia oleifera* under a technology-policy-enabling scenario (S3; 100% potential yield) to replace local vegetable oil crops according to five objectives: a) GHG emission; b) water use; c) land use; d) pesticide use; e) farmer income; and f) the comprehensive consideration of all objectives.

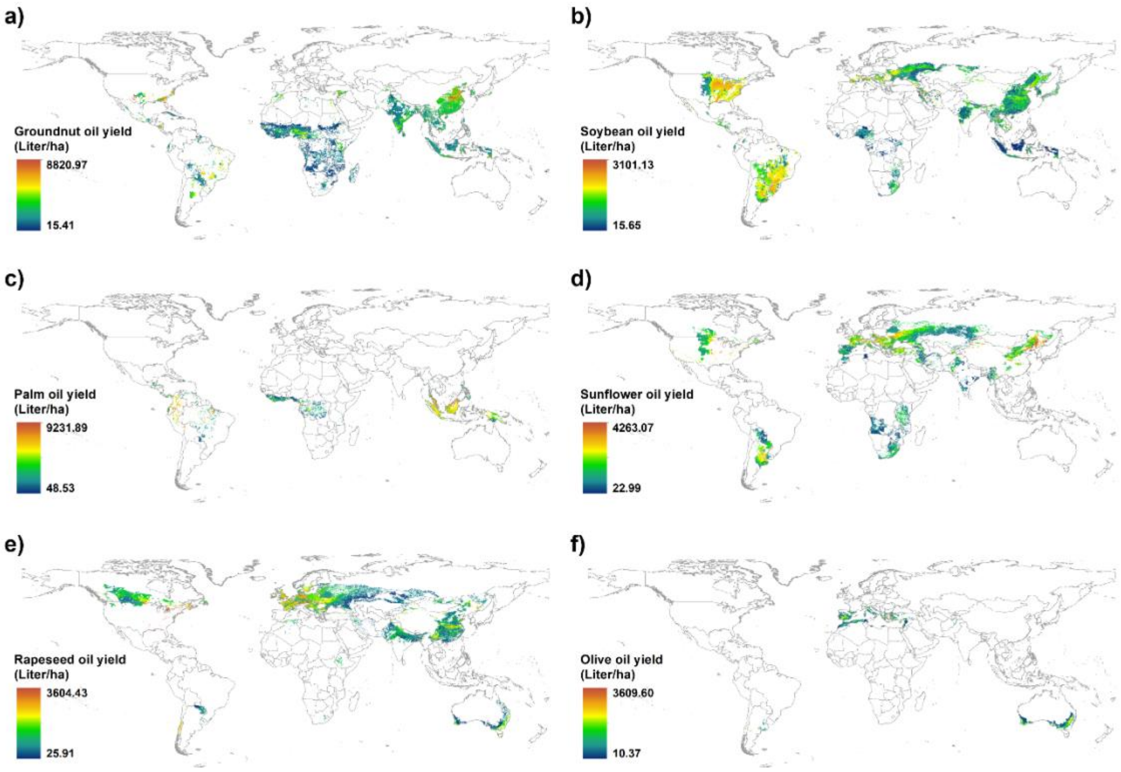


**Fig. S33.** The oil yield of all six vegetable oil crops: a) groundnut oil, b) soybean oil, c) palm oil, d) sunflower oil, e) rapeseed oil, and f) olive oil

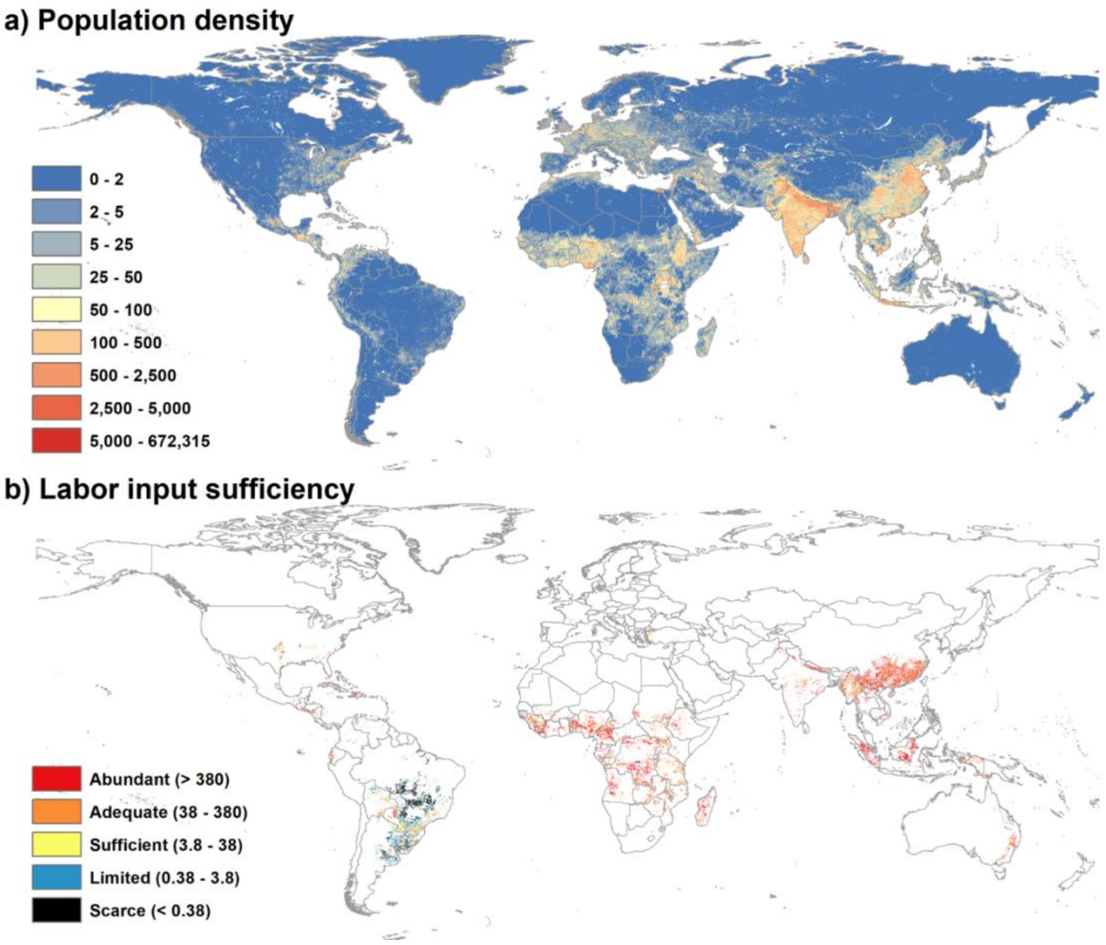


**Fig. S34.** Global population density with unit in number of people per $km^2$ (182) (a) and the five graded level of labor input sufficiency for *Camellia oleifera* under a conservative yield scenario (S1; 50% potential yield) to replace local vegetable oil crops according to the comprehensive consideration of all objectives (b). Note that *Camellia oleifera* cultivation is estimated need 0.38 worker per hectare harvested area.
